# Comparing arenavirus infection patterns in experimentally and naturally infected rodent hosts

**DOI:** 10.1101/111732

**Authors:** Joachim Mariën, Benny Borremans, Sophie Gryseels, Bram Vanden Broecke, Beate Becker-Ziaja, Rhodes Makundi, Apia Massawe, Jonas Reijniers, Herwig Leirs

## Abstract

Infectious diseases of wildlife are typically studied using data on antibody and pathogen presence/level. In order to interpret these data, it is necessary to know the course of antibodies and pathogen presence/levels after infection. Such data are typically collected using experimental infection studies in which host individuals are inoculated in the laboratory and sampled over an extended period, but because laboratory conditions are controlled and much less variable than natural conditions, the immune response and pathogen dynamics may differ. Here, we compared Morogoro arenavirus infection patterns between naturally and experimentally infected multimammate mice (*Mastomys natalensis*). Longitudinal samples were collected during three months of bi-weekly trapping in Morogoro, Tanzania, and antibody titer and viral RNA presence were determined to assess whether the natural temporal patterns are similar to those previously observed in the laboratory. A good match with laboratory data was found for 52% of naturally infected individuals, while most of the mismatches can be explained by the presence of chronically infected individuals (35%), maternal antibodies (10%) and an antibody detection limit (25%). These results suggest that while laboratory data are useful for interpreting field samples, there can still be differences due to conditions that were not tested in the laboratory.

**Important notice:** This is a pre-print version of the manuscript, made available through bioRxiv.org. Note that this manuscript has not yet been peer-reviewed, and has been submitted to a peer-reviewed journal.

## Introduction

Understanding disease transmission in wildlife populations relies on a correct interpretation of infection data. When data are difficult to measure in natural conditions, e.g. length of the infectious period, immune response duration and dynamics, or transmission mechanisms (vertical, horizontal, sexual, etc.), challenge experiments are typically used, where healthy animals are inoculated under controlled laboratory conditions (Fulhorst et al., 2002; Botten et al., 2000; Hardestam et al., 2008). The validity of such experimental data is conditional on the assumption that infection patterns in natural conditions are the same as those in a controlled laboratory setting. Dissimilarities between natural and laboratory infections can result from naturally occurring variance in many factors, including stress, presence of different genetic strains, inoculum volume, transmission route (e.g. nasal, gastro-intestinal, sexual) and individual life history (e.g. number of pregnancies, reproductive status, past infection) (Childs & Peters 1993; Martin et al., 2008; Billings et al., 2010a; Mills et al., 2010).

Comparing data from animals sampled in the wild to data from laboratory animals can however be difficult. An important reason for this is the need to know exactly when a sampled individual became infected in the field in order to interpret the consequent antibody or parasite kinetics, or to determine the infectious and immune period (Voutilainen et al., 2015; Begon et al., 2009). Determining the exact time of infection (TOI) in natural populations is most often unfeasible because usually not all individuals can be captured repeatedly, as a large effort is required to collect sufficient long-term data from wild animals, and because the sampling frequency (typically 4 weeks or more) is often too low for quantifying infection patterns (Pollock et al., 1990; Tersago et al., 2012; Telfer et al., 2002; Abbott et al., 1999; Sluydts et al., 2007; Cooch et al., 2012). While it is therefore not surprising that few studies have performed a cross-validation of infection data from laboratory and natural conditions, there is nevertheless a great need for such comparisons.

The aim of this study is to validate the dynamics of Morogoro virus (MORV), an African arenavirus in blood and excreta of its natural rodent host *Mastomys natalensis,* and the dynamics of the anti-MORV antibody response. A recent experimental infection study in controlled laboratory conditions revealed that immunoglobulin G levels as well as the presence of viral RNA (vRNA) exhibit a predictable temporal pattern post-inoculation, with relatively little variation between individuals (Borremans et al., 2015). As a result, it was possible to use data on antibody and/or vRNA levels from naturally infected individuals to estimate when an individual became infected in the field (Borremans et al., 2016). However, such time-since-infection estimates, when used on data collected in natural conditions, are based on the assumption that naturally infected animals exhibit antibody and virus dynamics that are sufficiently similar to those of laboratory animals.

Here, we compared temporal variations in virus shedding (vRNA presence/absence in blood and excretions) and host immune response (antibody titer dynamics) between experimentally and naturally infected rodents. By focusing our efforts on a brief but high-intensity sampling effort, we were able to obtain a dataset of sufficiently high temporal resolution to allow us to quantify individual infection dynamics of naturally infected rodents, and compare these qualitatively with temporal patterns in experimental data.

## Methods

### Model system

Morogoro virus (MORV) is an East African arenavirus closely related to Mopeia virus and Lassa virus, the latter of which causes severe haemorrhagic Lassa fever in humans in West Africa (Günther, et al., 2009; Wulff et al., 1977). These viruses all occur in the same host species, the Natal multimammate mouse (*Mastomys natalensis*), one of the most common rodents in sub-Saharan Africa and a known agricultural pest and carrier of pathogens (Mwanjabe et al., 2002; Katakweba et al., 2012). Due to their similarities and the fact that they are not pathogenic, MORV and Mopeia virus are considered to be safe alternatives for research on Lassa virus ecology and vaccine development (Lukashevich et al., 1999; Rieger et al., 2013; Borremans et al., 2011). In a recent challenge experiment under laboratory conditions, we determined the antibody (Ab) response and viral shedding patterns after intra-peritoneal inoculation of MORV in *M. natalensis* (Borremans et al., 2015). One day after inoculation, viraemia starts and continues for a period of 15 days after which the virus disappears in blood, while MORV RNA remains detectable in urine, saliva and faeces until about 40 days after inoculation. Antibodies (IgG) against MORV are detectable from day 7 post infection (pi), and follow a clear, predictable pattern characterized by a high initial increase phase peaking around day 20 pi, followed by a decrease phase that reaches a minimum approximately 70 days pi, after which Ab titers again start to increase until reaching a final equilibrium concentration from day 160 pi onwards (Borremans et al., 2015). The mice used in this challenge study were part of a breeding colony initiated in 1999 using mice from Morogoro (Tanzania), which is the same location where we collected field samples for the current comparative study.

### Field and molecular work

Between 30 July 2013 and 18 October 2013, a rodent capture-mark-recapture (CMR) study was performed in five grids of 100m by 100m in Morogoro, Tanzania. Blood and saliva samples were taken from each animal and preserved on filter paper (Borremans 2014). Animals were individually marked using toe clipping (Borremans et al. 2014), and weight, sex, and reproductive state were recorded. All individuals that were captured during at least two different trapping sessions were analysed for the presence of anti-MORV antibodies (Abs) by indirect immunofluorescence assay and viral RNA (vRNA) by RT-PCR. Antibody titers were estimated using two-fold dilution series, starting with a minimum dilution of 1:10. A more detailed explanation of the field and laboratory work can be found in the supplementary information (supplementary text 1 and 2).

### Comparing field and laboratory data

In order to compare the Ab and vRNA patterns of naturally infected mice with those obtained through experimental infection (Borremans et al., 2015), we needed to simultaneously plot both datasets on a figure showing Ab titer and vRNA presence against time after infection, and in order to do this it was necessary to know the exact time of infection (TOI). While the exact TOI is known for the experimentally infected mice, this is obviously not the case for those that were infected naturally. As the best possible alternative, we used the TOI that resulted in the best possible match between the measured data of a wild mouse and the laboratory data. If a good match between laboratory and natural infection patterns can be found, this would be a strong indication that experimentally acquired data are representative for natural infections; it is impossible to statistically test the match unless the exact TOI in natural conditions is known.

To estimate the TOI of naturally infected mice, we used the method described in Borremans et al., 2016. Briefly, this method integrates existing data on Ab presence/absence and titer, vRNA presence in blood and excretions (saliva and/or urine), and body weight into one semiparametric, Bayesian model that can be used to estimate the most likely TOI given the available information. The method takes into account all available data about a captured individual, which generally means that the error on the estimated TOI will be smaller for individuals that were re-captured more often. Here, the infection data used to inform the TOI estimation model originates from the experimental data described above, which means that the use of this method will result in the best possible match between the experimental and natural patterns. Therefore, it is important to bear in mind that this method is potentially positively biased towards finding a good match between temporal patterns, does not allow obtaining statistical proof for the matches and will only allow the identification of obvious discrepancies between experimental and natural temporal infection patterns.

Because it may be possible that discrepancies between temporal patterns of laboratory and field data only occur in either Ab or vRNA data and not necessarily in both simultaneously, the TOI was estimated using two scenarios. In the first (method 1), only Ab titer data were used, in order to test whether Ab titer patterns are similar regardless of the vRNA patterns. In the second scenario (method 2), both vRNA data from blood and excretions and Ab data were incorporated. Additionally, to slightly improve estimation of the TOI, all scenarios take into account body weight by limiting the maximum age of animals with a body weight below 20g (= juveniles) to 120 days following (Leirs 1994), which means that they could not have been infected more than 120 days before the date of capture. We also considered a third scenario where only vRNA data were used to estimate the TOI, but because this method lacks validity (see below) the results of this method were not further discussed in this article.

Only mice that were recaptured during multiple trapping sessions and were Ab-positive during at least one of these sessions were retained for TOI estimation. Thus, mice that were recaptured and were vRNA-positive at least once but never became Ab-positive were not included because we assume they became infected at a young age after which they developed Abs at titers under the detection threshold of the used immunofluorescence assay (Begon et al., 2009), which is a condition that was not investigated in the laboratory before. After the TOI estimations, field and experimental data were plotted together to allow a qualitative assessment of the match between infection patterns. R statistical software (R Core Team 2014) was used for data manipulation, TOI estimation and plotting.

We defined a good match for a tested individual if all its Ab titers and/or vRNA presence/absence data fell within the confidence band (CB) of the laboratory data. More detailed information about this calculation can be found in the supplementary information (supplementary information text 3).

### Testing the validity of the different TOI methods

Because the TOI method is biased towards finding a good match between field and laboratory data, we also tested whether the obtained results are statistically different from those resulting from random data, where observation data (Ab, vRNA) are randomized for each individual, i.e., when the presumed temporal pattern (which contains information on the TOI) was destroyed. If this was not the case, this would be an indication that: (1) there is insufficient information to estimate TOI sufficiently accurate to make a meaningful comparison between field and laboratory data (e.g. because animals were not recaptured frequently enough in this study); and/or (2) the observed temporal dynamics in the field data differ considerably from those measured in the laboratory. We assessed this using a permutation test. For each permutation, we randomly permuted for each individual the Ab titers and/or vRNA presence in blood and excretions and then calculated the percentage of individuals that matched the laboratory data. We considered 10,000 permutations of the field data.

## Results

During 10,800 trap nights (the number of traps times the number of trapping nights), we captured 1,133 *M. natalensis* individuals of which 220 were recaptured at least once during a different trapping session. All samples of recaptured individuals (590 samples) were analysed for both Abs and MORV vRNA (Table 1).

**Table 1:** Number of *M. natalensis* recaptured during different trapping sessions.

| Times captured | Total | Number of individuals |  |  |  |
| --- | --- | --- | --- | --- | --- |
|  |  | Antibody + MORV RNA neg | Antibody pos | MORV RNA pos | Antibody + MORV RNA pos |
| 2 | 149 | 104 | 15 | 17 | 14 |
| 3 | 39 | 30 | 3 | 2 | 4 |
| 4 | 27 | 20 | 1 | 1 | 5 |
| 5 | 3 | 2 | 0 | 1 | 0 |
| 6 | 2 | 0 | 0 | 0 | 0 |
| 7 | 1 | 1 | 0 | 0 | 0 |
| **Total** | **221** | **157** | **19** | **21** | **23** |

<sup>a</sup> Ab-positive during at least one trapping session, but vRNA-negative
<sup>b</sup> vRNA-positive during at least one trapping session, but Ab-negative
<sup>c</sup> Ab and vRNA-positive during at least one trapping session

We found that 10% (21/220) of rodents were vRNA-positive at least once and never Ab-positive, 9% (19/220) were at least Ab-positive once and never vRNA-positive, 10% (23/220) were vRNA- and Ab-positive at least once at the same or a later moment of which 78% (18/23) were simultaneously vRNA- and Ab-positive at least once (Table 1). From the 21 vRNA-positive mice that were never found to be Ab-positive, 13 mice were vRNA-positive during one capture session but not during later recaptures (eight times two weeks, four times four weeks and once eight weeks later); two individuals were vRNA-positive during two consecutive recaptures (one x two weeks and one x four weeks later); and six individuals were positive during their last recapture. This means that 26% (15/57) of infected mice showed no signs of an active Ab response two weeks or more after infection. The six mice that were vRNA-positive at their last recapture were not included in this sum because we cannot determine whether they were sampled before they could produce Abs (day six after inoculation in laboratory conditions).

For the TOI analyses, we used only the 42 individuals that were Ab-positive at least once. When TOI was estimated based on Ab levels only (method 1), the temporal patterns of natural and experimental Ab dynamics were remarkably similar (i.e. there were very few instances of a bad match). We found that 88% (37/42) of individuals matched the laboratory immune response, of which 91% (82/90) of all the collected field data observations fell within the 95% confidence band (CB) of the laboratory data (Table 2;Fig 1). Of the remaining ten data points, seven were Ab-negative while based on the laboratory data they would have been expected to be Ab-positive (Fig 1 and e.g. supplementaryFig S21).

**Table 2:** Matches between field and laboratory data of *M. natalensis* infected with MORV for the two different TOI methods

|  | Time of infection method |  |  |  |
| --- | --- | --- | --- | --- |
|  | 1. Antibody | 2. Antibody + MORV vRNA |  |  |
|  | Ab titers | Ab titers | Blood | Excretions |
| **Individuals** | 88% (37/42) | 67% (28/42) | 71% (27/38) | 87% (34/39) |
| **Observations** | 91% (82/90) | 78% (73/94) | 85% (69/81) | 93% (76/82) |

For both TOI estimation methods the observed percentages of matches between field and laboratory data are given on the individual level and total number of observations: method 1 is based on Ab titers only and method 2 on both Ab titres and vRNA presence in blood and excretions.

**Fig 1.**
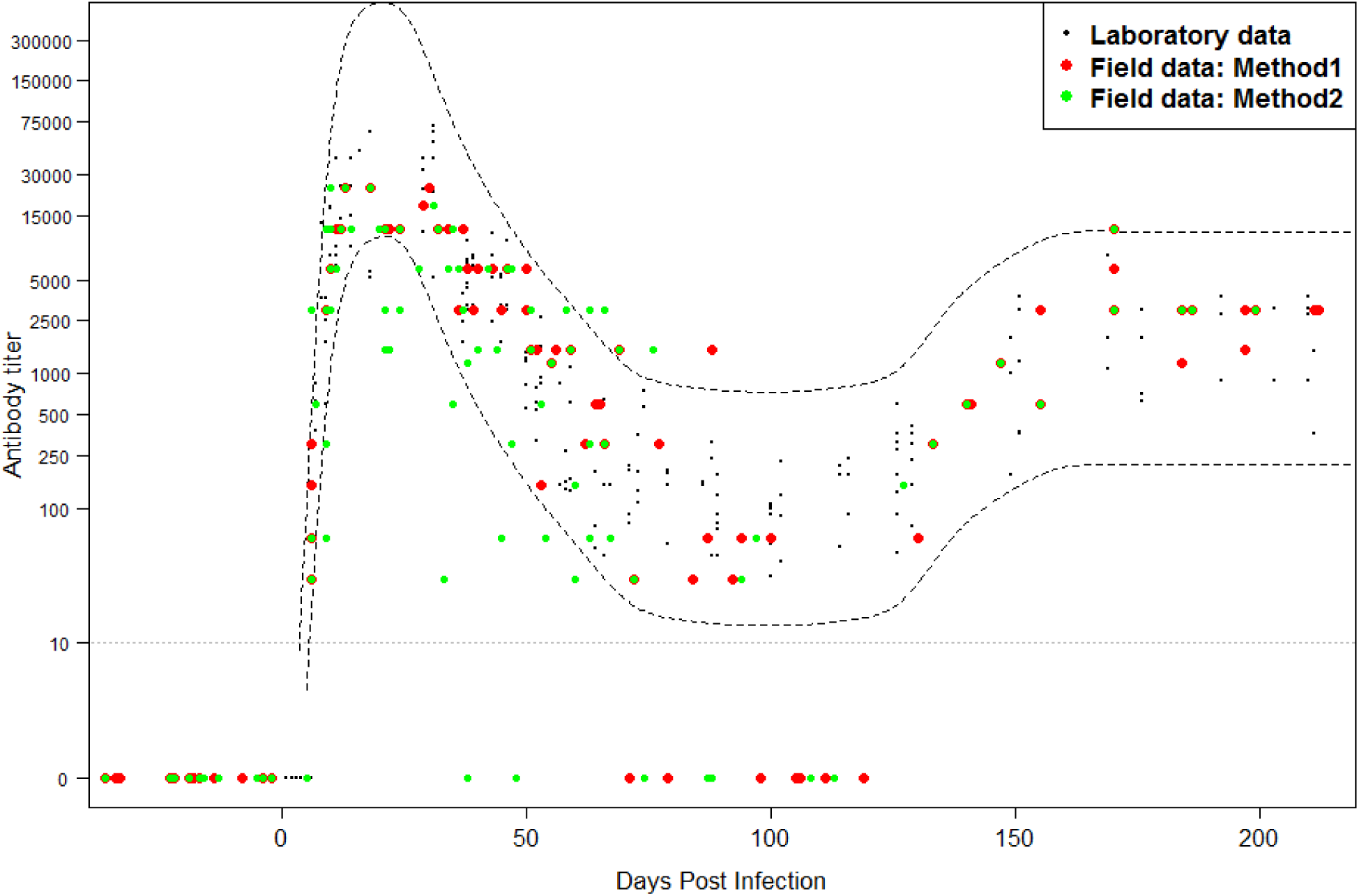
Temporal Ab dynamics of naturally and experimentally MORV-infected *M. natalensis.* Small black dots represent Ab levels from experimentally infected rodents in the laboratory (data derived from (Borremans et al., 2015)), red and green dots represent Ab titers from naturally infected rodents observed in this study. The TOI of the red field data (dots) was estimated using the TOI estimation method based on Ab titer only (method 1). The TOI of the green field data (dots) was estimated using the TOI estimation method based on Ab titer and vRNA (method 2). Ab patterns of 42 naturally infected individuals were plotted together in this graph.

When both Ab levels and presence of vRNA were taken into account (method 2), vRNA and Ab level patterns were still roughly in agreement with laboratory results. For the Ab response, we found that 67% (28/42) of individuals matched the laboratory results, of which 78% (73/94) of all the collected field data fell within the 95% CB of the laboratory data (Table 2; Fig 1). We found that 71% (27/38) of individuals matched the vRNA dynamics of the laboratory data in blood and 87% (34/39) in excretions, of which 85% (69/81) and 93% (76/82) of all the collected field data fitted the predicted probabilities respectively (Fig 2). For the combined results of temporal Ab and vRNA patterns, we found a good match with the laboratory patterns in 52% (22/42) of naturally infected mice (supplementaryTable 2). The mismatches were due to the prolonged presence of vRNA in blood or excretions (e.g. supplementary Fig S6), the absence of Abs at times that they would be expected to be positive (Fig S21), the potential presence of maternal Abs (Fig S29), and overall lower Ab titers in naturally infected animals (Fig S1).

**Fig 2.**
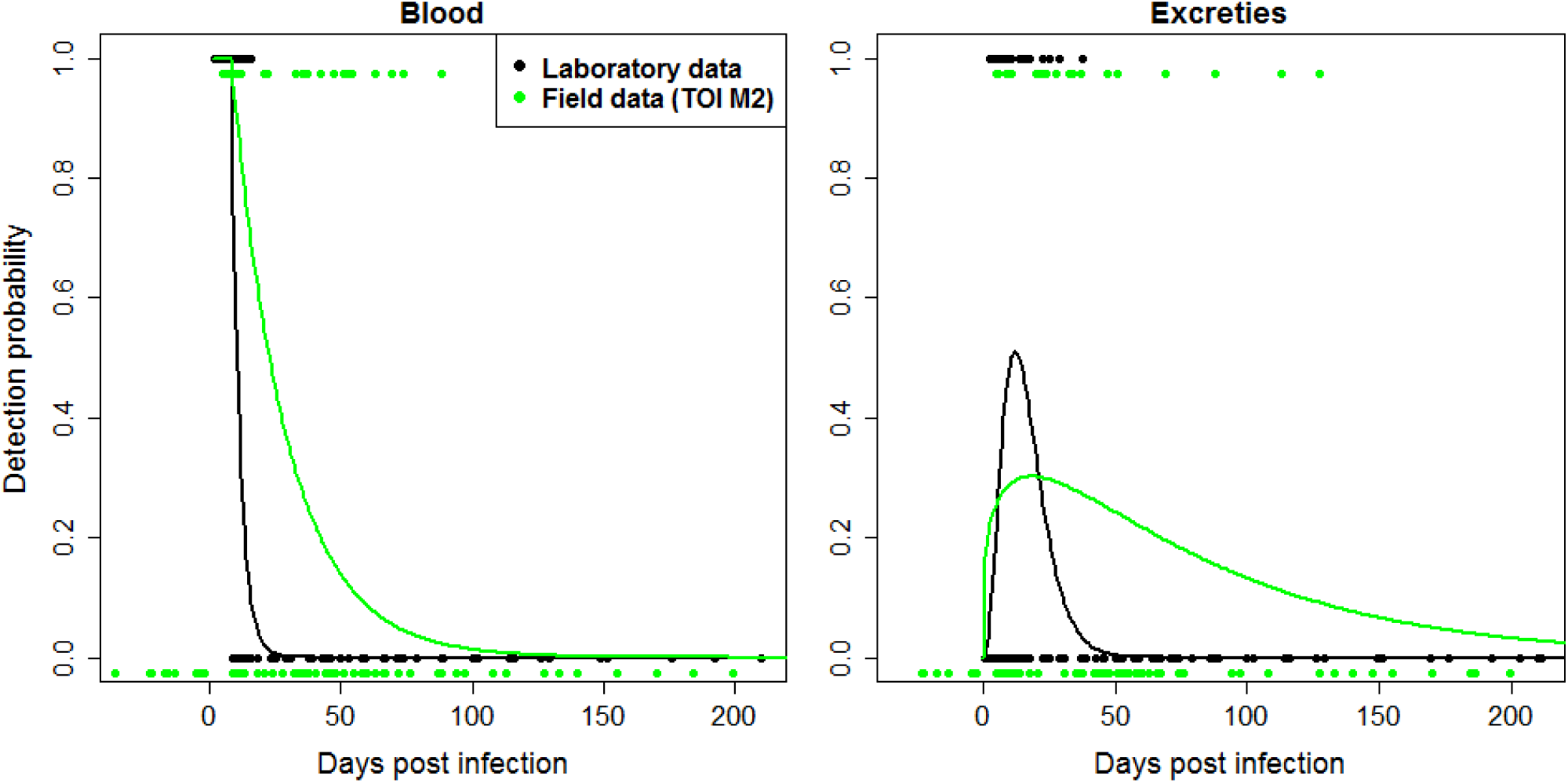
Temporal detection probability of MORV vRNA presence in blood (left) and excretions (right). The black points/curve represent raw data (points) and the proportion of samples (curve) that were vRNA-positive in laboratory conditions (i.e. vRNA presence probability; data derived from [11]). The green points/curve represent raw data (points) and the proportion of samples (curve) that were vRNA-positive in field conditions, and were estimated using the TOI estimation method based on all available data (method 2, Ab titer, vRNA presence in blood and excretions, weight cutoff). vRNA patterns of 42 naturally infected individuals were plotted together in this graph.

### Testing validity of TOI methods

When TOI was estimated based on Ab titers alone (method 1) or on both Ab titers and vRNA in blood and excretions (method 2), real field data matched laboratory data significantly better than random permutations (p-value < 0.0005 for both methods) (Fig 3). In contrast, if TOI was estimated based on vRNA observations alone (method 3), real field data did not match laboratory data better than permutations (p-value = 0.9291) (Fig 3). We conclude that methods 1 and 2 are appropriate to make a meaningful comparison between field and laboratory data, but that it is not possible to make such a comparison based on vRNA data alone (method 3) because good matches were much too likely. Results of method 3 were therefore not further discussed in this paper.

**Fig 3.**
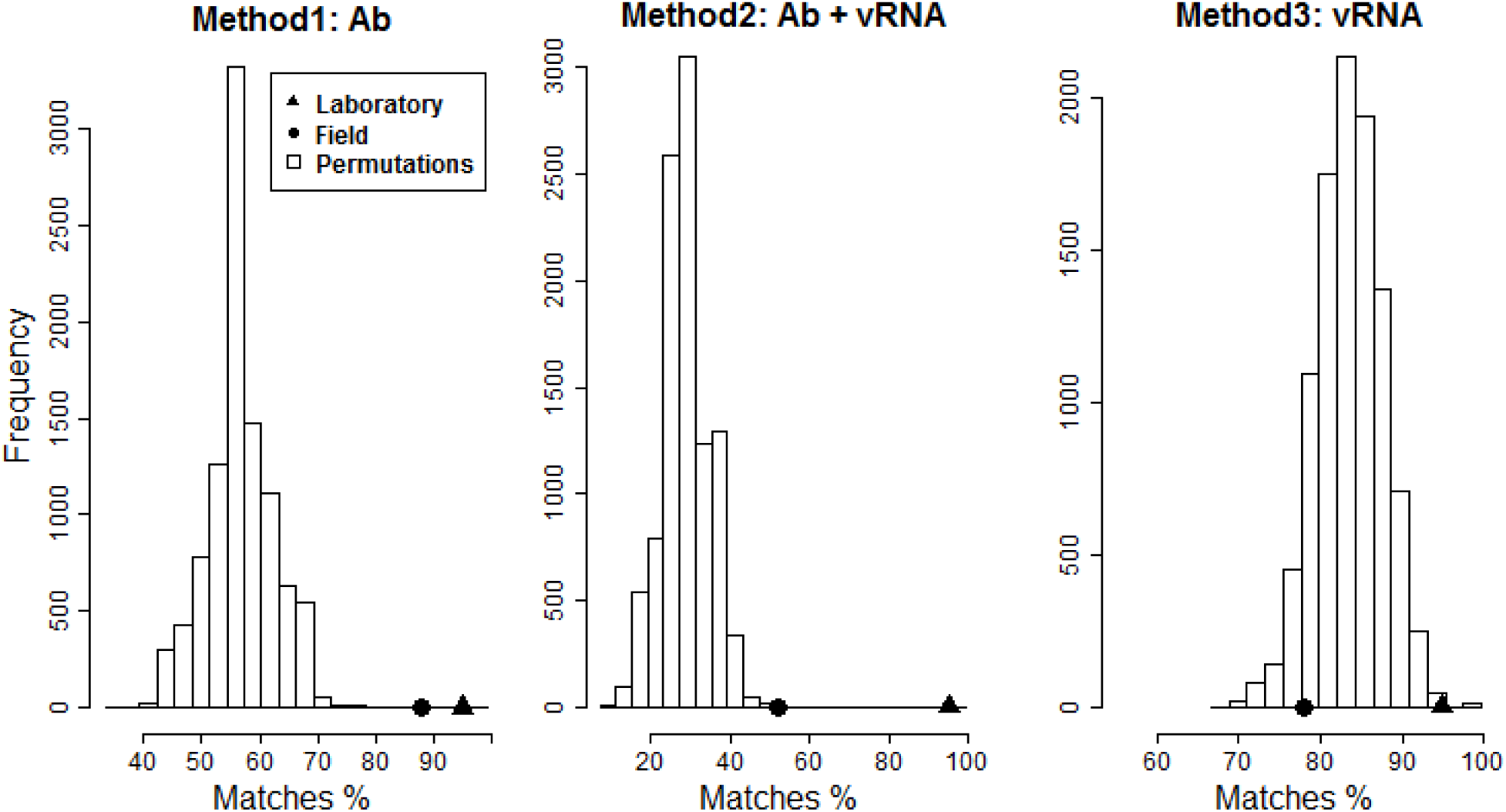
Percentages of individuals for which the infection patterns in viral shedding and/or host immune response matched the laboratory data. Random permutations of field data (bars) were compared with real field data (dots) and laboratory data (triangles, by default 95%) for the three different TOI methods: Ab titers (method 1), Ab titers and vRNA in blood and excretions (method 2), and vRNA in blood and excretions (method 3).

## Discussion

We found that 74% of infected individuals in the field developed an Ab response within at least two weeks after the infection was detected. When both Ab and vRNA data of these Ab-positive individuals were taken into account, we found a good match between field and laboratory infection patterns for 52% of individuals. This percentage is significantly higher than in case the Ab and/or vRNA observations are randomized for every individual, but is clearly lower than the 95% matching percentage that is to be expected in the laboratory. This lower percentage could be due to several mechanisms, which are discussed below.

### MORV vRNA-positive without antibody response

First, we found that 26% of mice developed no Ab response after MORV infection in the field (with animals tested at least two weeks after MORV infection was detected), in contrast to laboratory conditions where all inoculated mice seroconverted within 6 days. One possibility to explain this result is that these mice developed Abs at titers that fell below the detection threshold of the used immunofluorescence assay. Such lower Ab titers could be due to infection at a young age. Studies based on early serological methods thought that neonatal *M. musculus* infected with lymphocytic choriomeningitis arenavirus (LCMV) did not develop Abs and were assumed to be immuno-tolerant (Burnet & Fenner 1949; Weigand & Hotchin 1961). However, when analysed later using more sensitive immune assays, Abs did appear to be present, but at low titers (Oldstone & Dixon 1967; Oldstone 2002). This may also be the case for neonatally infected *M. natalensis* in nature, although we have previously observed that laboratory-infected neonatals develop a chronic MORV infection with the presence of Abs at normal titres (Borremans et al., 2015).

Another hypothesis would be that these mice indeed never produced Abs, but that the virus was cleared by the cell-mediated immune response alone. Although this response is not yet investigated for MORV infections in *M. natalensis,* it is known that cytotoxic T-cells can indeed clear LCMV infections without the help of Abs in B cell-deficient mice (Matloubian et al., 1994; Asano & Ahmed 1996).

### Lower antibody titers

The majority of mice (74%) did develop a clear immune response in the field, although Ab titers were systematically lower than in the experimental data (Fig 1, Fig 3). This was most obvious in the supplementary Fig S1 (individual 354060F1), Fig S2 (364060F1) and Fig S33 (760F6) where the Ab titers just fell outside the CB of the laboratory data. It may also have been the case for individuals shown in Fig S12 (390F3), Fig S21 (260F5), Fig S22 (370F5), Fig S29 (210F6) and Fig S37 (3100F6), where Ab titers of positive samples were generally low and where the laboratory results would be matched perfectly if the negative samples were in fact low, undetectable titers. Such an apparent loss of detectable Abs was also observed for a small percentage (<5%) of *Microtus agrestis* naturally infected with cowpox virus when analysed by indirect immunofluorescence assay (Begon et al., 2009; Chantrey 1999). Because *M. agrestis* normally shows a long-term Ab production, these samples were considered to be false negatives. Combined, our results imply that Ab titer values should be increased slightly when they are used for detailed analyses such as TOI estimation (Borremans et al., 2016).

### Chronic infections

Evidence for chronic infection was found for a number of individuals (13%), and was most convincingly seen in Fig S26 (individual 38F6), where an individual is depicted for which all samples were positive for Ab and vRNA in blood and almost all samples in excretions. There is also some evidence for chronic infection during which virus presence in blood or excretions is not constant but intermittent, suggesting temporary flare-ups of excretion; examples are shown in Fig S6 (15F3a), Fig S10 (330F3), Fig S22 (370F5), Fig S27 (48F6), Fig S36 (2680F6) and Fig S37 (3100F6). A chronic infection could also be an explanation for two individuals (20100F4 and 355060F1) that remained Ab-negative but were vRNA-positive in blood and/or excretions during two consecutive capture sessions (respectively 2 and 4 weeks interval).

If these patterns are indeed the result of chronic infection, this means that viraemia is not always transient in field conditions, as opposed to laboratory conditions where vRNA is only detectable for a short period in infected adults (until day 15 in blood and day 40 in excretions). For hantaviruses, chronic infection usually seems to result in temporary viraemia, after which the virus retreats into certain organs and is shed at lower concentrations (Yanagihara et al., 1985; Voutilainen et al., 2015; Fulhorst et al., 2002), although chronic infection can also result in persistent viraemia in some individuals, which has been observed for Black Creek Canal hantavirus and for natural infection with Puumala hantaviruses (Billings et al., 2010b; Voutilainen et al., 2015). For arenaviruses, both transient and persistent viraemia have been observed in rodent hosts. Most arenaviruses establish a chronic infection when experimentally inoculated in newborn hosts, but are cleared from the rodent’s body quickly when inoculated in adults: less than two weeks in blood and less than one month in excreta. This age-at-infection effect for natural hosts has been experimentally observed for the Old World arenaviruses (LCMV in *M. musculus,* Lassa and Morogoro viruses in *M. natalensis* (Buchmeier et al., 1978; Borremans et al., 2015; Walker et al., 1975), and the New World arenaviruses (Tamiami virus in *Sigmodon hispidus,* Catarina virus in *Neotoma micropus* and Whitewater Arroyo virus in *Neotoma albigula* (Milazzo & Fulhorst 2012; Murphy et al., 1976; Fulhorst et al., 2001). Other new world arenaviruses (Machupo virus in *Calomys callosus,* Junin virus in *Calomys musculinus* and Guanarito virus in *Zygodontomys brevicauda)* establish a chronic infection in all inoculated newborns and in about half of the adults (Fulhorst et al., 1999; Vitullo et al., 1987; Webb et al., 1975). In contrast, Latino arenavirus causes acute or chronic infections in newborn and acute infections in adult *C. callosus* (Webb et al., 1975).

### Maternal antibodies

The presence of maternal Abs could explain one, and perhaps two of the 20 animals with different infection patterns than expected from the laboratory. Fig S29 (individual 210F6) shows a young individual (body weight at first capture was 19g) of which the first sample was Ab-positive and the second negative, while all samples were vRNA-negative. It could therefore be the case that this individual still had some maternal Abs at very low titers when it was first sampled, but then lost the Abs. Another aberrant pattern was observed for a young individual shown in Fig S10 (330F3) (body weight at first capture were 20g). Here, maternal Abs might explain the presence and rapid decrease of Ab titers, followed by an vRNA-positive blood and excretion sample. For the latter individual, initial Ab titers were as high as those observed after inoculation in the laboratory, which is unusual for maternal Abs although high maternal Ab titers have also been observed against Sin Nombre hantavirus in naturally infected *Peromyscus maniculatus* (Borucki et al., 2000) and for a number of different pathogens in *Microtus pennsylvanicus* (Glass et al., 1990).

### Recent infection without positive MORV vRNA sample

Finally, there is some indication that blood is not always vRNA-positive during the first week after infection. Five individuals [depicted in Fig S5 (3710100F1), Fig S9 (210F3), Fig S11(340F3), Fig S23 (630F5) and Fig S34 (880F6)] show an Ab-titer pattern that strongly suggests recent infection, albeit without an vRNA-positive blood sample shortly after infection. This pattern can however also be explained by Ab titers that temporarily (e.g. somewhere between the 100-120 day time interval) lie below the detection threshold and increase again two weeks later, as such mimicking the situation in the laboratory where Ab-titres show the highest-slope increase between day 6 and 20 after infection.

## Conclusion

The majority of MORV infection patterns observed in the field seems to fit the laboratory data, which means that more often than not it is possible to use laboratory patterns of MORV as a basis for the interpretation of field samples. Note that although we found a relatively good match between laboratory and field data, this does not equal proof that natural infection patterns are generally the same as those in the laboratory (it is possible to reject a statistical null hypothesis but not to prove it). What we did find in this study is that based on the observed similarities there is no evidence to reject the assumption that natural and laboratory infection patterns are similar. For the remaining cases where we did observe a mismatch between field and laboratory patterns, simple hypotheses (Ab detection threshold, chronic infection, and maternal Abs) exist that could explain the patterns, and should thus be considered when interpreting field samples. Overall, our results are encouraging, as they support the use of experimental infection studies for analysing infection patterns in natural as well as laboratory studies, although they do show that extrapolation to field data should be done with caution. Results of controlled infection experiments can then be used to estimate TOI of animals in natural populations, which in turn enables estimating epidemiological parameters (e.g. incidence or basic reproductive number) more accurately than when based on momentary absence/presence information only.

## Ethics Approval

All the procedures followed the Animal Ethics guidelines of the Research Policy of Sokoine University of Agriculture as stipulated in the “Code of Conduct for Research Ethics” (Revised version of 2012), and the guidelines in Sikes and Gannon (Gannon 2007). The used protocol was approved by the University of Antwerp Ethical Committee for Animal Experimentation (2015-69) and adhered to the EEC Council Directive 2010/63/EU.

## Acknowledgements

We thank the staff of the Pest Management Center for support, in particular Shabani Lutea and Geofrey Sabuni. This work was supported by the University of Antwerp and the Antwerp Study Centre for Infectious Diseases (ASCID), University of Antwerp grant number GOA BOF FFB3567 and Deutsche Forschungsgemeinschaft Focus Program 1596. Benny Borremans was a research fellow of Research Foundation Flanders (FWO) during the study, and Joachim Mariën of the Flemish Interuniversity Council (VLIR-UOS). The authors declare that no conflicts of interest exist.

## Supplementary information

### Supplementary text 1: Field work

*Between* 30 July 2013 and 18 October 2013, a rodent capture-mark-recapture (CMR) study was performed in five grids of 100m by 100m in Morogoro, Tanzania. In each grid, 100 Sherman live traps (Sherman Live Trap Co., Tallahassee, FL, USA) were placed at 10m intervals and baited with a mixture of peanut butter and ground maize. Distances between the grids were at least 1.5km (coordinates of the fields are given in supplementary table 1). Trapping sessions of three consecutive nights each were repeated every other week. Captured M. natalensis were transported to the nearby laboratory (Pest Management Centre - Sokoine University of Agriculture, approx. 2km from the field sites), where weight, sex and reproductive status were determined, and mice were individually marked using toe clipping (Borremans, Sluydts, et al., 2015). Blood samples were taken from the retro-orbital sinus and preserved on prepunched filter paper (±15 mL/punch; Serobuvard, LDA 22, Zoopole, France). Saliva was collected by putting a small filter paper slip into the mouth of the animal for a few seconds. If the animal urinated, a urine sample was taken directly from the animal using filter paper. After sampling, animals were again released on the exact capture location. Filter papers were dried and stored in the dark, at ambient temperature (maximum temperature was 28°C) for two months, after which they were preserved at −20°C in a locked plastic bag with dehydrating silica gel, as described by Borremans (Borremans 2014).

**Table 1:** Coordinates of field on which rodents were trapped

| Field | Corner | Coordinates |
| --- | --- | --- |
| F1 | 1a | S06° 50.695' E037° 39.240' |
| F1 | 10a | S06° 50.675' E037° 39.202' |
| F1 | 1j | S06° 50.633' E037° 39.227' |
| F1 | 10j | S06° 50.657' E037° 39.262' |
| F3 | 1a | S06° 50.640' E037° 39.630' |
| F3 | 10a | S06° 50.600' E037° 39.617' |
| F3 | 1j | S06° 50.624' E037° 39.578' |
| F3 | 10j | S06° 50.664' E037° 39.599' |
| F4 | 1a | S06° 51.604' E037° 37.771' |
| F4 | 10a | S06° 51.573' E037° 37.791' |
| F4 | 1j | S06° 51.595' E037° 37.832' |
| F4 | 10j | S06° 51.595' E037° 37.832' |
| F5 | 1a | S06° 51.279' E037° 37.704' |
| F5 | 10a | S06° 51.237' E037° 37.726' |
| F5 | 1j | S06° 51.302' E037° 37.747' |
| F5 | 10j | S06° 51.301' E037° 37.758' |
| F6 | 1a | S06° 51.044' E037° 37.969' |
| F6 | 10a | S06° 51.002' E037° 37.995' |
| F6 | 1j | S06° 51.604' E037° 37.771' |
| F6 | 10j | S06° 51.064' E037° 38.015' |

### Supplementary text 2: Detection and quantification of antibody and MORV RNA

All individuals that were captured during at least two different trapping sessions were analysed for the presence of anti-MORV antibodies (Abs) and viral RNA (vRNA). Blood samples were analysed for the presence of Abs by indirect immunofluorescence assay using MORV-infected Vero cells *as antigens and* polyclonal rabbit anti-mouse IgG (Dako, Denmark) as secondary antibodies. Dried blood spots on filter paper were punched out and eluted in phosphate buffer saline. Antibody titers were estimated using two-fold dilution series, starting with a minimum dilution of 1:10. Viral RNA extraction was performed on both blood and excretion (saliva and urine) samples as described by Borremans et al., (Borremans, Vossen, et al., 2015), using the QIAmp vRNA Mini Kit (Qiagen, Hilden, Germany). RT-PCR was done following the protocol described in Günther et al., 2009. MoroL3359-forward and MoroL3753-reverse primers were used to target a 340 nucleotide portion of the RNA-dependent RNA polymerase gene of MORV (Günther, et al., 2009). All amplicons were confirmed by Sanger-sequencing at the Vlaams Instituut voor Biotechnologie (Antwerp, Belgium) and comparing the sequences to known MORV sequences in the software Geneious 7.0.6 (Kearse et al., 2012).

### Supplementary text 3:Definition of a good match between field and laboratory data

We defined a good match for a field tested individual if all the Ab titers and/or vRNA presence/absence data fell within a particular confidence band (CB). This CB was calculated following Borremans et al (Borremans et al., 2016), and adjusted such that, given the number of recaptures for this particular individual, there is 95% chance that all observations fall within the CB. Consequently, the width of the CB depends on the number of recaptures of the individual. For each TOI method we calculated the percentage of *individuals* that matched the laboratory data. Note that the use of a 95% CB implies that in the theoretical situation where the Ab and/or vRNA temporal dynamics are identical to those measured in the laboratory, 95% of the mice would match the laboratory data. If the dynamics are different to those measured in the lab, one expects that the percentage of matches would be lower. We also calculated the percentage of *observations* that matched (fell inside) the laboratory data’s 95% CB for each TOI method. Again, in case of dynamics similar to those in the laboratory, 95% of the data would match the laboratory data, and this percentage will be lower if the dynamics are different.

## Supplementary figures

### Explanation how to interpret the figures

The supplementary information contains the Ab titer and viral RNA detection results for all individuals of which the TOI was estimated (i.e. individuals that were recaptured and found the be Ab-positive at least once). Each of the three panel Figs shows all samples for each individual separately, where large coloured dots represent field data and a small black dots laboratory data. On each plot, the results are shown two times, once for each TOI estimation method that estimates the best fit to the laboratory data based on Ab titers, vRNA presence in blood and/or excretions and weight (explained in the methods section of the main text). The estimation was done based on Ab titers only (method 1, red dots), or including information on both Ab titer and vRNA presence in blood/excretions (method 2, green dots). Fig S7, for example, should be interpreted as follows: this individual was sampled twice, where both blood samples were Ab-positive (titers 12000 and 6000), one blood sample vRNA-positive (detection probability one), and the two excretion samples vRNA-negative (detection probability zero). When taking into account only Ab titer information, the best possible fit to laboratory data is found at day 9, which means that the most likely time between infection and first sampling was 9 days. The red dots on the left panel indeed show that the samples fit well within the laboratory data (black dots). When only taking into account both virus and Ab data (method 2, green dots), we observe a similar match for the Ab titer data (green dots on the left panel). The green dots shown on the right panels, plotted again on day 9, show that there is a bad match for vRNA presence in blood, as the probability of observing vRNA in blood is almost 0 at that point (the black lines on the right panels show the probability - not the quantity - of being vRNA-positive at a certain time since infection in the laboratory). There was a good match for vRNA in excretions (lower right panel).

When a sample was inconclusive for the presence of vRNA, it was not shown in the right panels nor used for TOI estimation. As a consequence, some Ab titer panels contain more sample data points than the vRNA panels.

**Fig S1:**
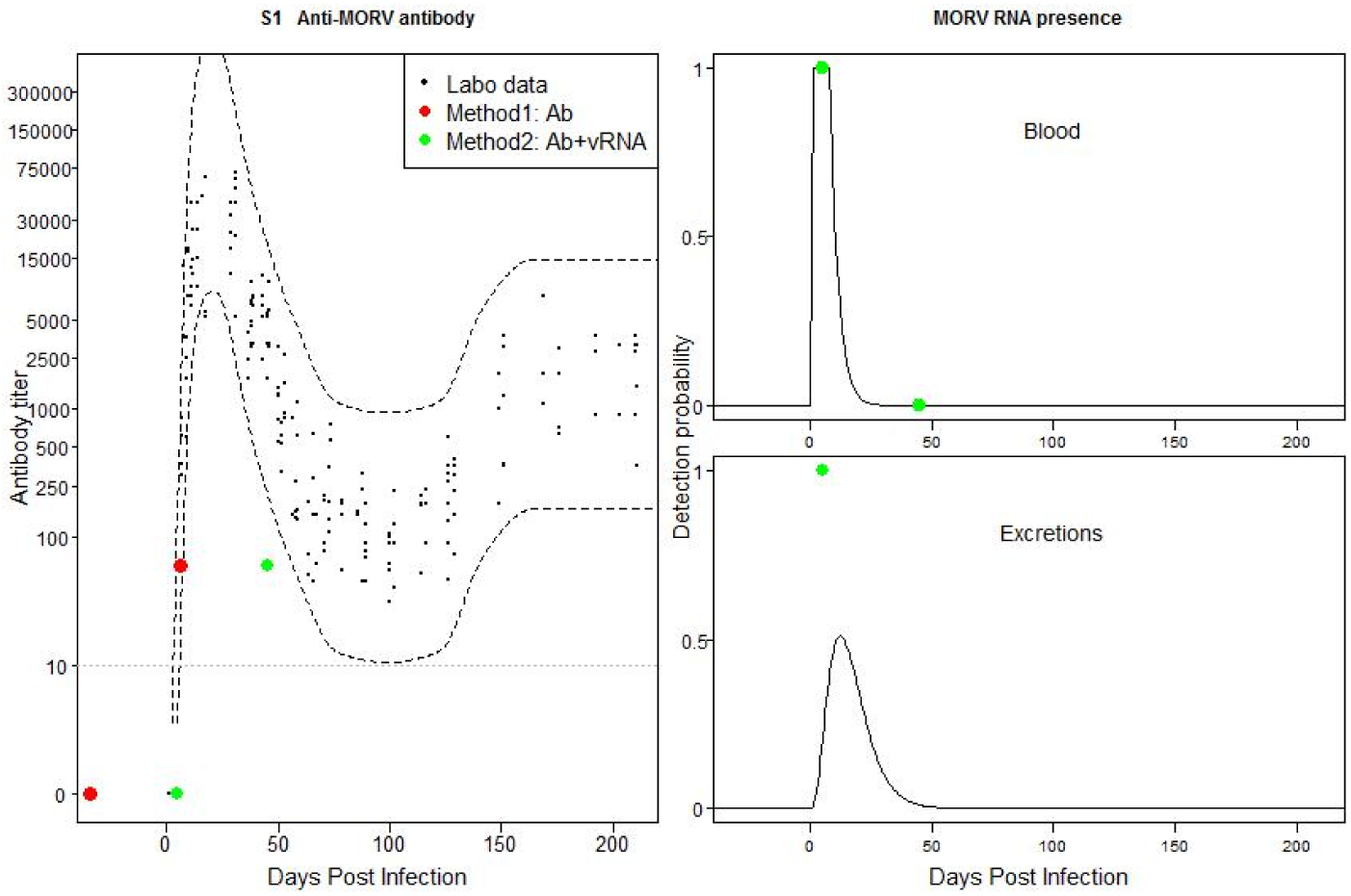
Individual 354060F1: Mismatch between field and laboratory data based on time of infection method 2. Possible recently acquired active infection with Ab titer that just falls out of the confidence band (CB) of the laboratory data.

**Fig S2:**
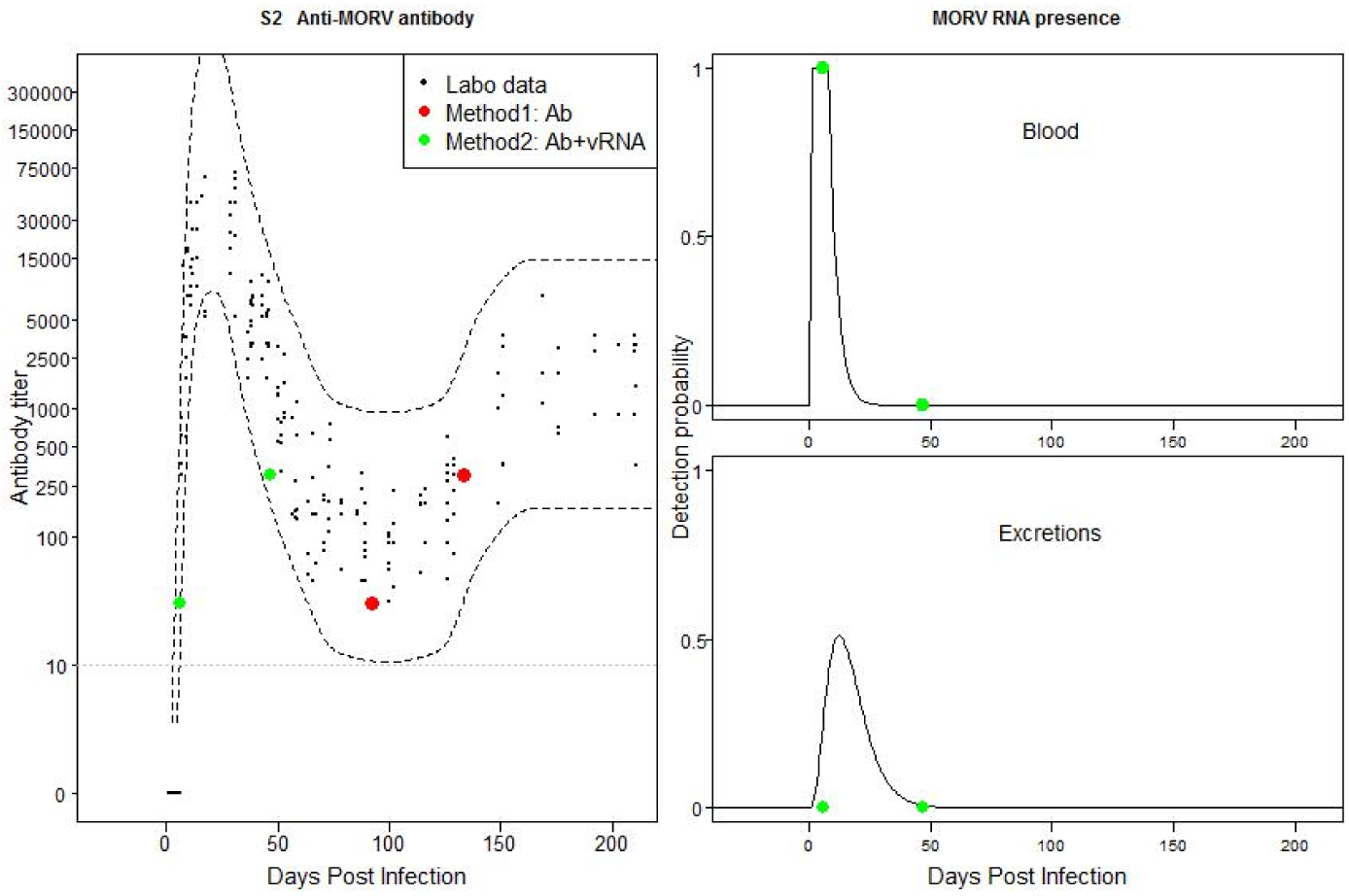
Individual 364060F1: Mismatch between field and laboratory data based on method 2. Possible recently acquired active infection with Ab titer that just falls out of the CB of the laboratory data.

**Fig S3:**
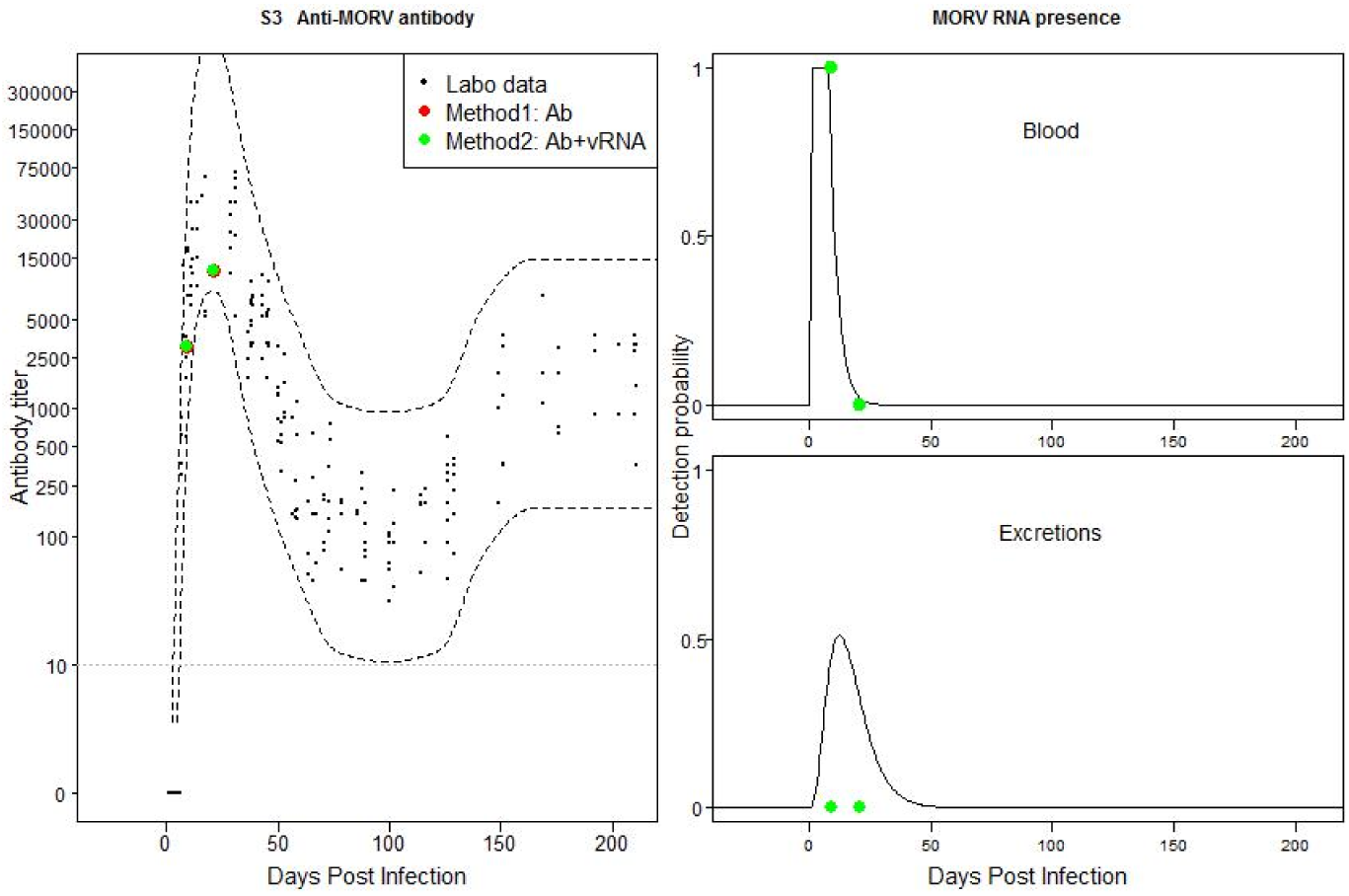
Individual 371090F1: Match between field and laboratory data. Possible recently acquired active infection.

**Fig S4:**
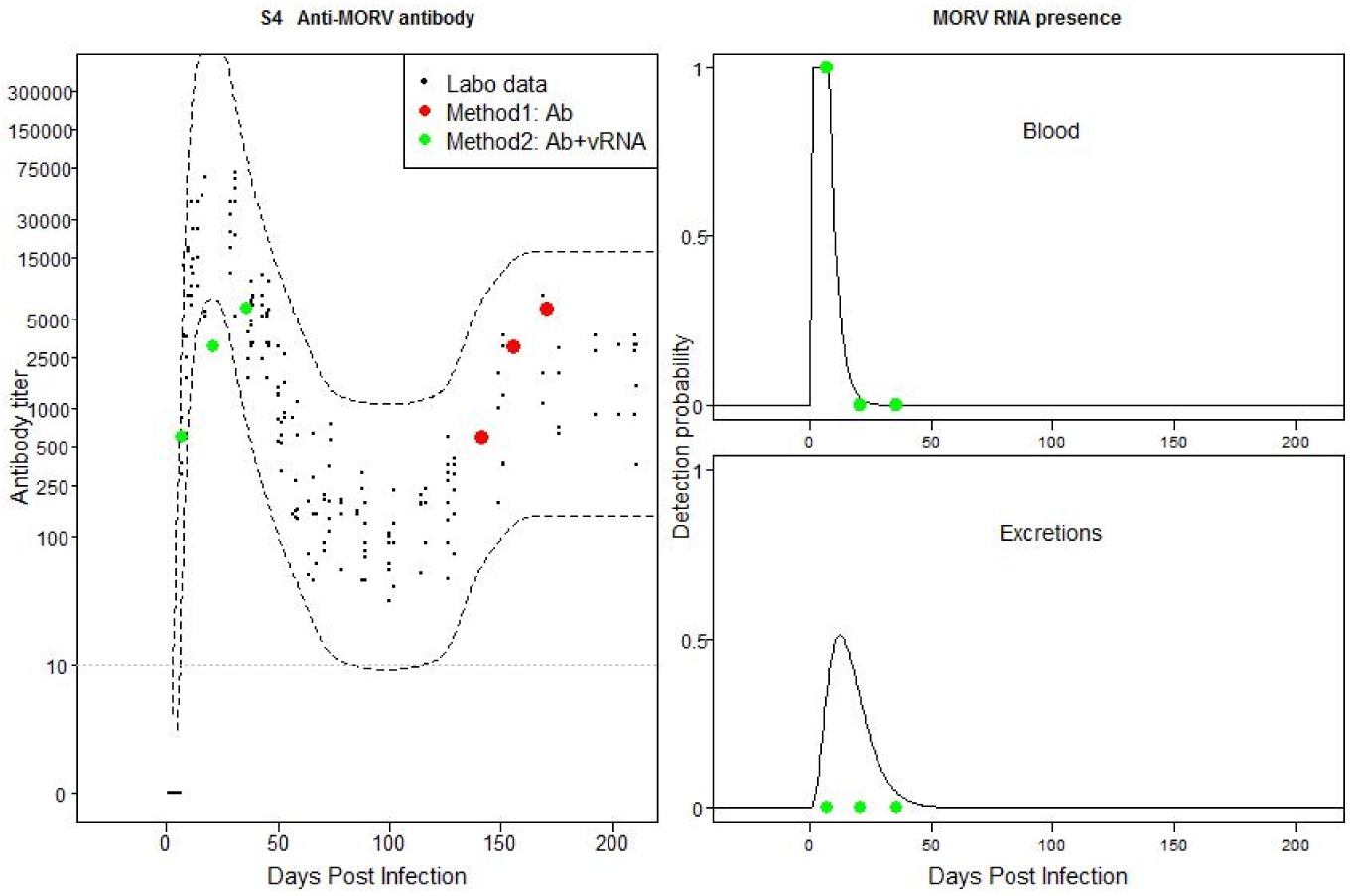
Individual 3510100F1: Mismatch between field and laboratory data based on method 2. Possible recently acquired active infection with Ab titer that just falls out of the CB of the laboratory data.

**Fig S5:**
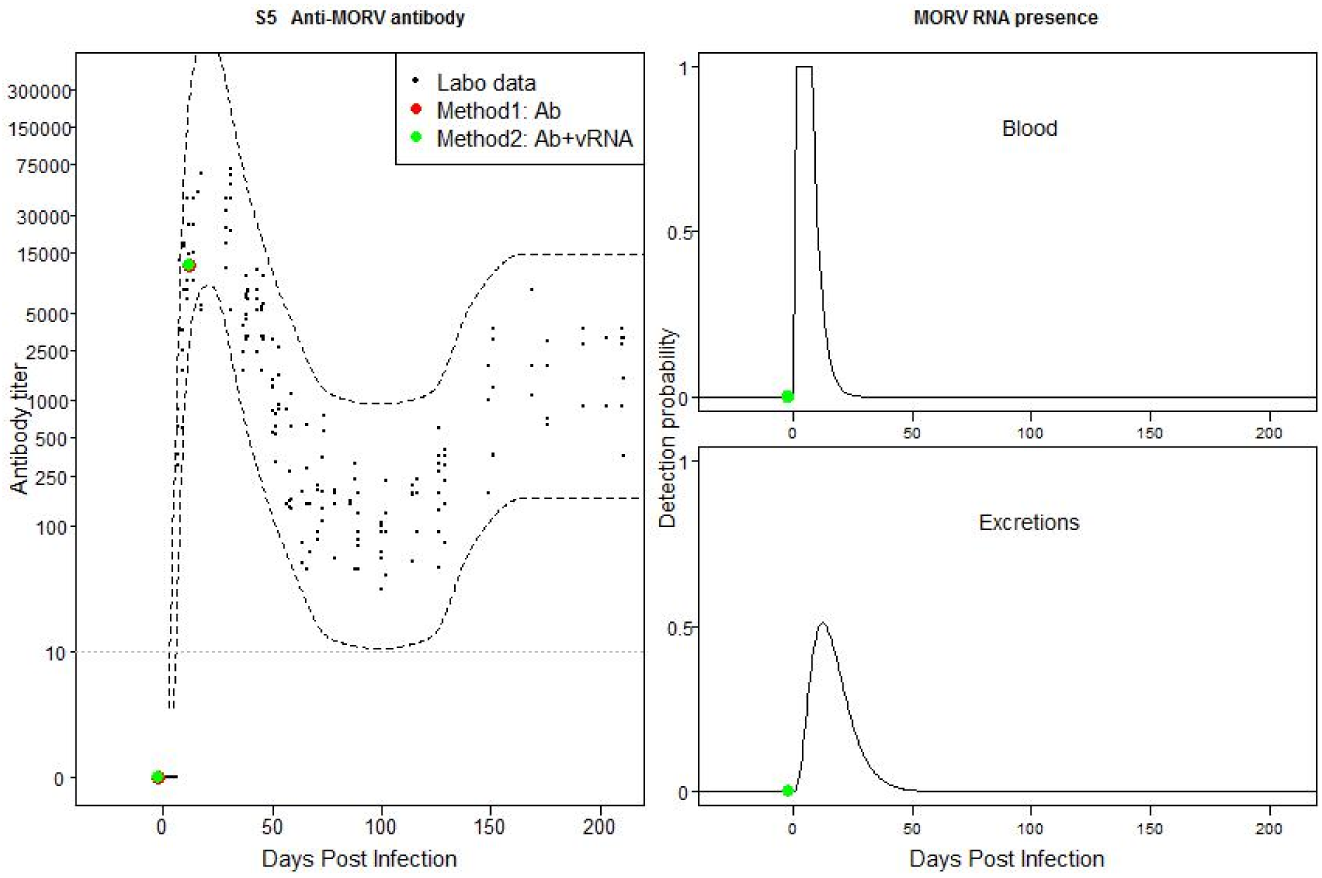
Individual 3710100F1: Match between field and laboratory data. Possible recently acquired infection without evidence for viraemia.

**Fig S6:**
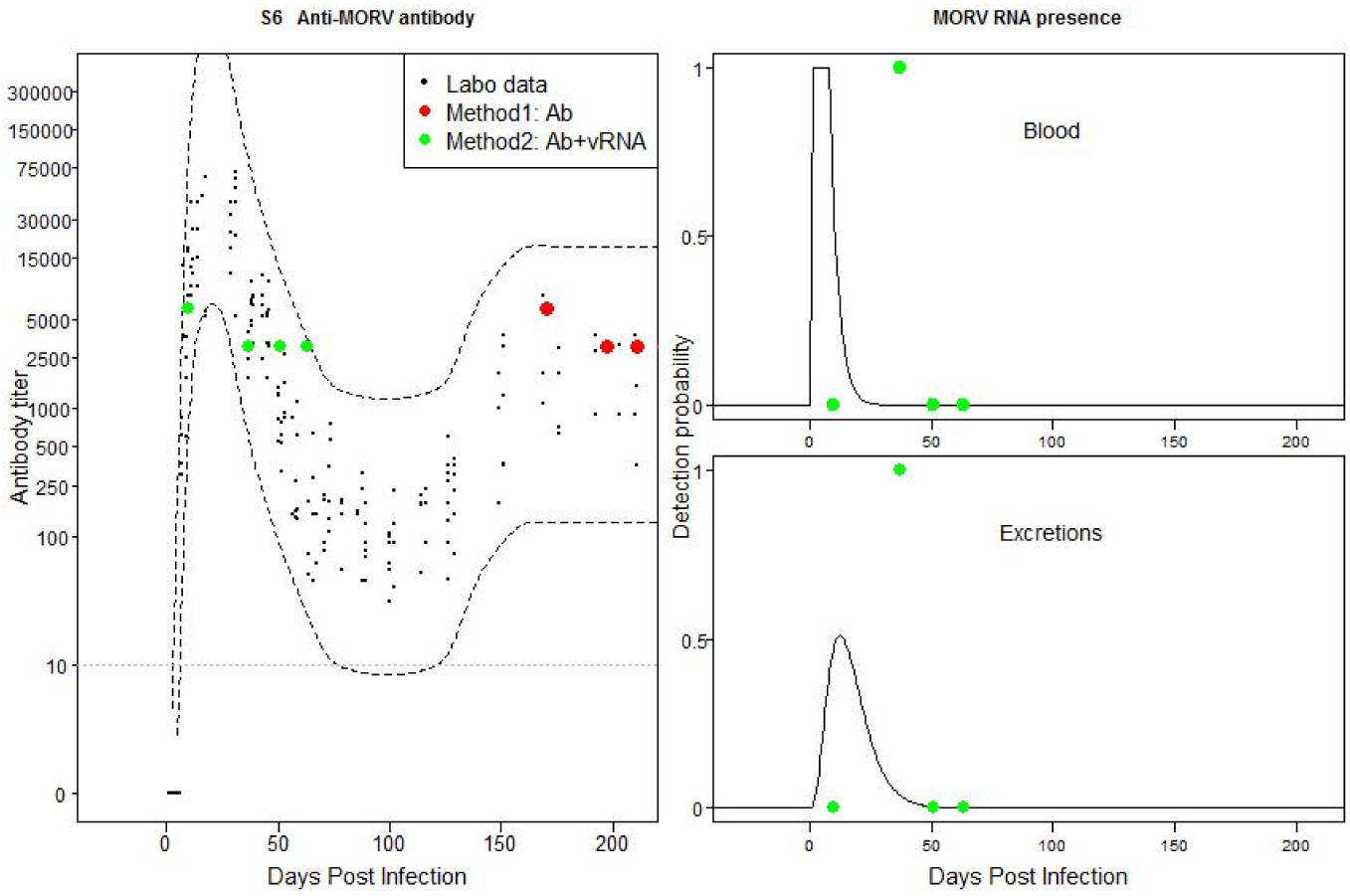
Individual 15F3a: Mismatch between field and laboratory data based on method 2. Possible recently acquired infection with temporally flare-ups of MORV in blood and excretions.

**Fig S7:**
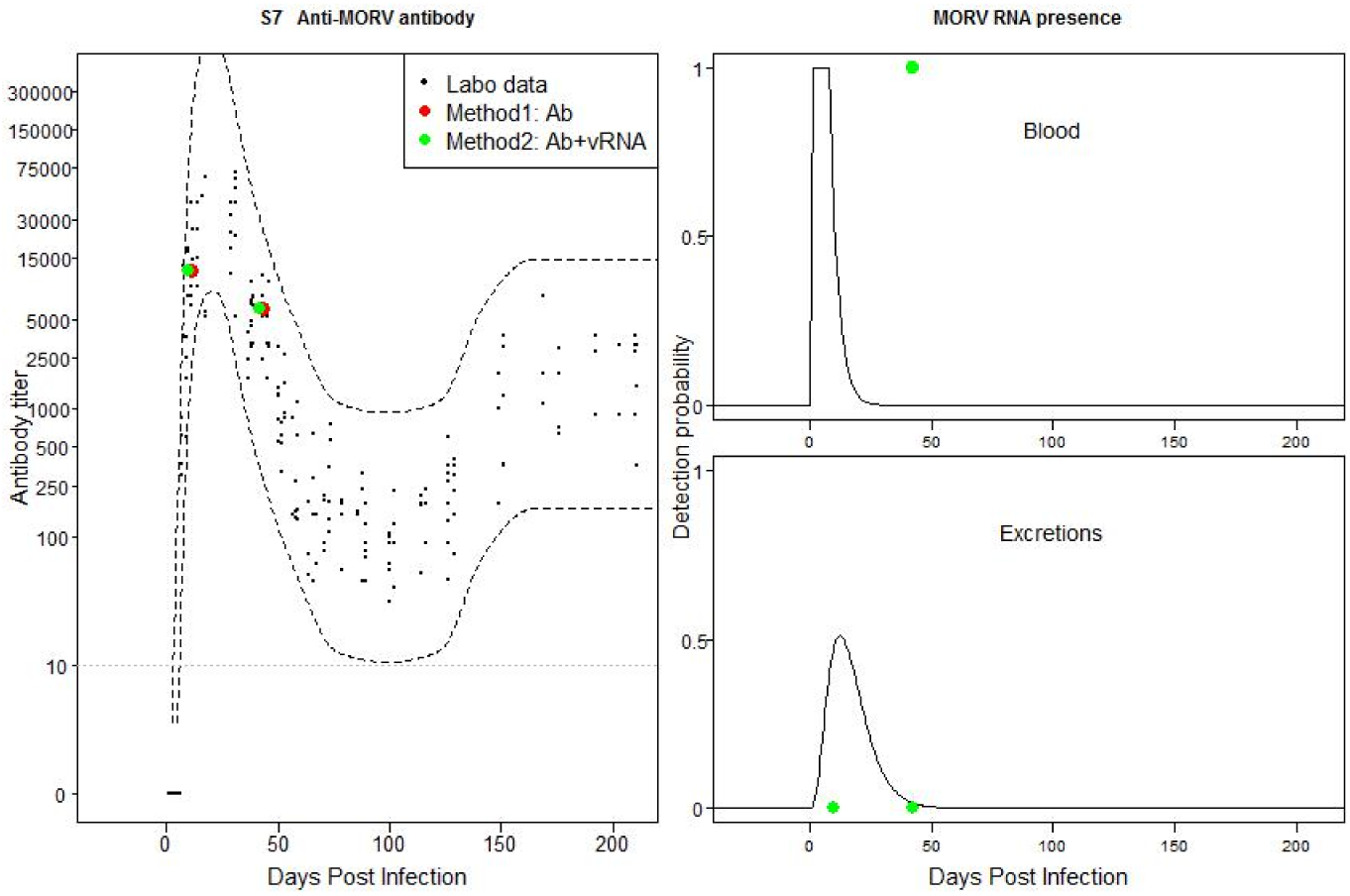
Individual 15 F3b: Mismatch between field and laboratory data based on method 2. Possible recently acquired infection with temporally flare-ups of MORV in blood.

**Fig S8:**
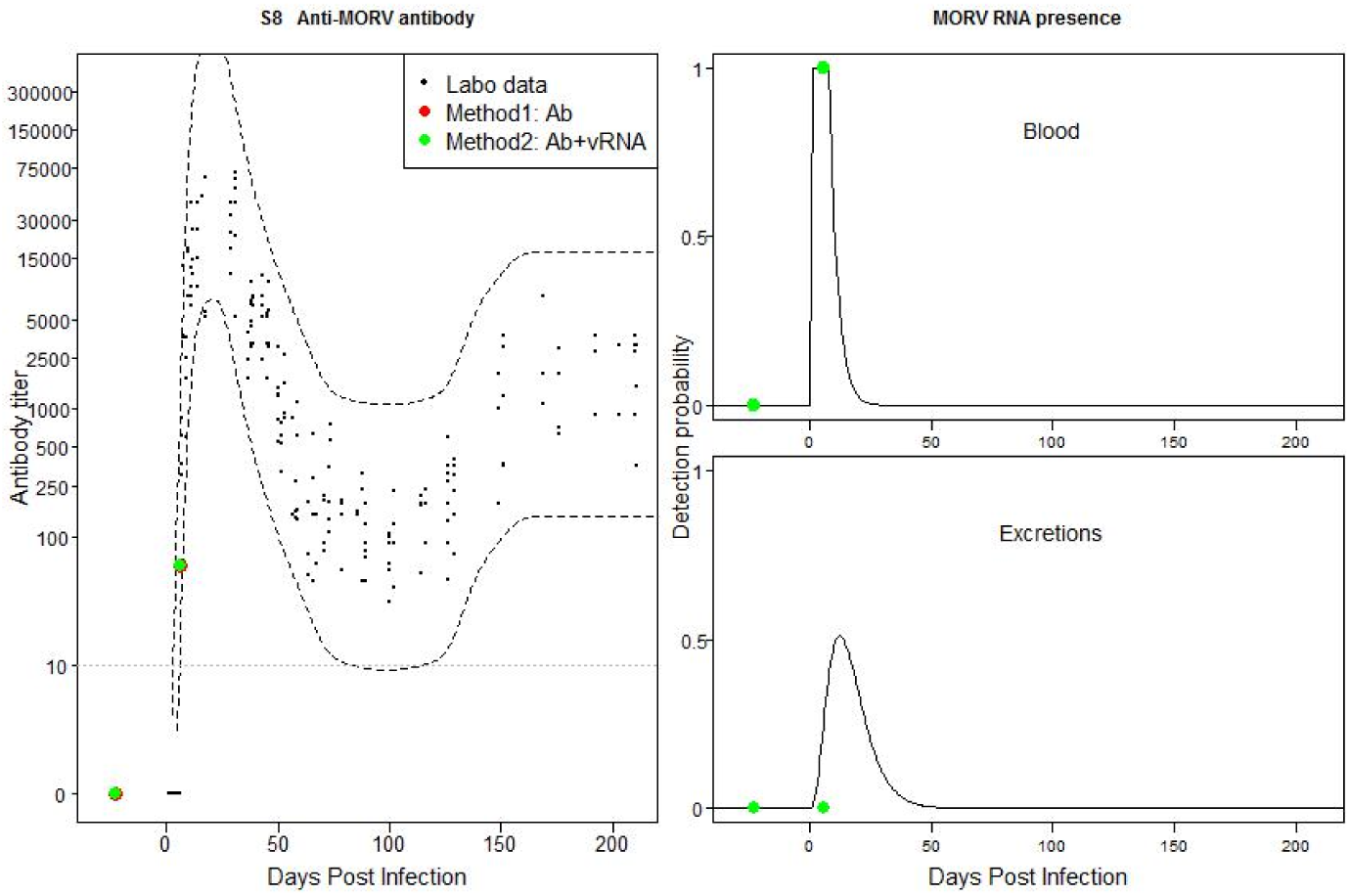
Individual 38F3: Match between field and laboratory data. Possible recently acquired active infection.

**Fig S9:**
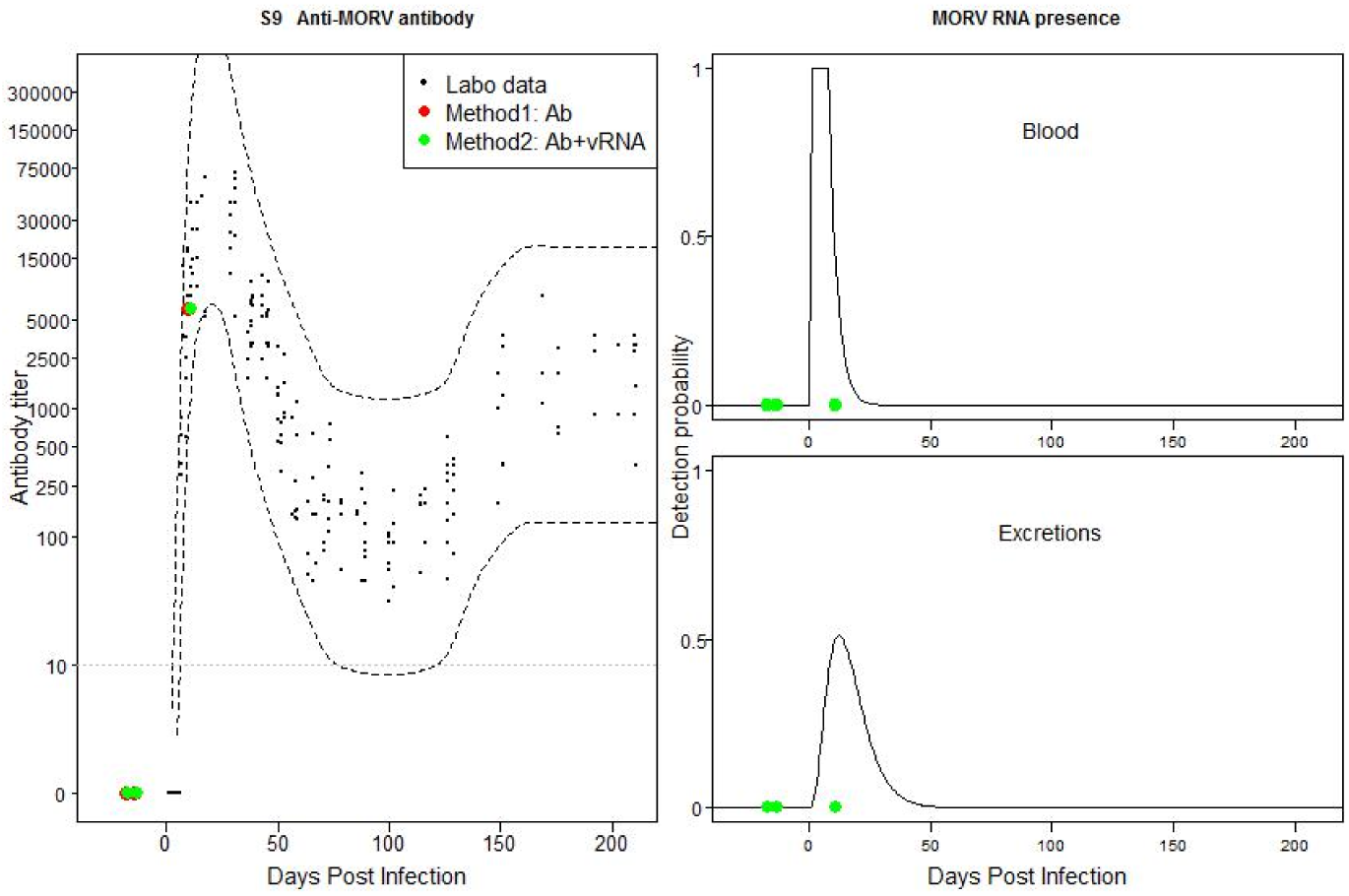
Individual 210 F3: Match between field and laboratory data. Possible recent infection without positive vRNA samples

**Fig S10:**
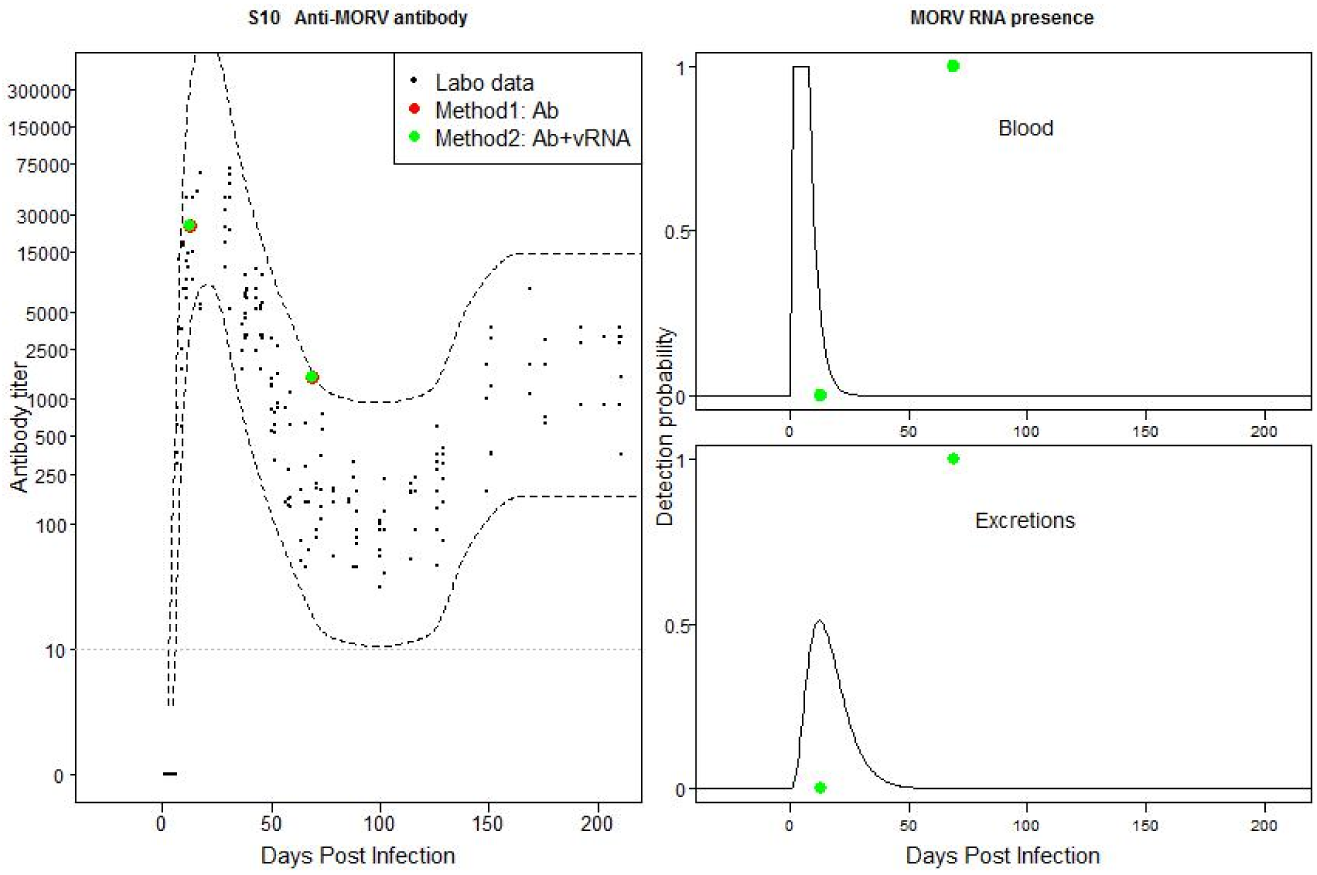
Individual 330F3: Mismatch between field and laboratory data based on method 2. Possible recently acquired infection with temporally flare-up of MORV in blood and excretions. Because the animal is still young (weight at first capture is 20g), the Ab titer pattern might also be explained by the presence of maternal Abs, followed by an vRNA-positive blood and excretion sample.

**Fig S11:**
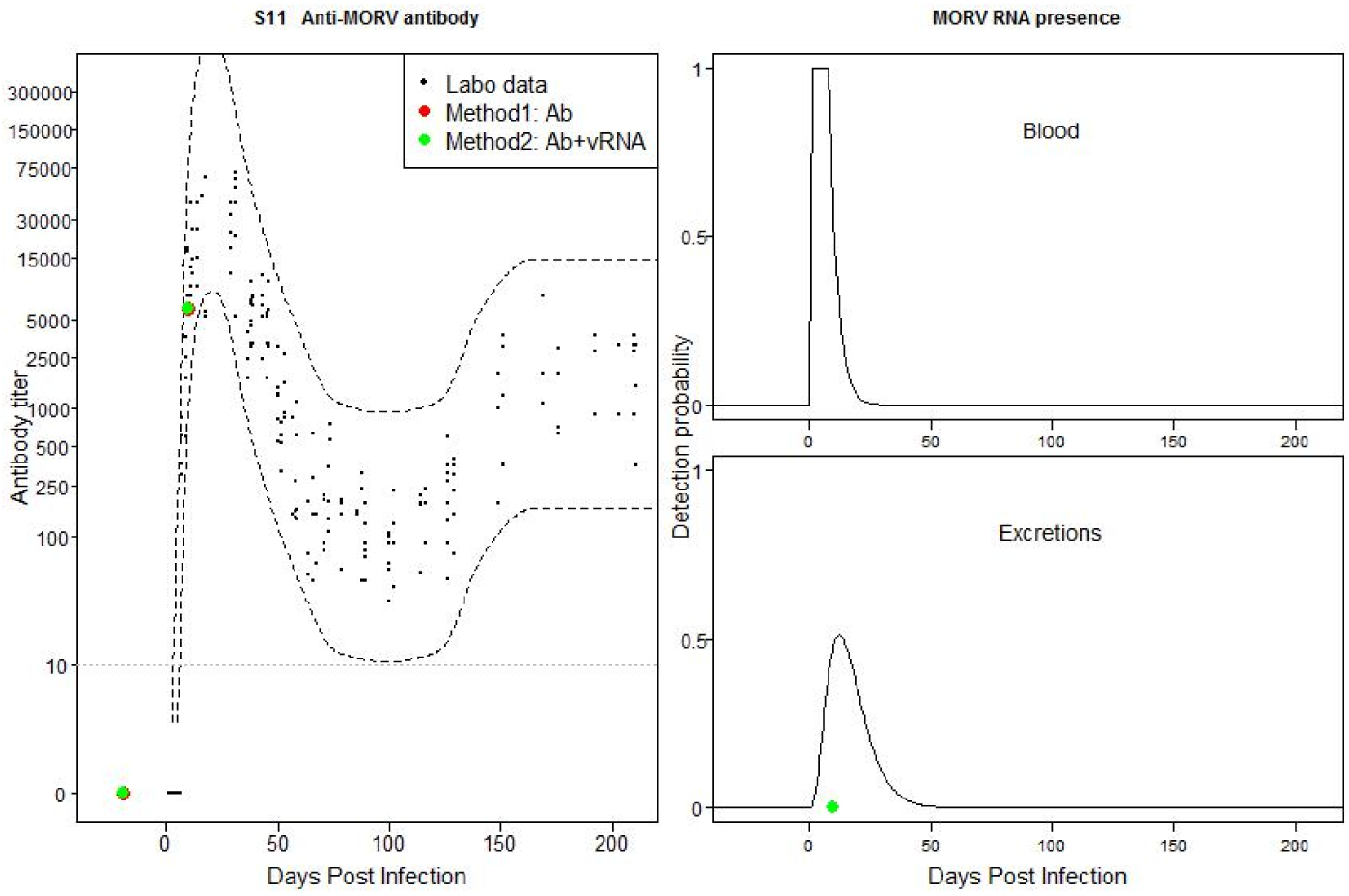
Individual 340F3: Match between field and laboratory data. Possible recently acquired active infection without evidence for positive vRNA sample.

**Fig S12:**
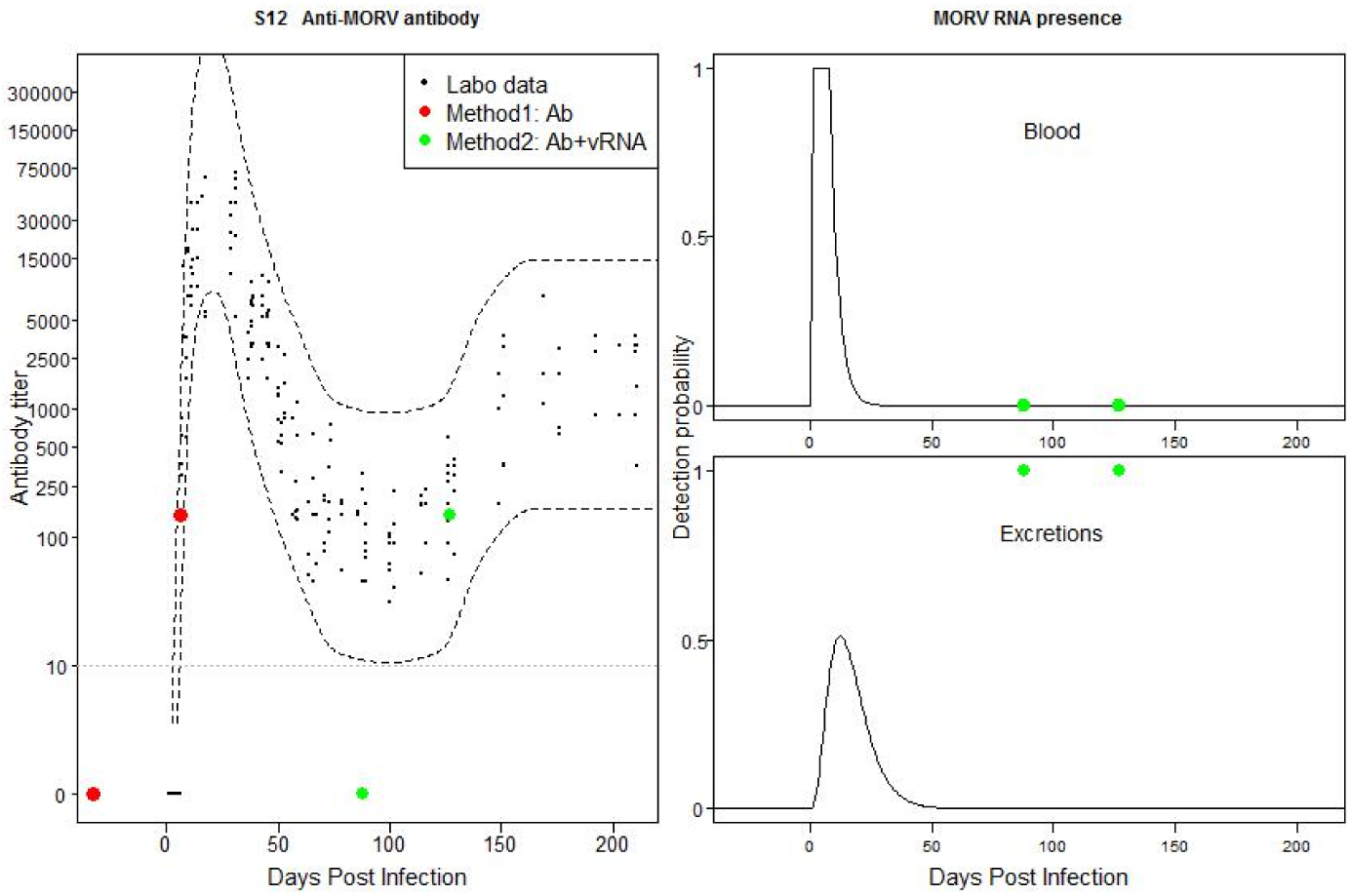
Individual 390 F3: Mismatch between field and laboratory data based on method 2. Possible recently acquired active infection with Ab titer that just fell out of the confidence interval of the laboratory data and presence of MORV in excretions.

**Fig S13:**
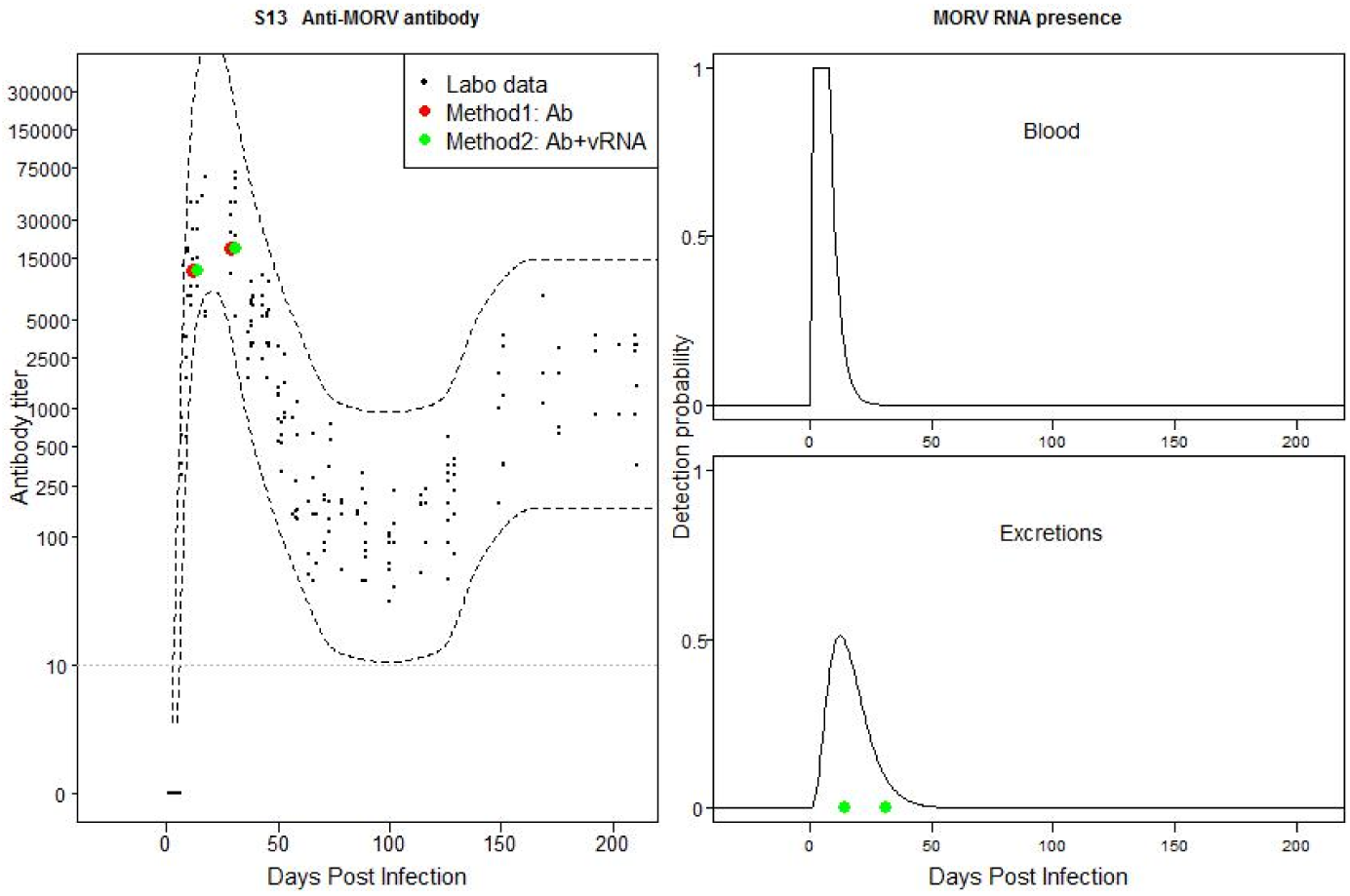
Individual 550F3: Match between field and laboratory data. Possible past, old acute infection.

**Fig S14:**
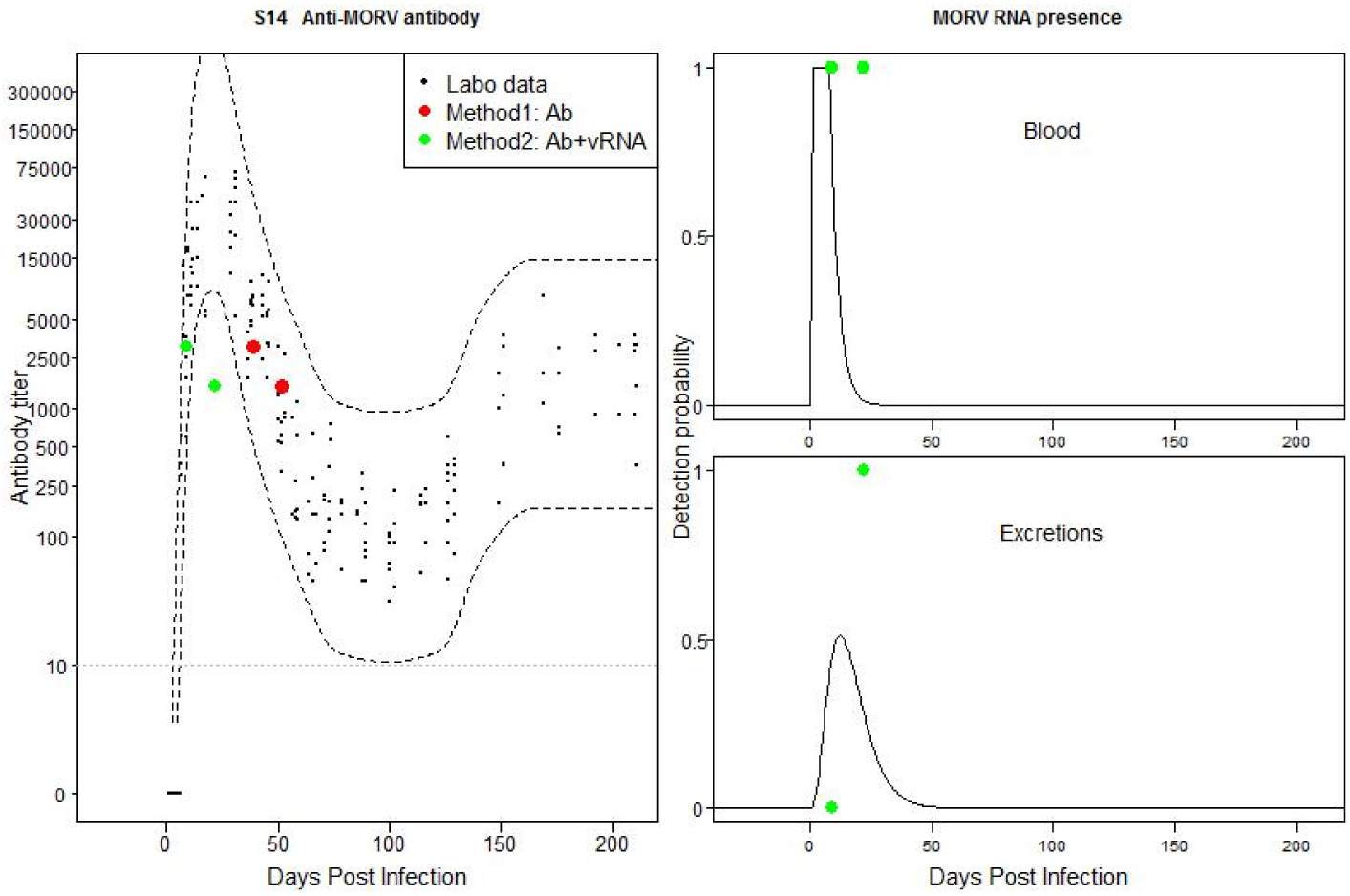
Individual 1670F3: Mismatch between field and laboratory data based on method 2. Possible recently acquired active infection with Ab titer that falls out of the confidence interval of the laboratory data.

**Fig S15:**
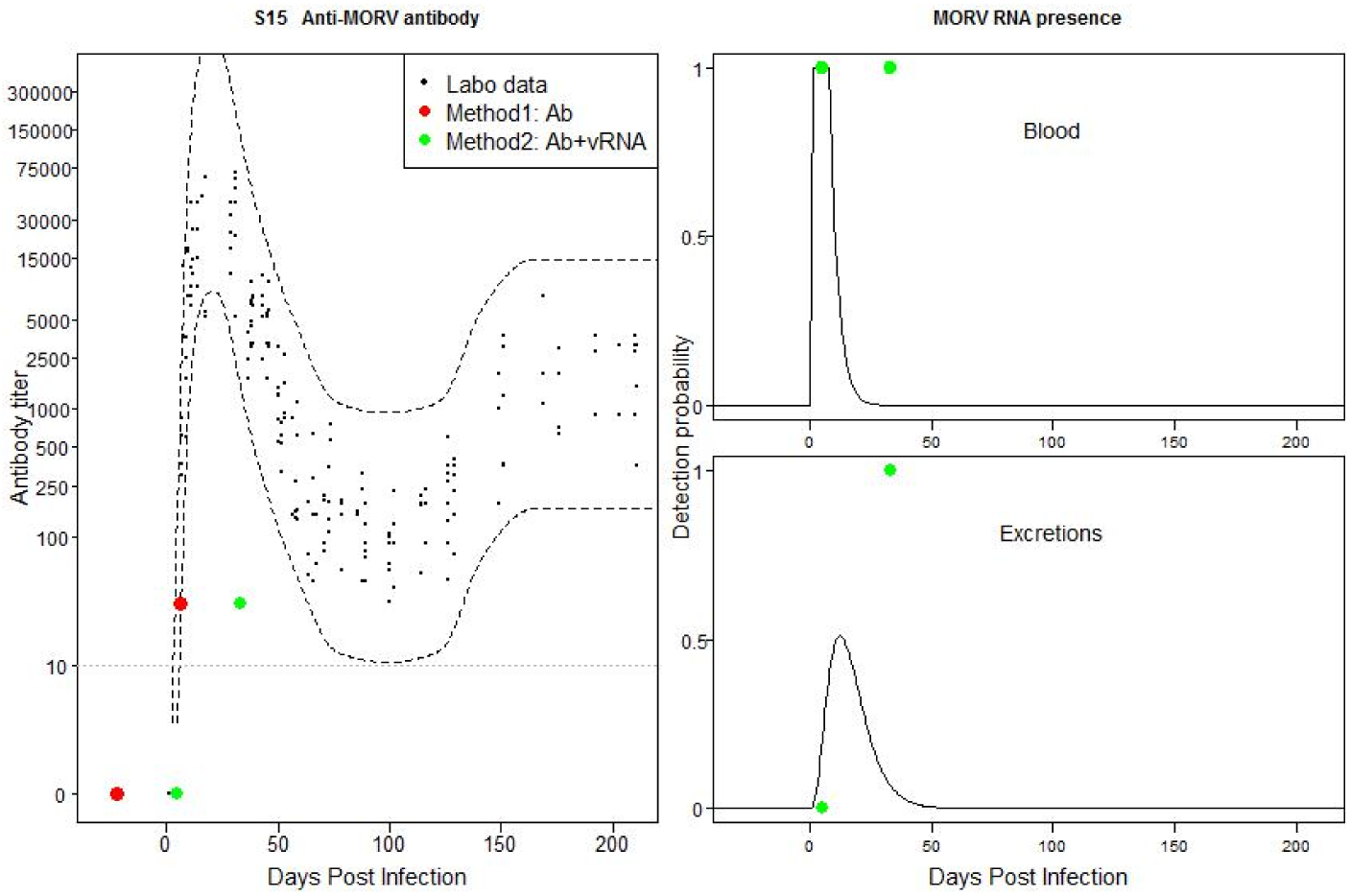
Individual 20100F3: Mismatch between field and laboratory data based on method 2. Possible recently acquired active infection with Ab titer that falls out of CB of the laboratory data.

**Fig S16:**
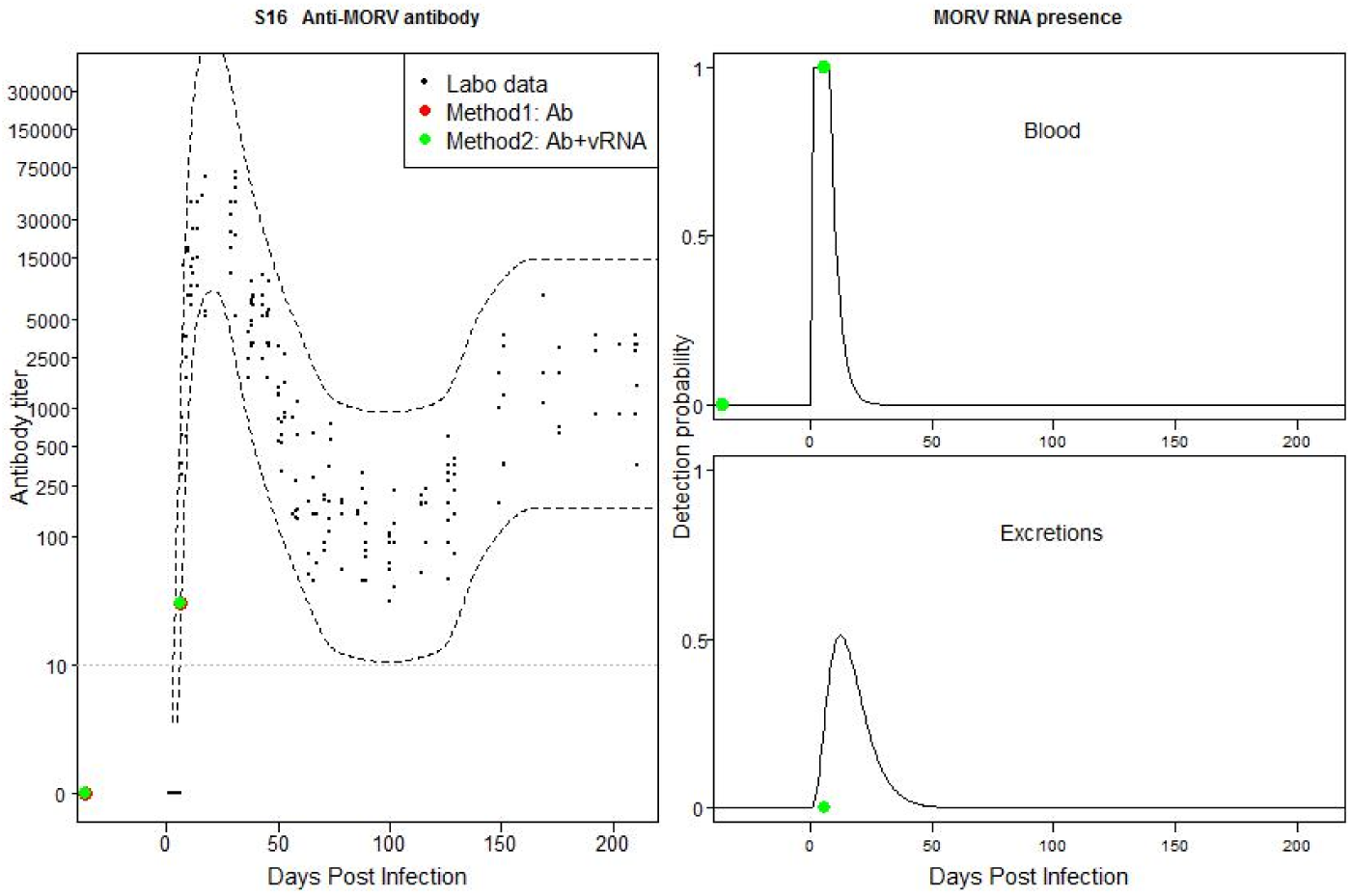
Individual 140F6: Match between field and laboratory data. Possible recently acquired active infection.

**Fig S17:**
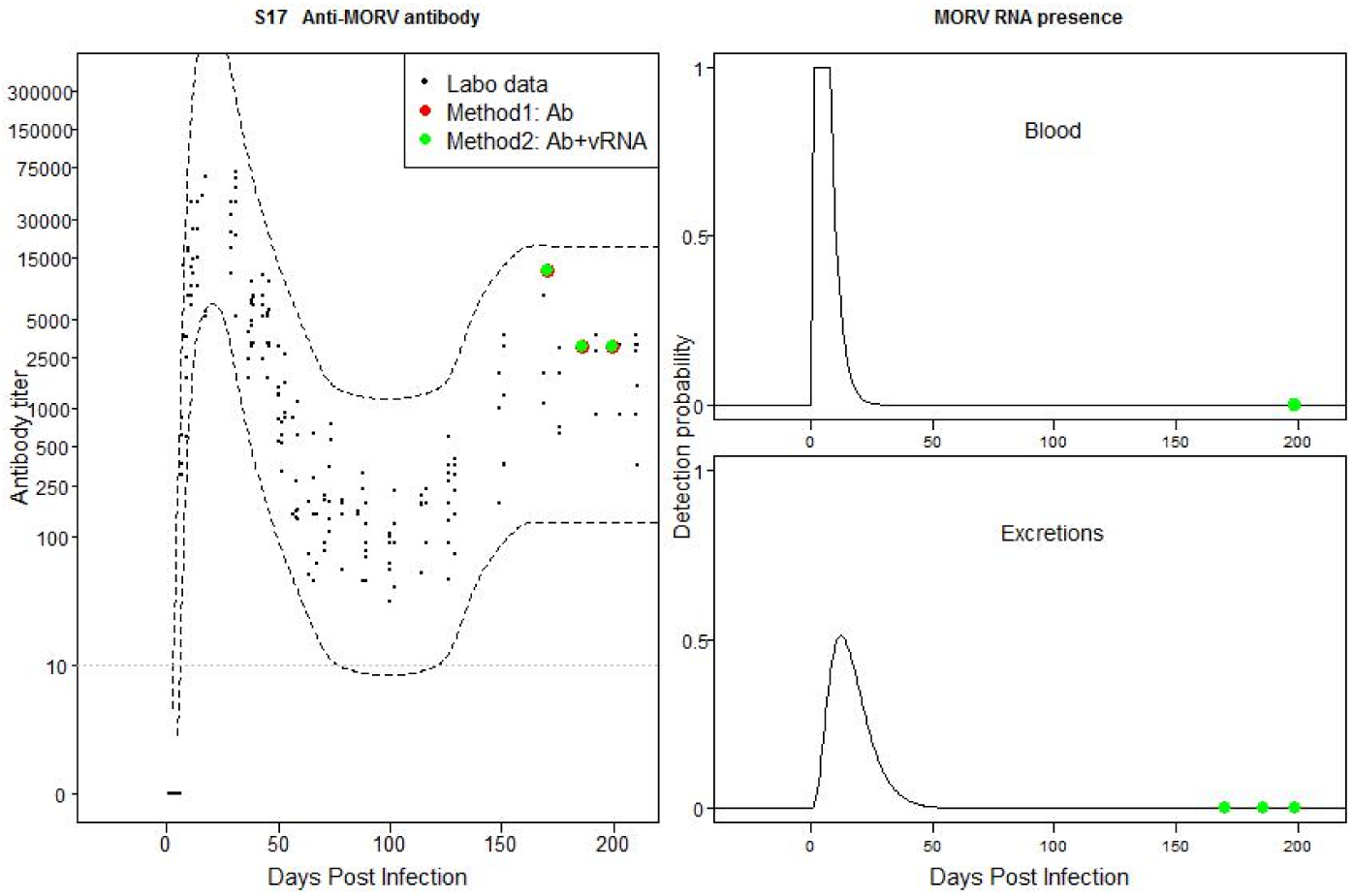
Individual 360F4: Match between field and laboratory data. Possible past old acute infection.

**Fig S18:**
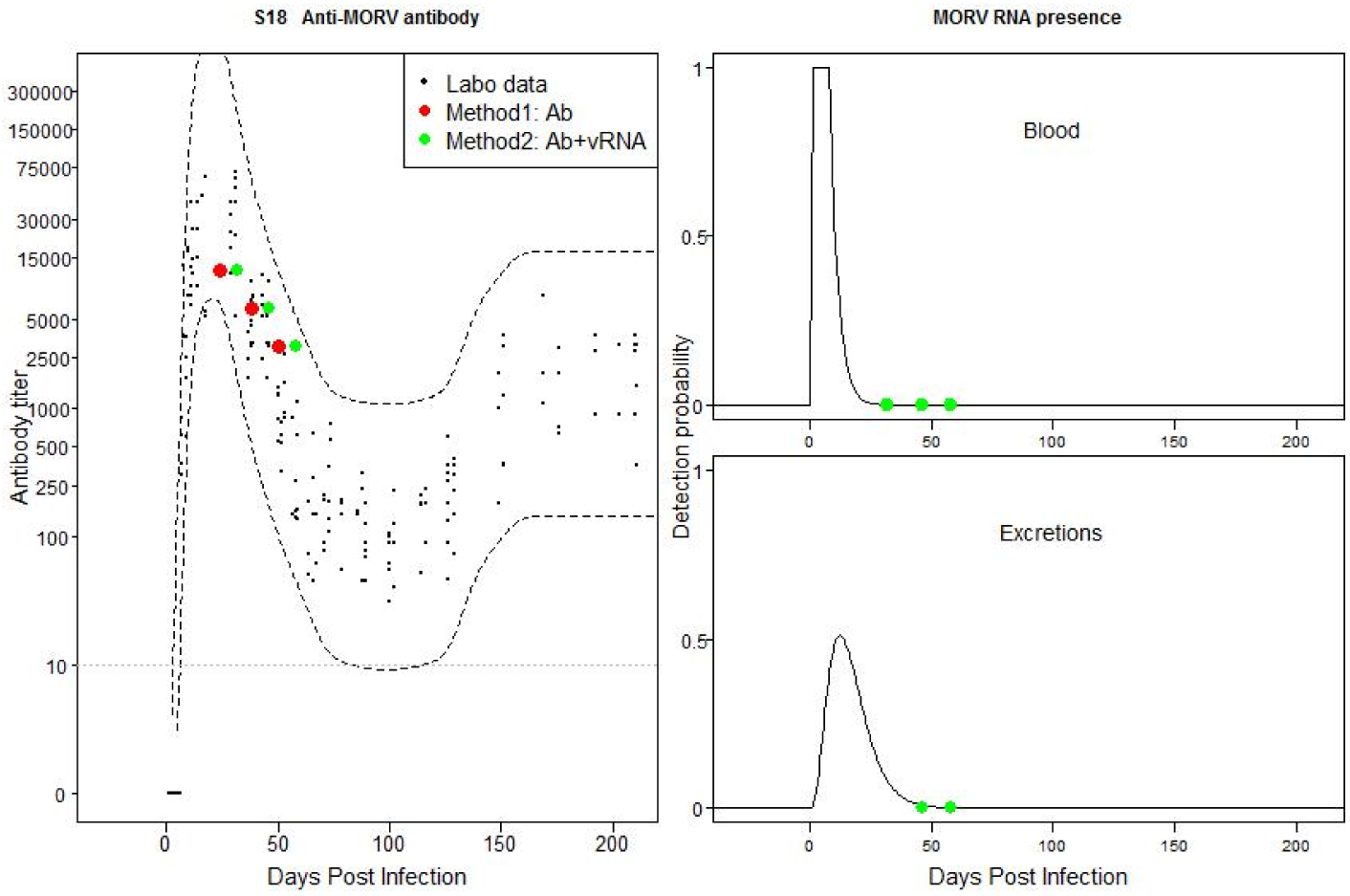
Individual 890F4: Match between field and laboratory data. Possible past old acute infection.

**Fig S19:**
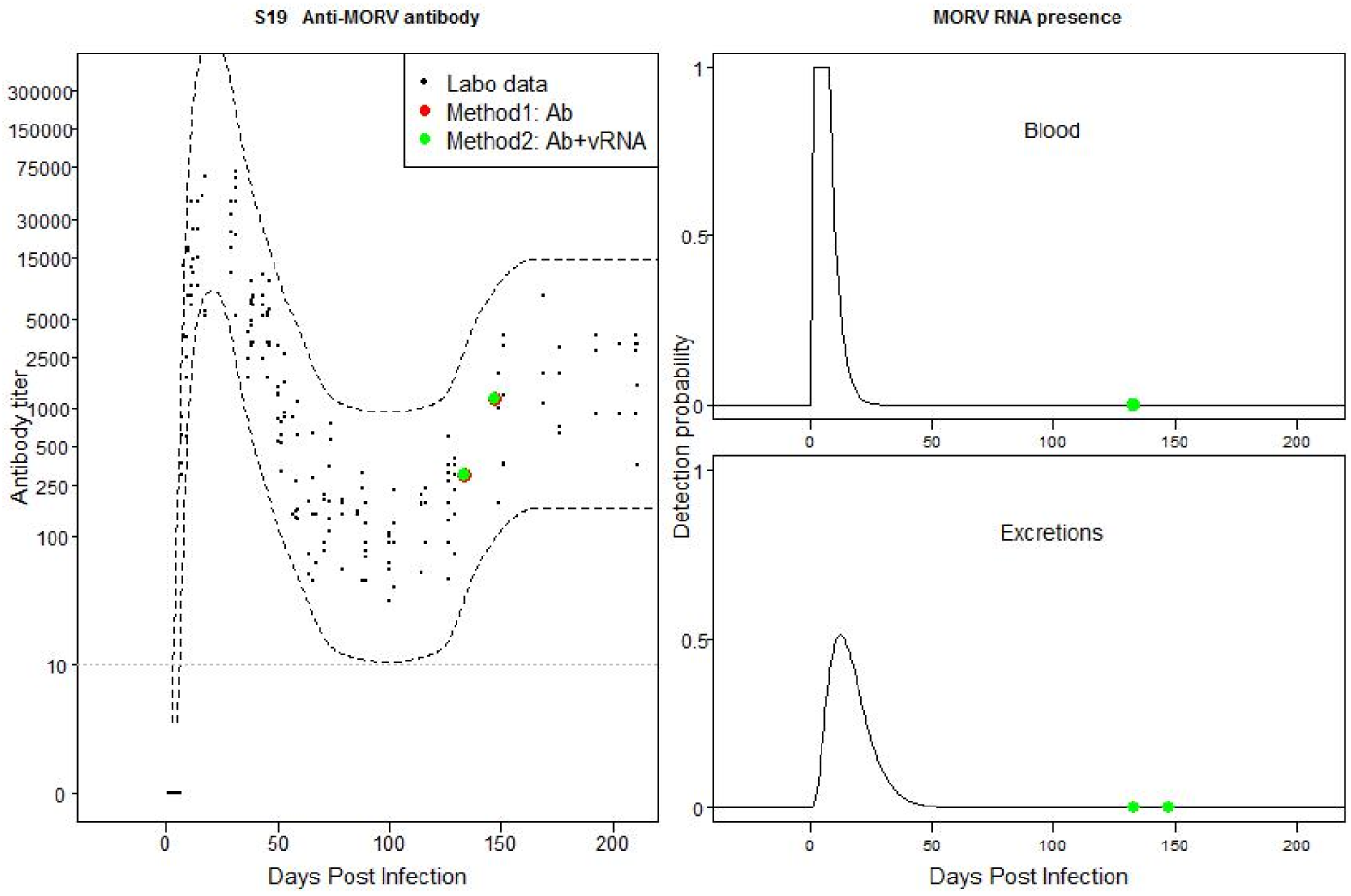
Individual 1100F4: Match between field and laboratory data. Possible past old acute infection.

**Fig S20:**
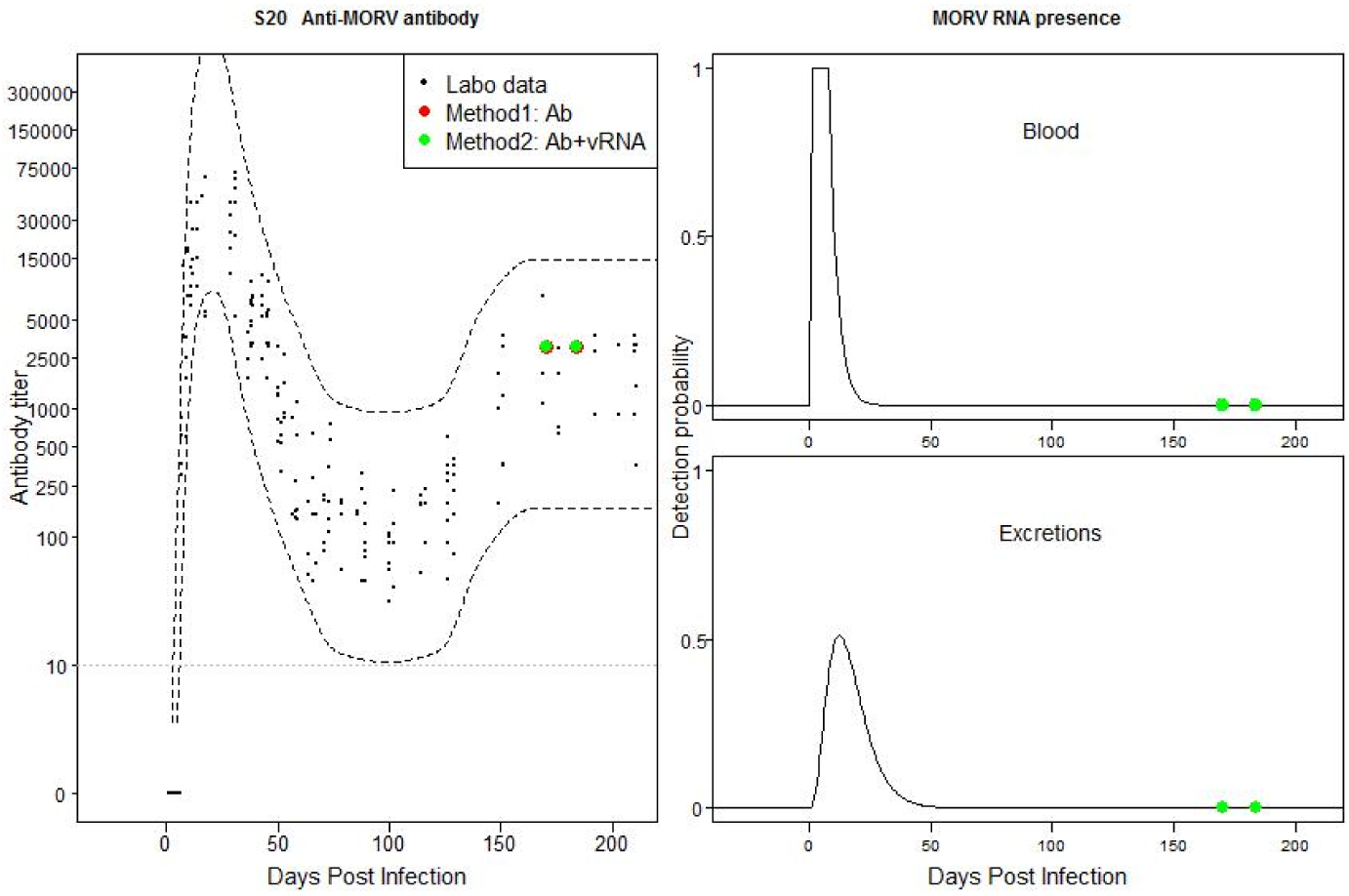
Individual 4080F4: Match between field and laboratory data. Possible past old acute infection.

**Fig S21:**
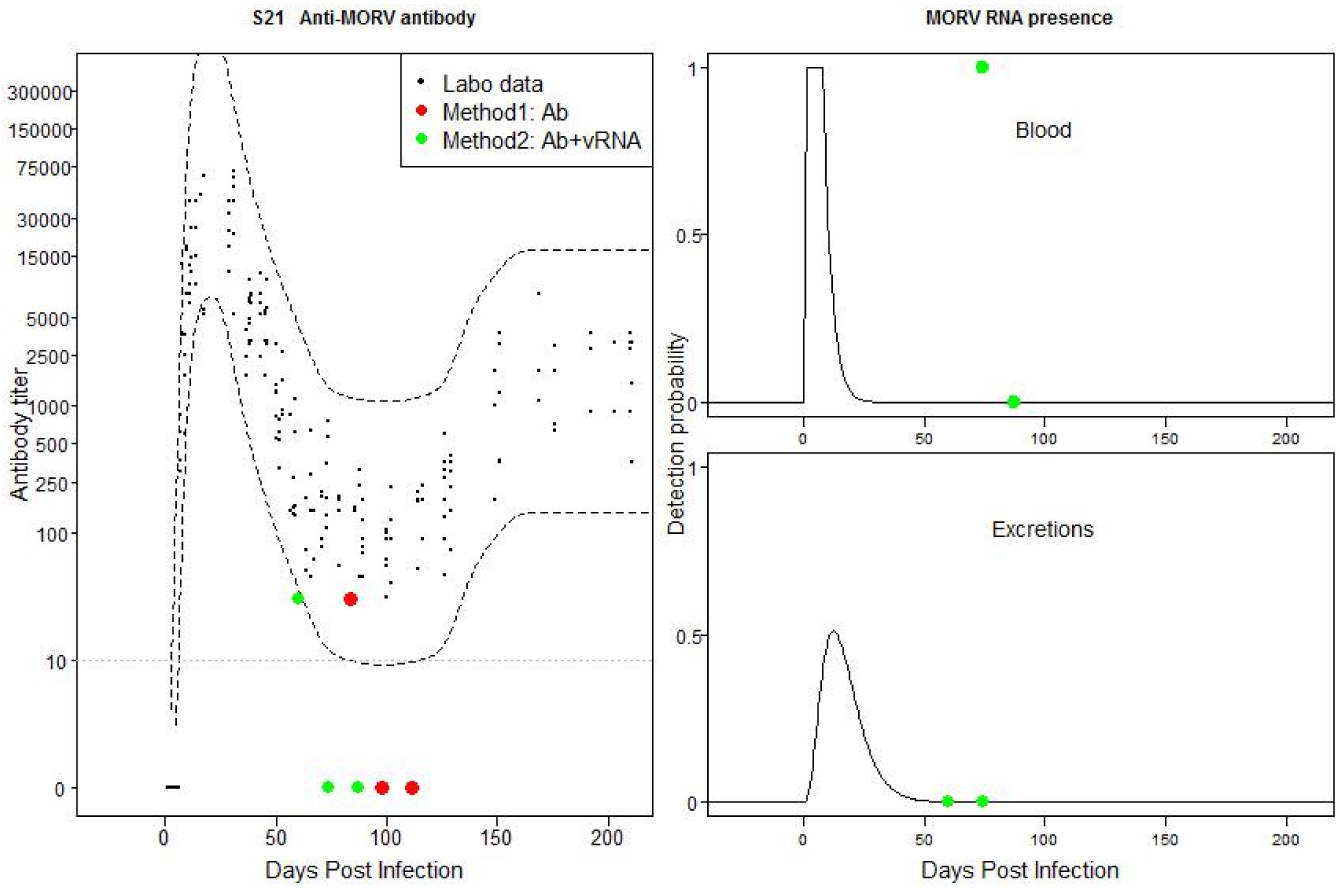
Individual 260F5: Mismatch between field and laboratory data based on both TOI method estimations. Negative Ab titers could be low titers. Possible flare-ups of vRNA in blood.

**Fig S22:**
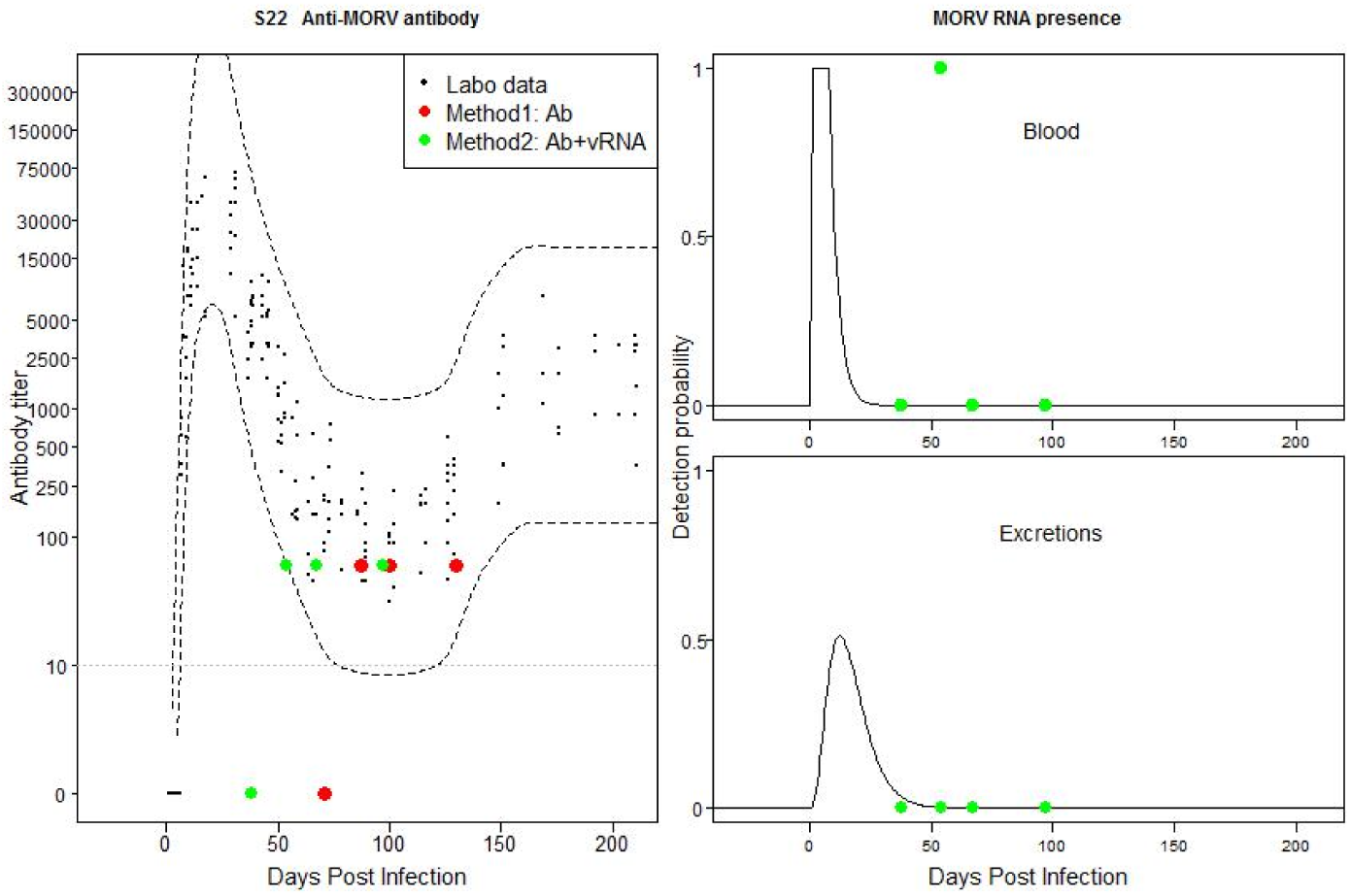
Individual 370F5: Mismatch between field and laboratory data based on both TOI method estimations. Negative Ab titer could be low titer. Possible flare-ups of vRNA in blood.

**Fig S23:**
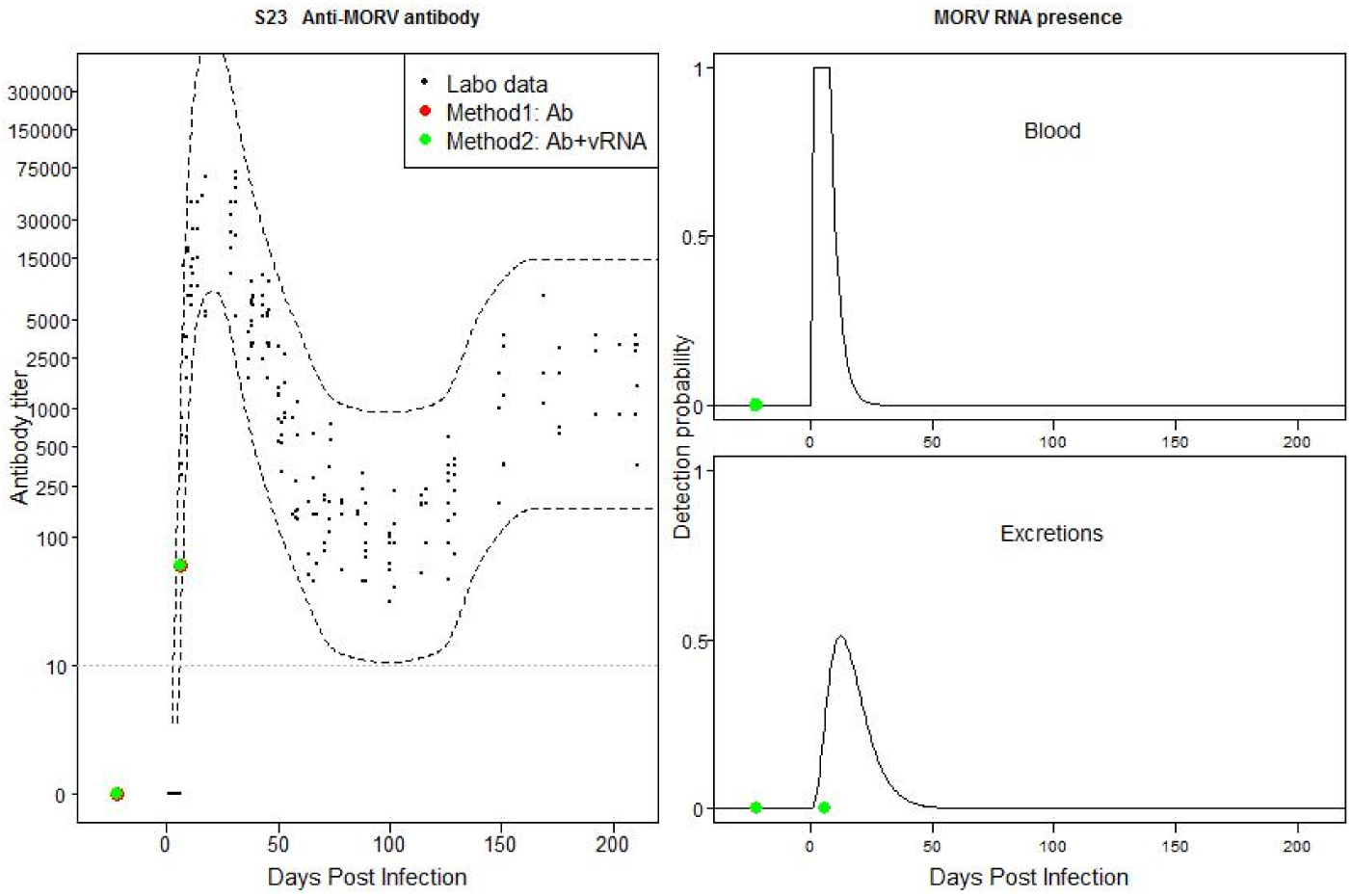
Individual 630F5: Match between field and laboratory data. Possible recently acquired active infection without evidence for positive vRNA sample.

**Fig S24:**
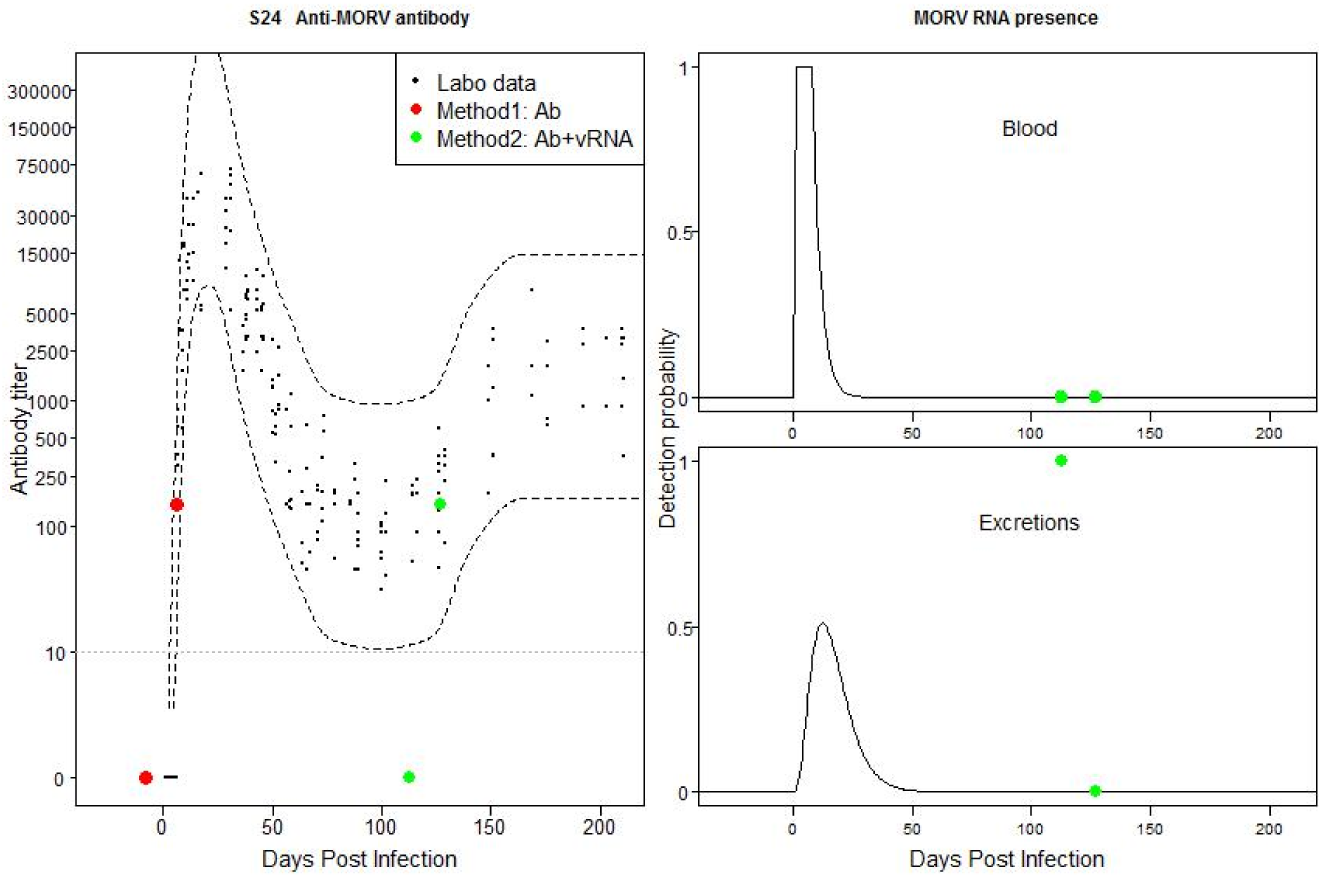
Individual 3090F5: Mismatch between field and laboratory data based on method 2. Possible recently acquired active infection. Ab titer falls just out of CB of the laboratory data.

**Fig S25:**
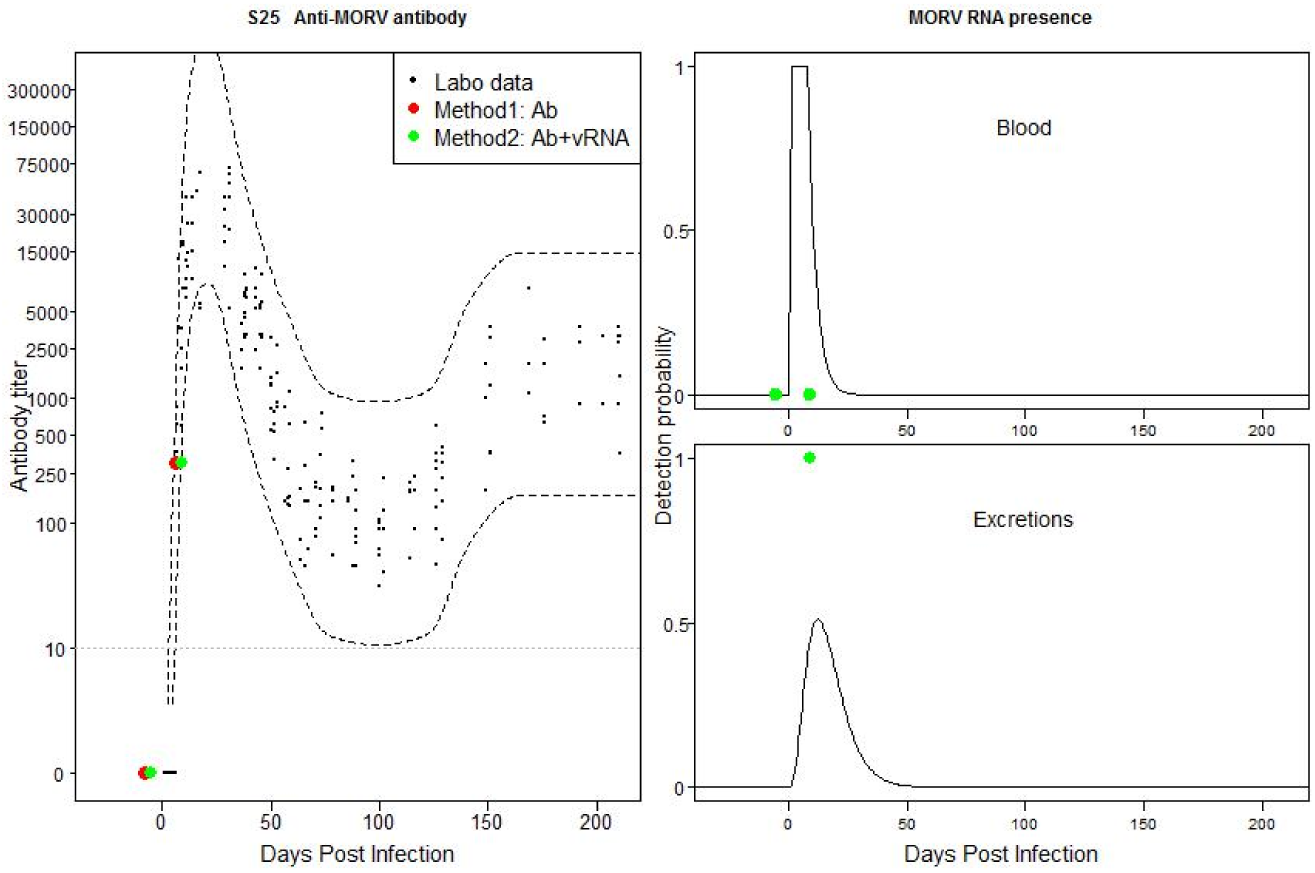
Individual 4090F5: Match between field and laboratory data. Possible recently acquired active infection.

**Fig S26:**
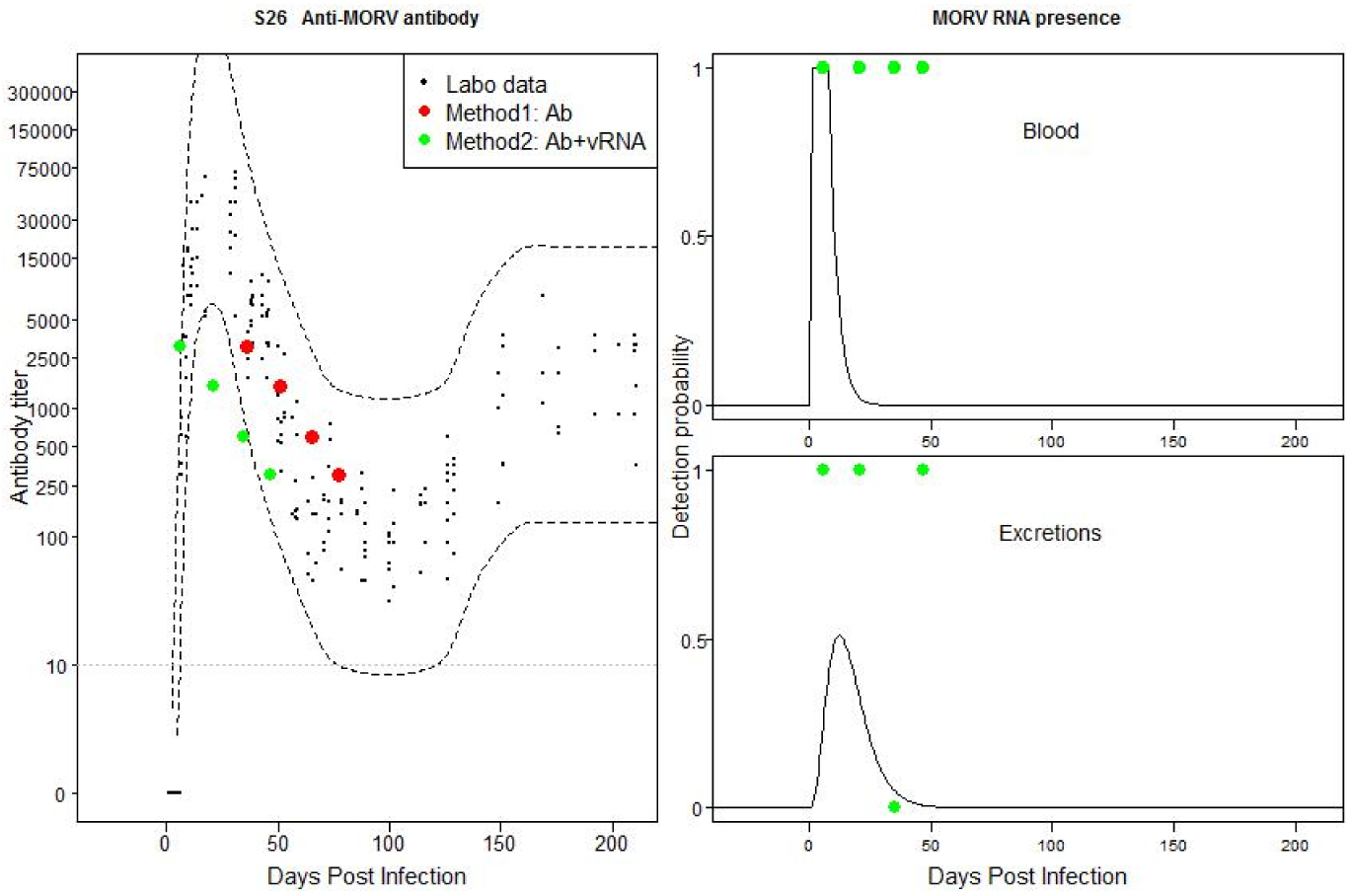
Individual 38F6: Mismatch between field and laboratory data based on method 2. Possible recently acquired chronic infection.

**Fig S27:**
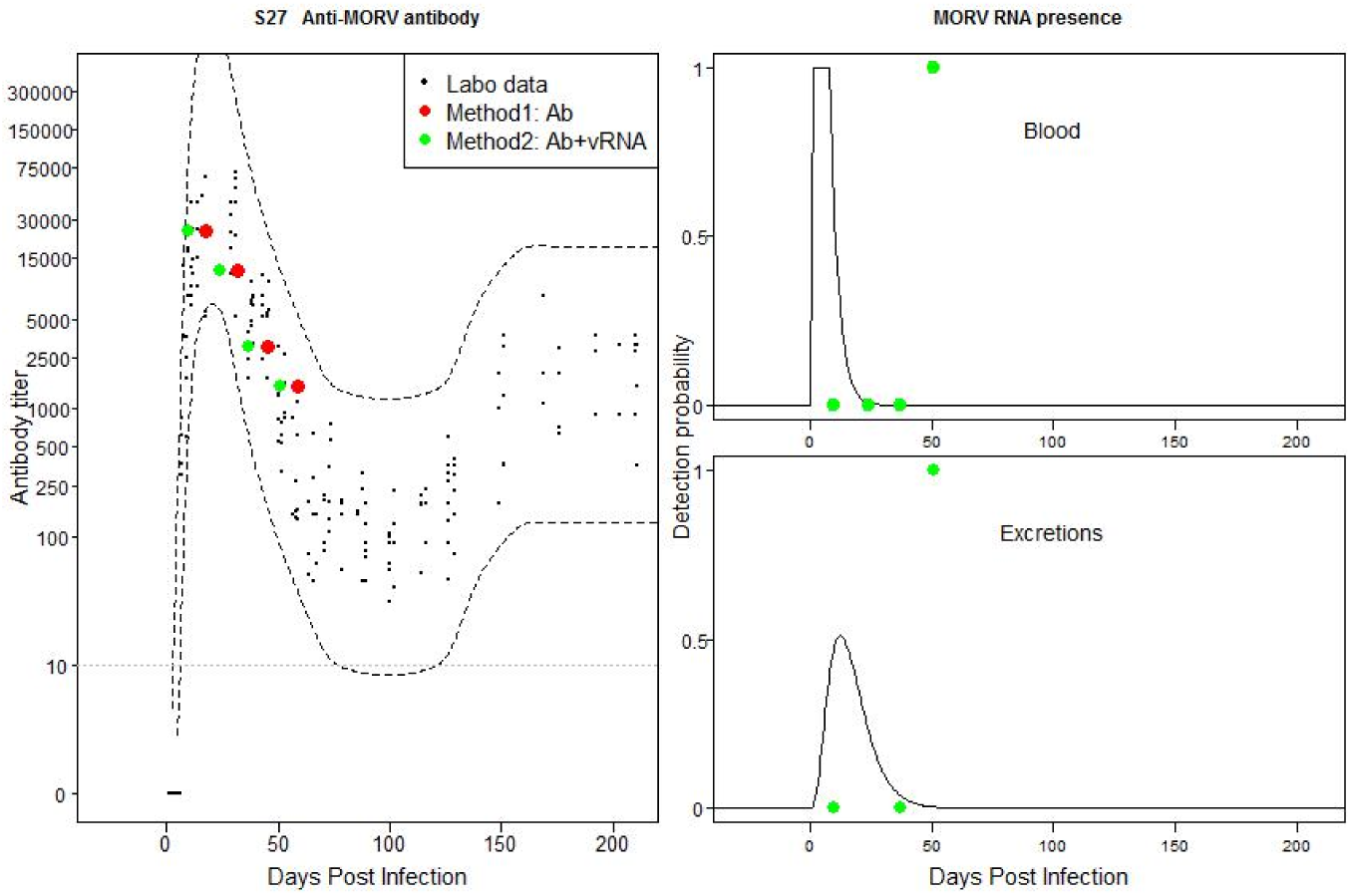
Individual 48F6: Mismatch between field and laboratory data based on method 2. Possible recently acquired infection with temporally flare-ups of MORV in blood and excretions.

**Fig S28:**
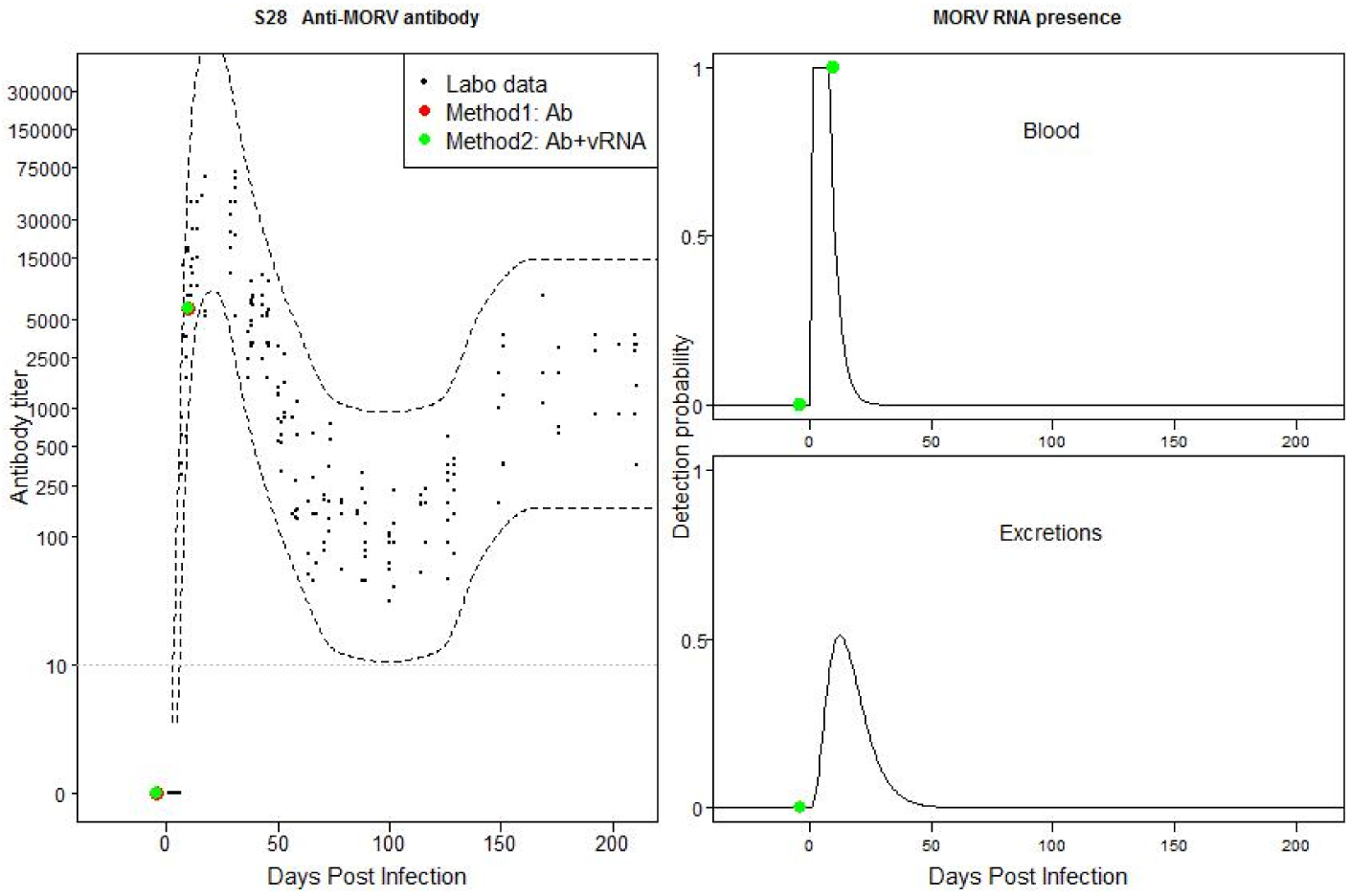
Individual 180F6: Match between field and laboratory data. Possible recently acquired active infection.

**Fig S29:**
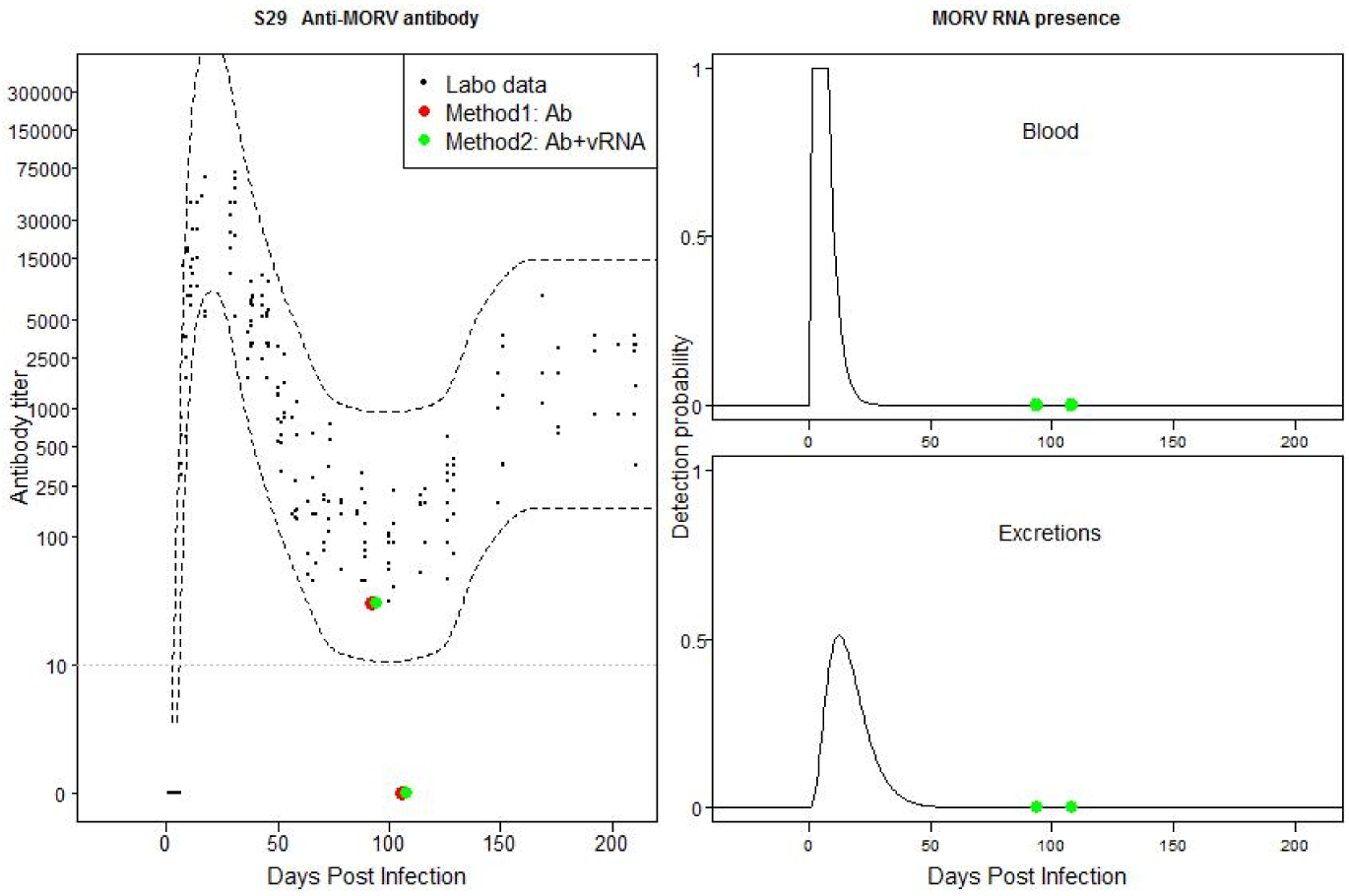
Individual 210F6: Mismatch between field and laboratory data based on both TOI method estimations. Possible past, old acute infection. Negative Ab titer could be low titer. Because the animal is still young (weight at first capture is 19g), the Ab titer pattern might be explained by the presence of maternal Abs.

**Fig S30:**
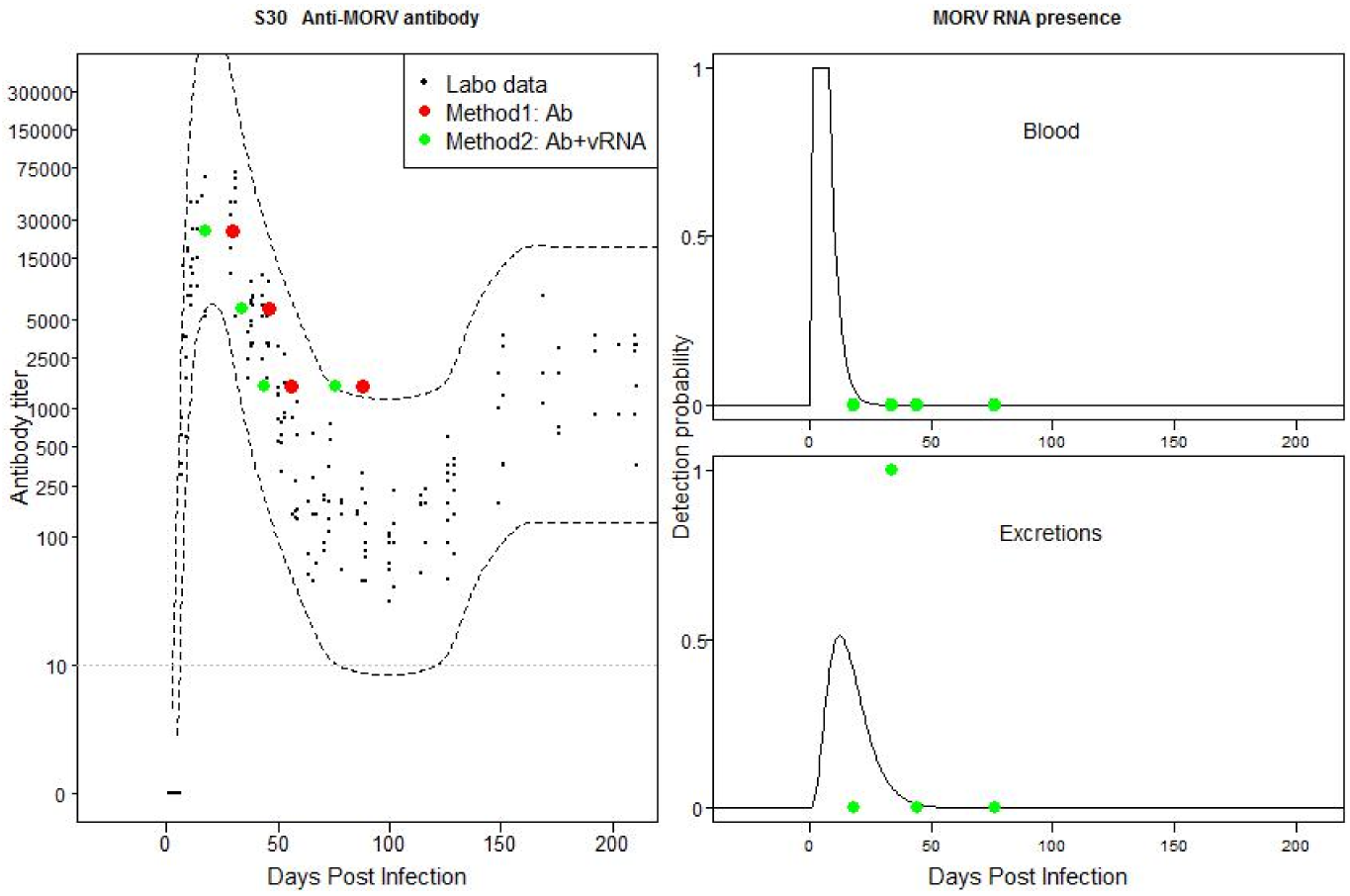
Individual 340F6: Match between field and laboratory data. Possible recently acquired acute infection.

**Fig S31:**
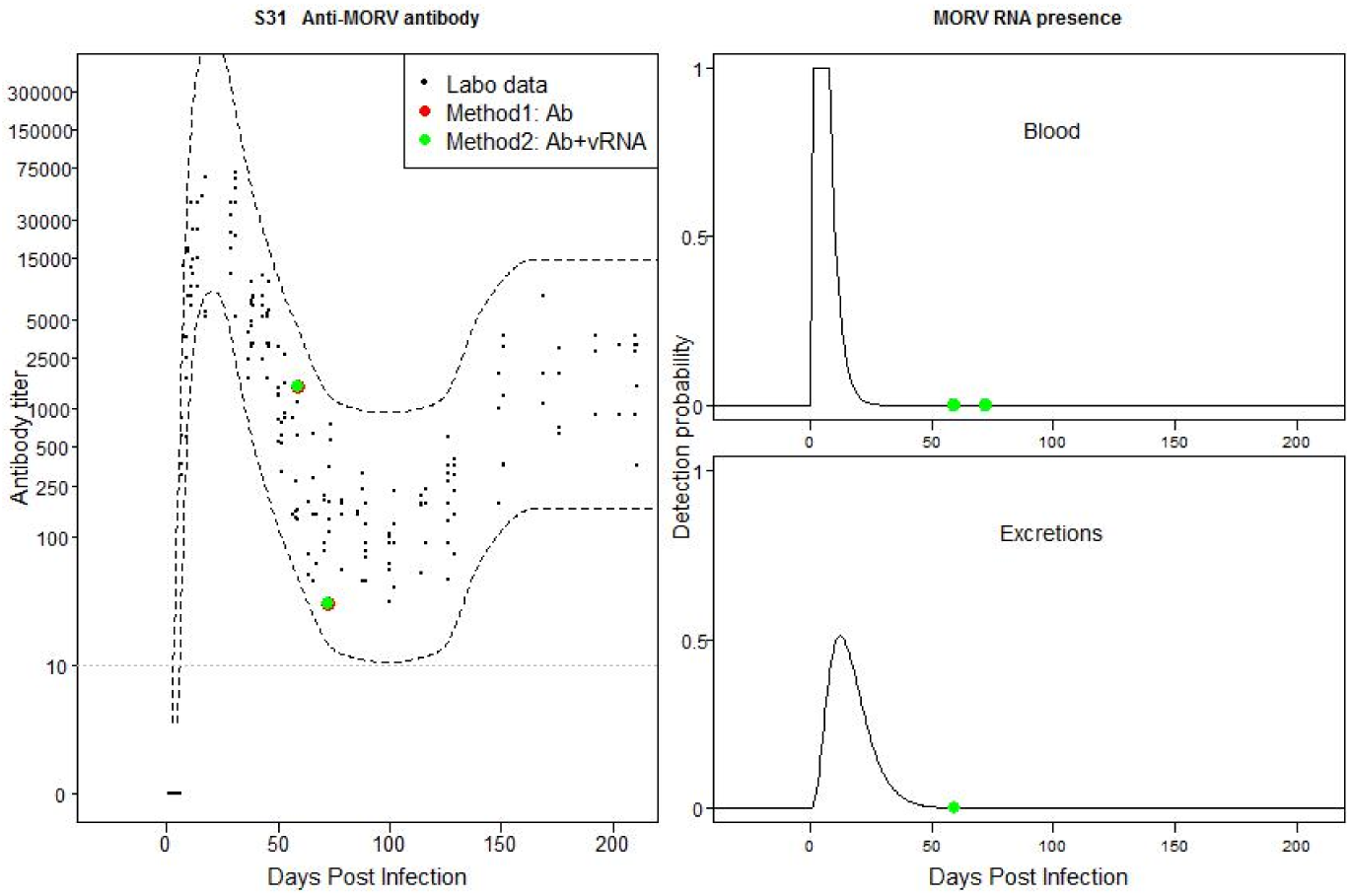
Individual 420F6: Mismatch between field and laboratory data based on both TOI methods.

**Fig S32:**
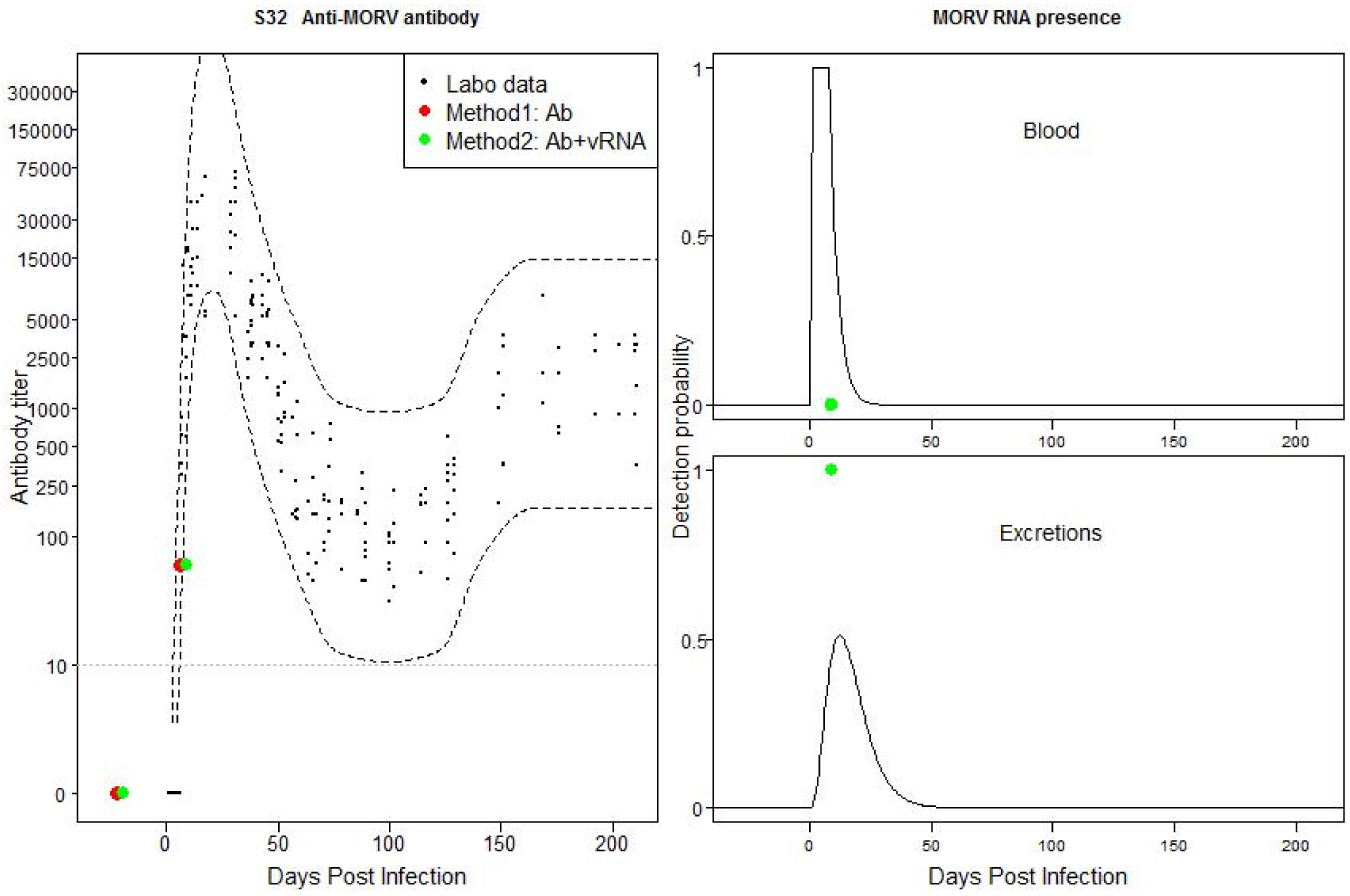
Individual 670F6: Mismatch between field and laboratory data (Ab titer is too low). Possible recently acquired active infection.

**Fig S33:**
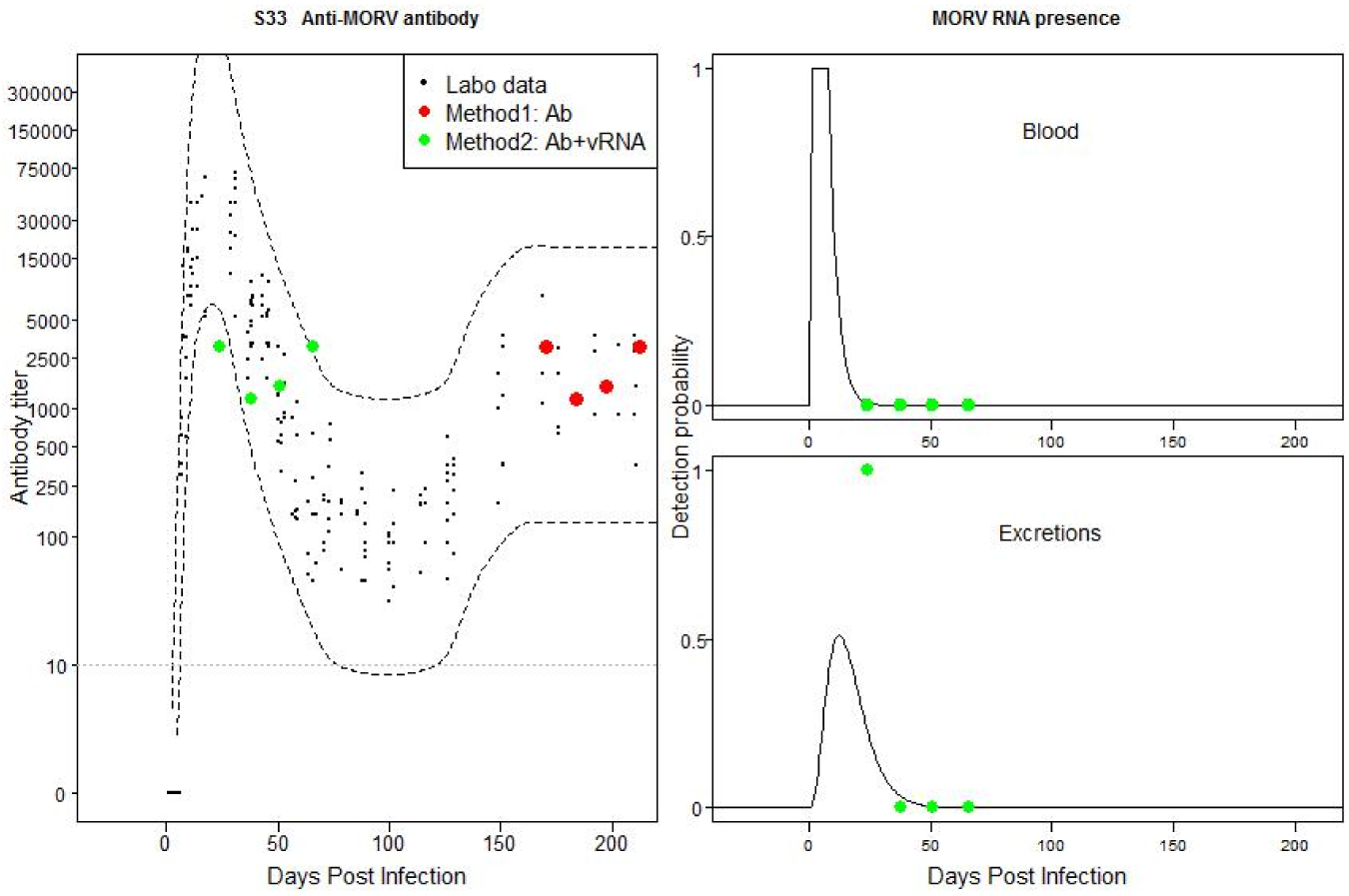
Individual 760F6: Mismatch between field and laboratory data based on method 2. Possible recently acquired active infection with Ab titers that just falls out of the CB of the laboratory data.

**Fig S34:**
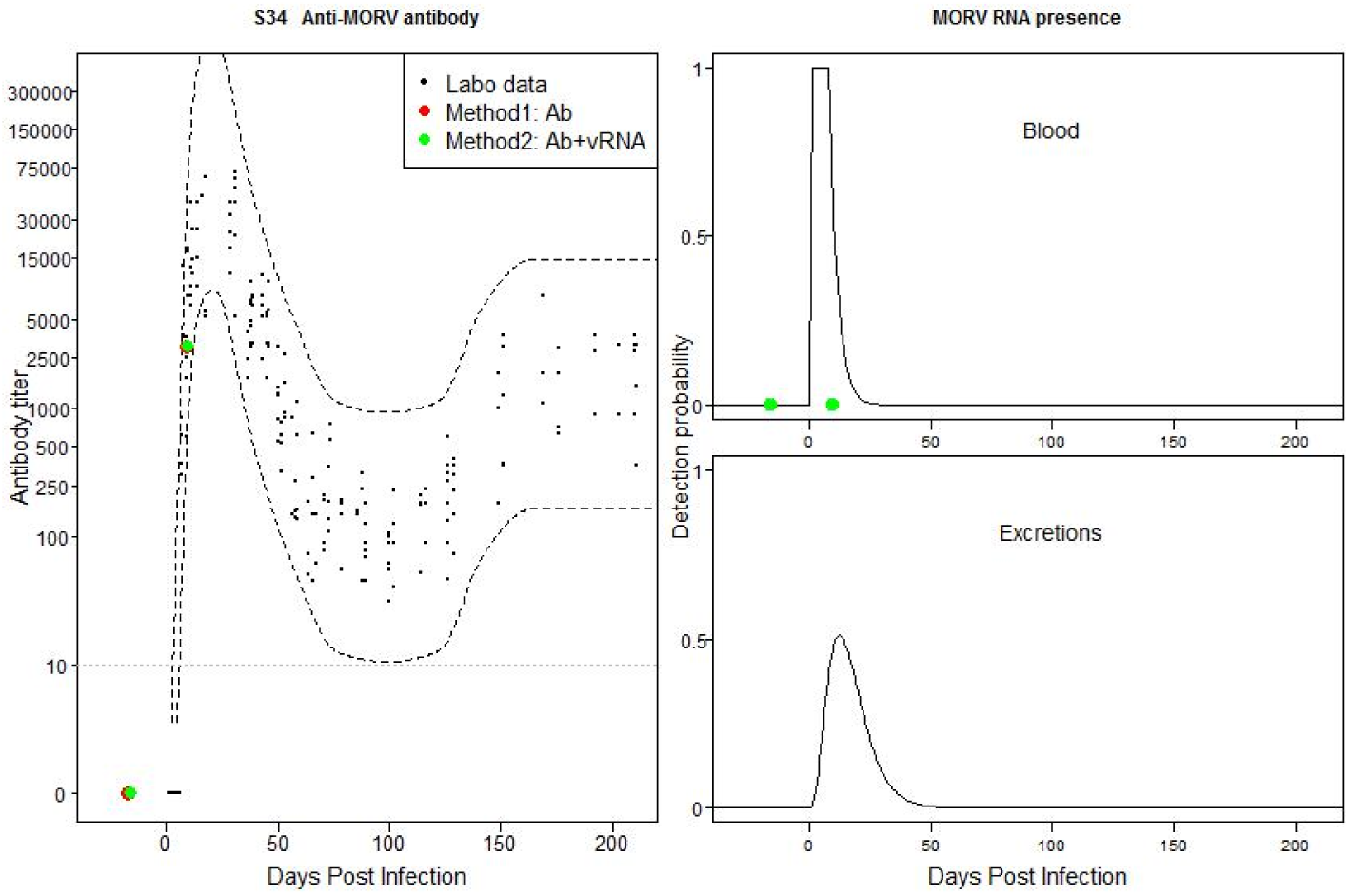
Individual 880F6: Match between field and laboratory data. Possible recently acquired active infection without evidence for positive vRNA sample.

**Fig S35:**
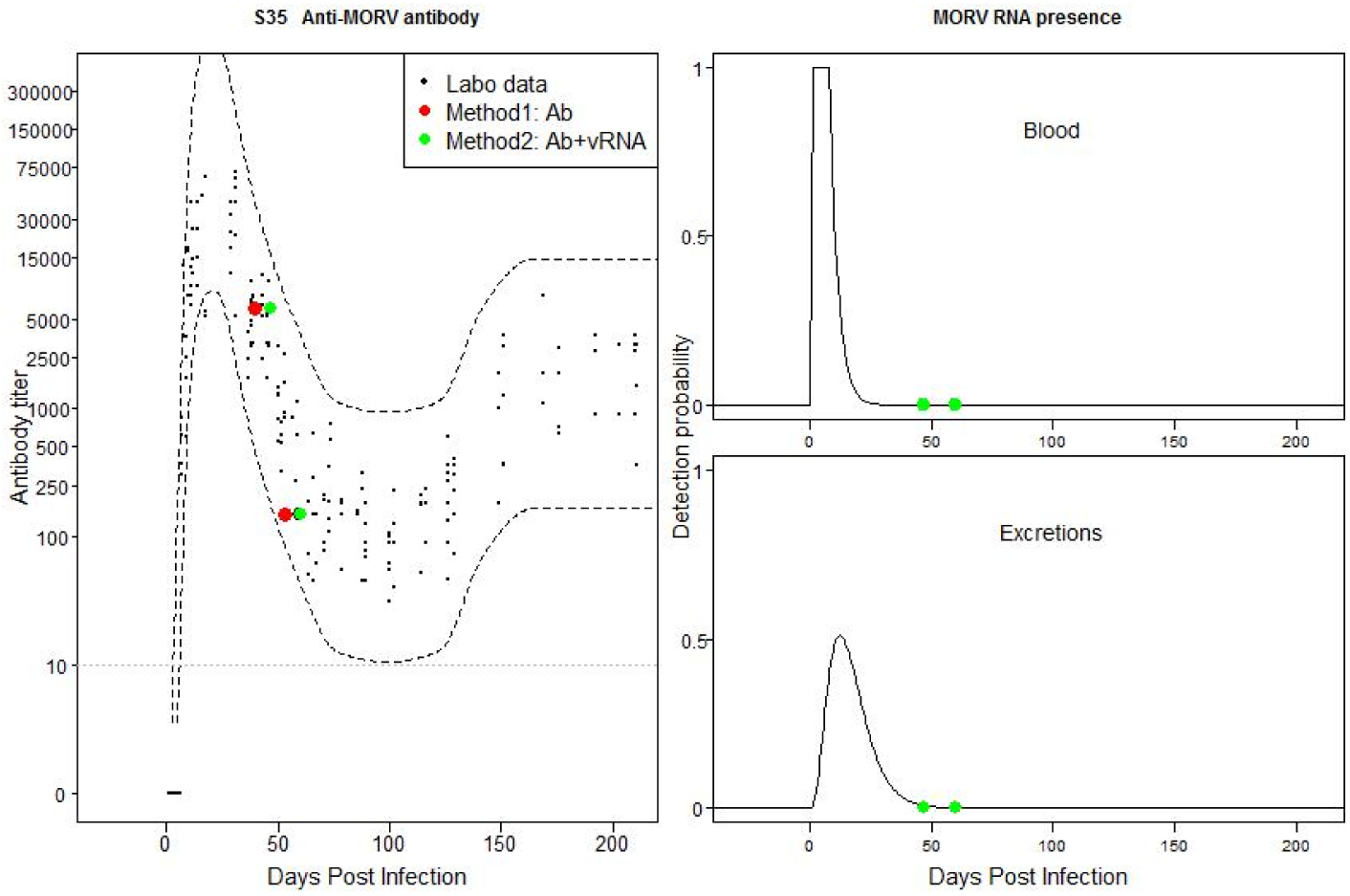
Individual 1770F6: Match between field and laboratory data. Possible past, old acute infection.

**Fig S36:**
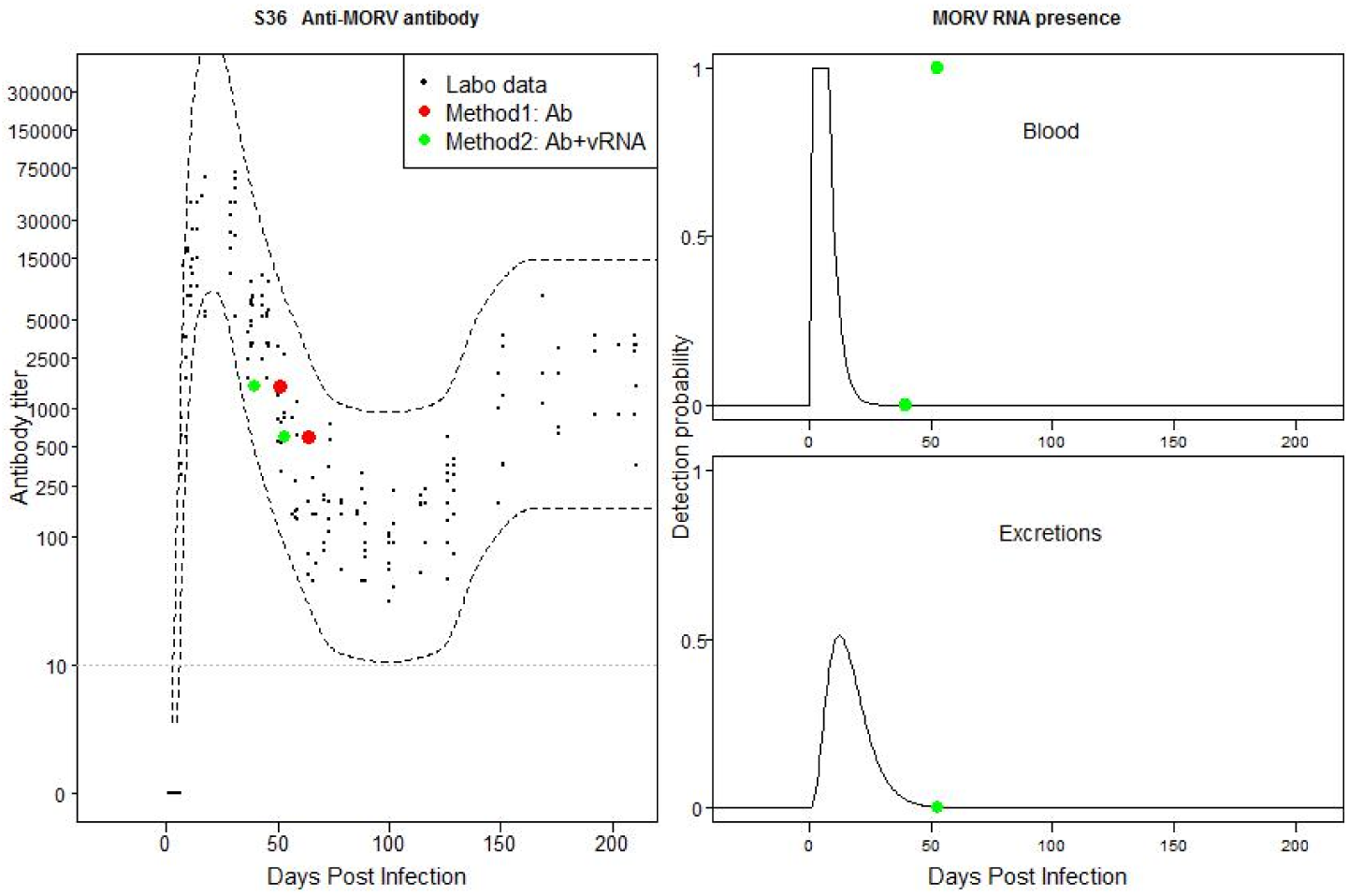
Individual 2680F6: Mismatch between field and laboratory data based on method 2. Possible old infection with temporally flare-ups of MORV in blood and excretions.

**Fig S37:**
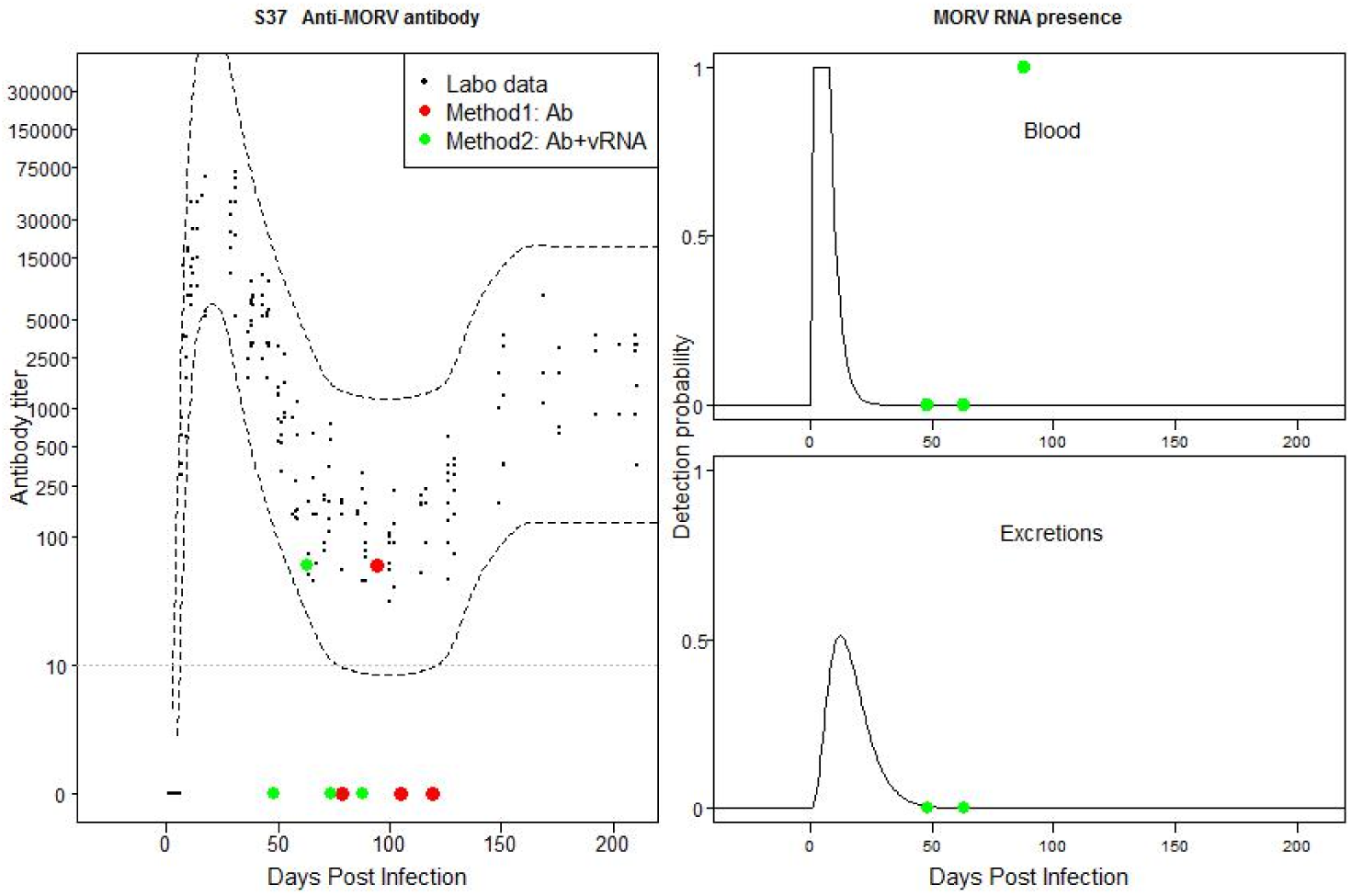
Individual 3100F6: Mismatch between field and laboratory data based on all TOI method estimations. Negative Ab titer could be low titer. Possible flare-ups of vRNA in blood.

**Fig S38:**
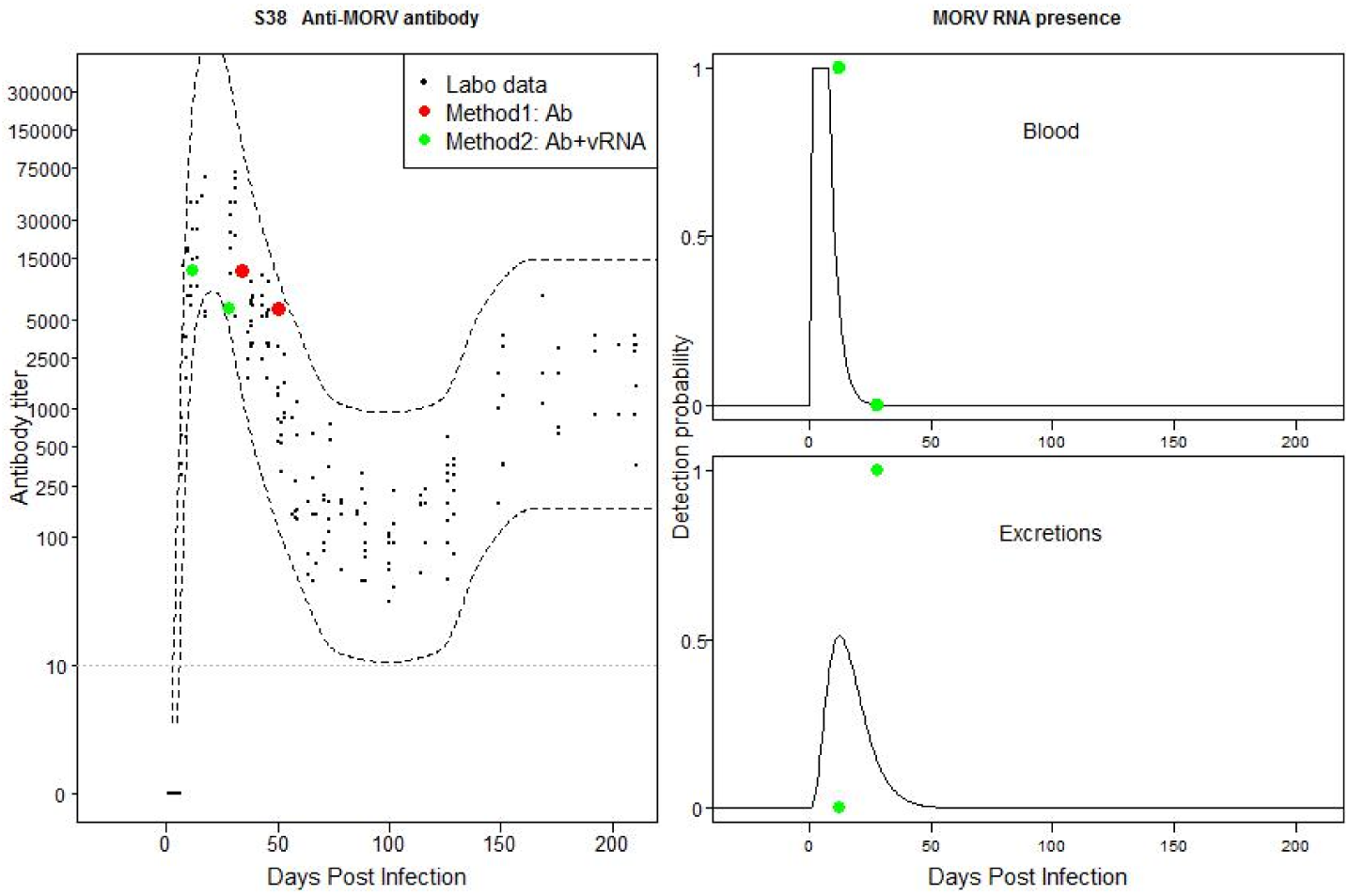
Individual 4530F6: Match between field and laboratory data. Possible recently acquired active infection.

**Fig S39:**
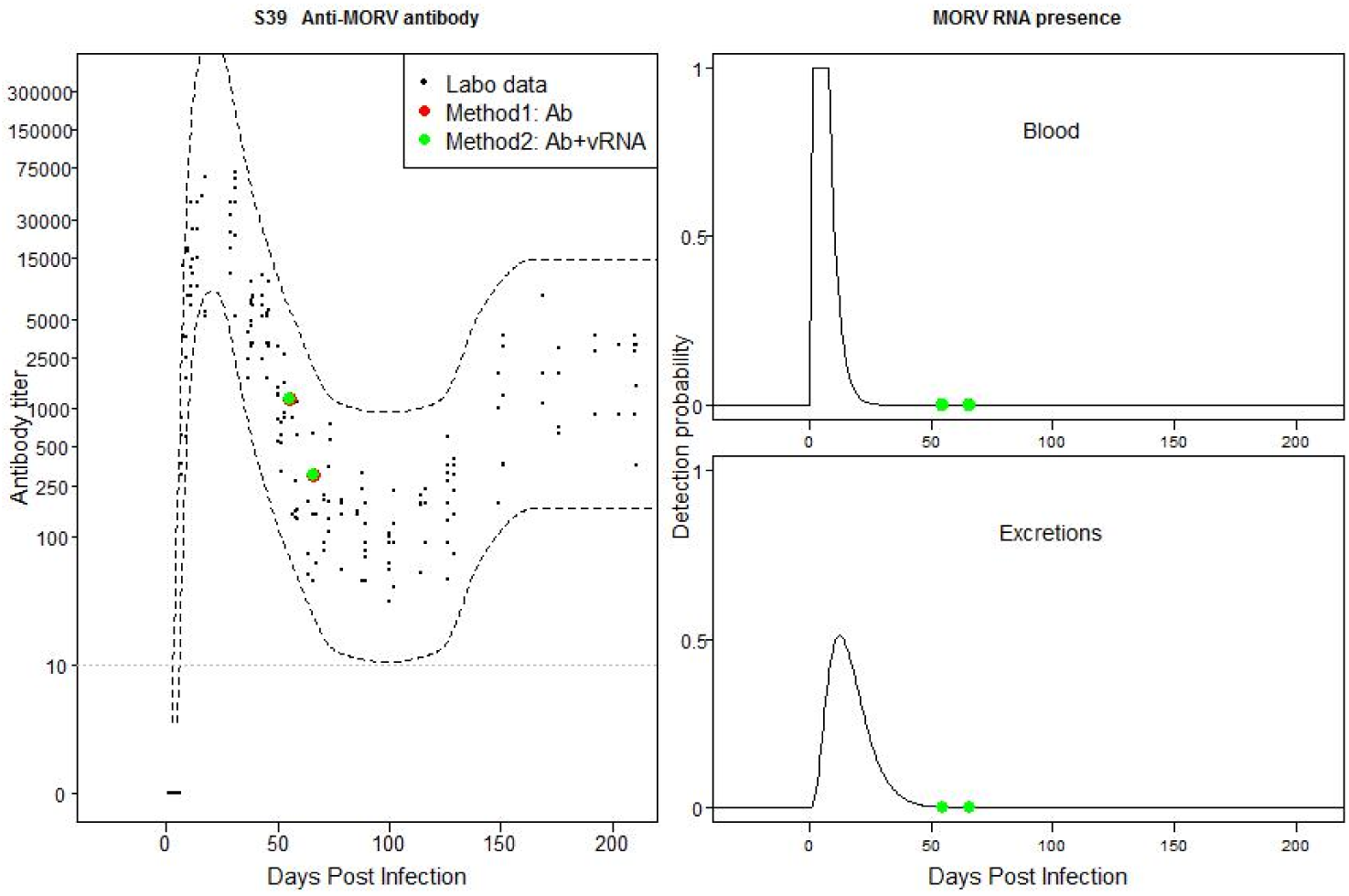
Individual 8100F6: Match between field and laboratory data. Possible past, old acute infection.

**Fig S40:**
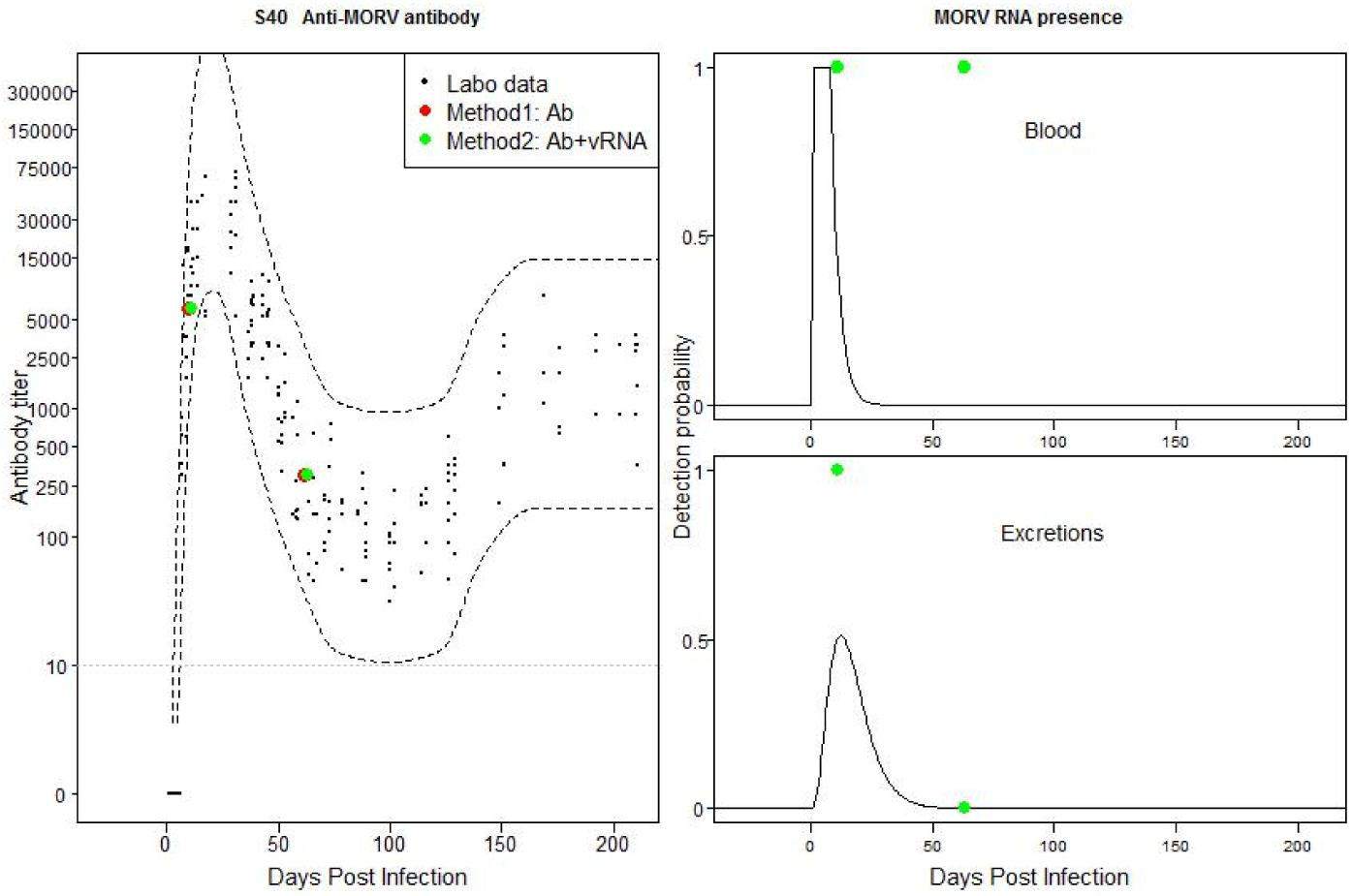
Individual 11090F6: Mismatch between field and laboratory data based on method 2. Possible recently acquired active infection. Possible recently acquired infection with temporally flare-ups of MORV in blood.

**Fig S41:**
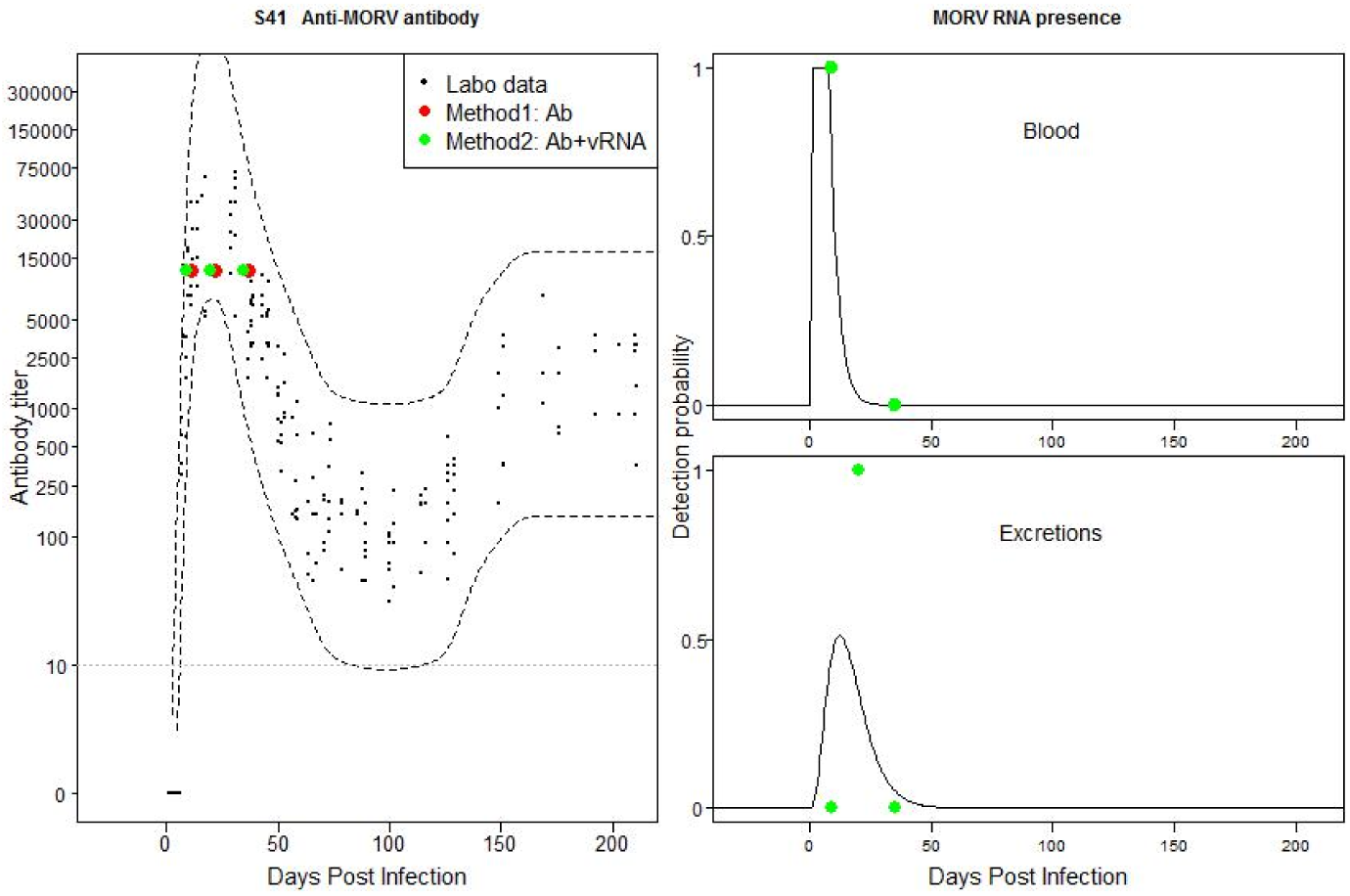
Individual 30100F6: Mismatch between field and laboratory data based on both TOI estimations. Possible recently acquired active infection, but first Ab titer falls out of CB.

**Fig S42:**
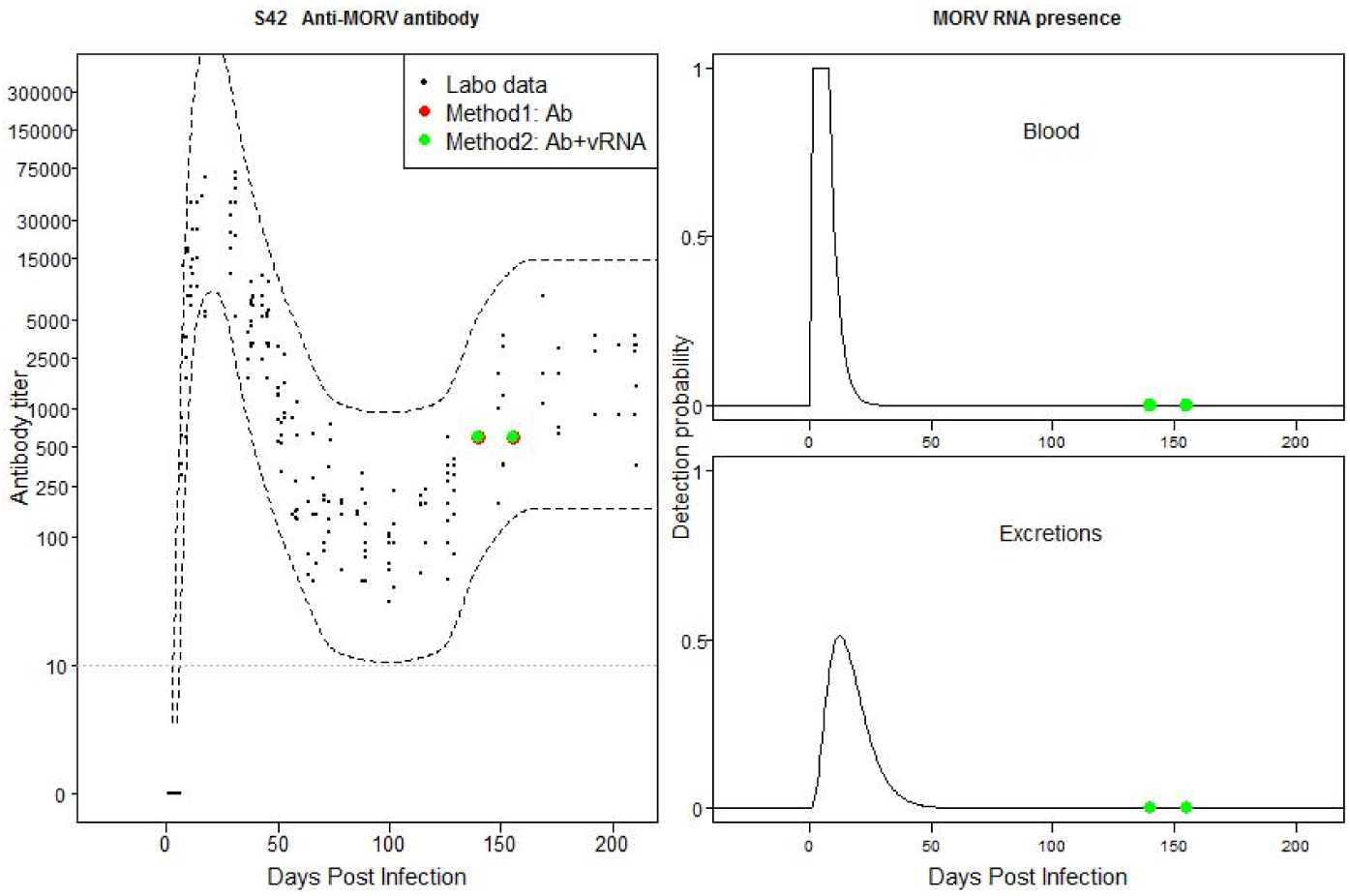
Individual 110100F6: Match between field and laboratory data. Possible past, old acute infection.

## Supplementary Tables

**Table 2:**
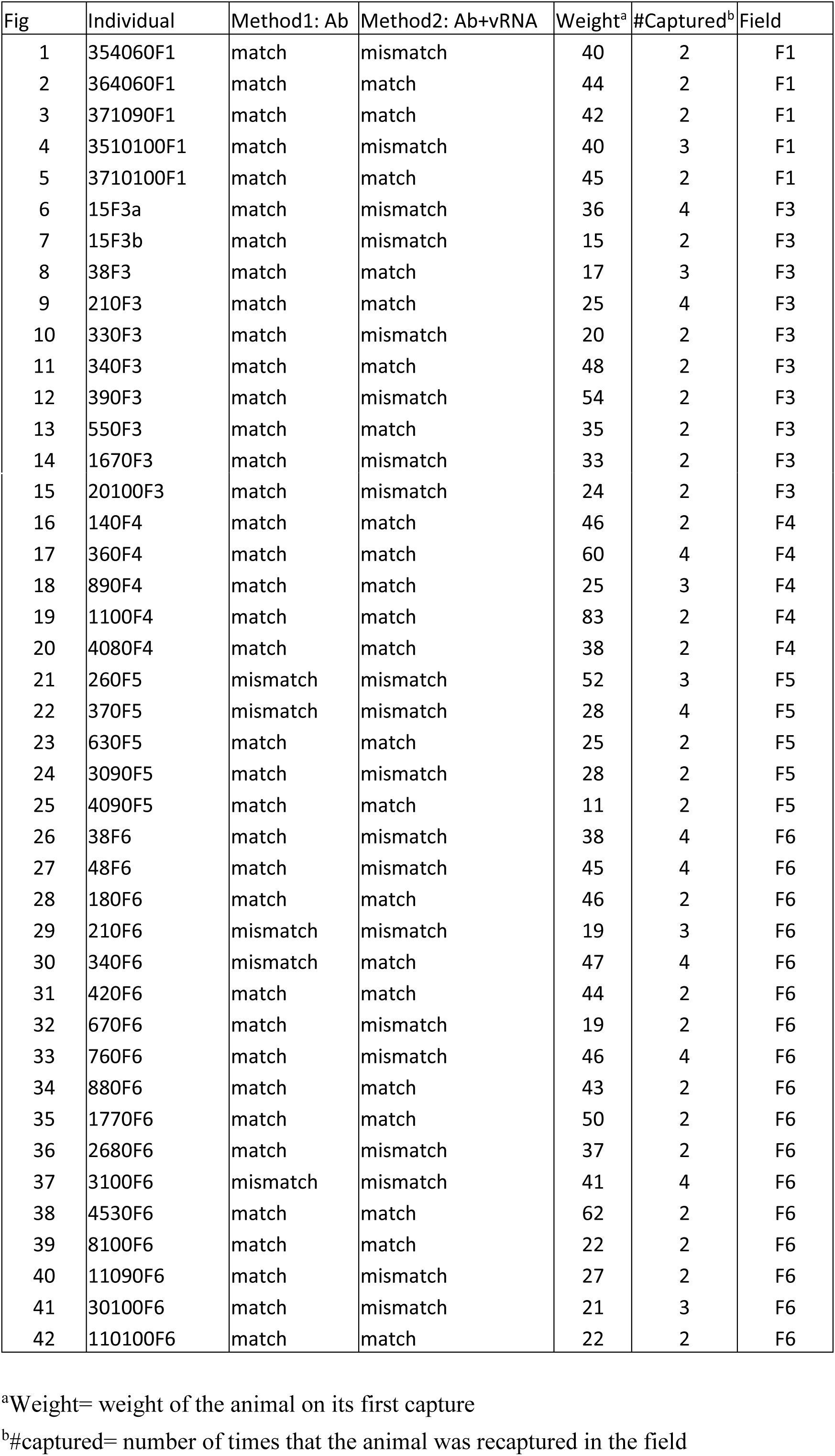
Comparison of the Ab and vRNA patterns between field and laboratory mice for each recaptured Ab-positive mouse and both TOI methods

**Table 3:** Comparison of Morogoro virus (MORV) antibody (Ab) titers between field and laboratory mice. For each field data point the difference with the confidence band (CB) of the laboratory data was given. ‘CB’ indicates that the field data point falls within the CB. Time of infection (TOI) was based on method 1 (only Ab titer).

| Individual | TOI | Ab titers | Day capture | Difference with CB |
| --- | --- | --- | --- | --- |
| 354060F1 | -34 | 0 | 0 | Not possible |
| 354060F1 | -34 | 60 | 40 | CB |
| 364060F1 | 92 | 30 | 0 | CB |
| 364060F1 | 92 | 300 | 41 | CB |
| 371090F1 | 9 | 3000 | 0 | CB |
| 371090F1 | 9 | 12000 | 12 | CB |
| 3510100F1 | 141 | 600 | 0 | CB |
| 3510100F1 | 141 | 3000 | 14 | CB |
| 3510100F1 | 141 | 6000 | 29 | CB |
| 3710100F1 | -2 | 0 | 0 | Not possible |
| 3710100F1 | -2 | 12000 | 14 | CB |
| 15F3a | 170 | 6000 | 0 | CB |
| 15F3a | 170 | 3000 | 27 | CB |
| 15F3a | 170 | 3000 | 41 | CB |
| 15F3a | 170 | 3000 | 53 | CB |
| 15F3b | 11 | 12000 | 0 | CB |
| 15F3b | 11 | 6000 | 32 | CB |
| 38F3 | -50 | 0 | 0 | Not possible |
| 38F3 | -50 | 0 | 27 | Not possible |
| 38F3 | -50 | 60 | 56 | CB |
| 210F3 | -47 | 0 | 0 | Not possible |
| 210F3 | -47 | 0 | 29 | Not possible |
| 210F3 | -47 | 0 | 33 | Not possible |
| 210F3 | -47 | 6000 | 57 | CB |
| 330F3 | 13 | 24000 | 0 | CB |
| 330F3 | 13 | 1500 | 56 | CB |
| 340F3 | -19 | 0 | 0 | Not possible |
| 340F3 | -19 | 6000 | 29 | CB |
| 390F3 | -33 | 0 | 0 | Not possible |
| 390F3 | -33 | 150 | 39 | CB |
| 550F3 | 12 | 12000 | 0 | CB |
| 550F3 | 12 | 18000 | 17 | CB |
| 1670F3 | 39 | 3000 | 0 | CB |
| 1670F3 | 39 | 1500 | 13 | CB |
| 20100F3 | -22 | 0 | 0 | Not possible |
| 20100F3 | -22 | 30 | 28 | CB |
| 140F4 | -36 | 0 | 0 | Not possible |
| 140F4 | -36 | 30 | 42 | CB |
| 360F4 | 170 | 12000 | 0 | CB |
| 360F4 | 170 | 3000 | 16 | CB |
| 360F4 | 170 | 3000 | 29 | CB |
| 360F4 | 170 | 3000 | 59 | CB |
| 890F4 | 24 | 12000 | 0 | CB |
| 890F4 | 24 | 6000 | 14 | CB |
| 890F4 | 24 | 3000 | 26 | CB |
| 1100F4 | 133 | 300 | 0 | CB |
| 1100F4 | 133 | 1200 | 14 | CB |
| 4080F4 | 170 | 3000 | 0 | CB |
| 4080F4 | 170 | 3000 | 14 | CB |
| 260F5 | 84 | 30 | 0 | CB |
| 260F5 | 84 | 0 | 14 | 2.23 |
| 260F5 | 84 | 0 | 27 | 2.26 |
| 370F5 | 71 | 1.1 | 0 | 2.72 |
| 370F5 | 71 | 60 | 16 | CB |
| 370F5 | 71 | 60 | 29 | CB |
| 370F5 | 71 | 60 | 59 | CB |
| 630F5 | -22 | 0 | 0 | Not possible |
| 630F5 | -22 | 60 | 28 | CB |
| 3090F5 | -8 | 0 | 0 | Not possible |
| 3090F5 | -8 | 150 | 14 | CB |
| 4090F5 | -8 | 0 | 0 | Not possible |
| 4090F5 | -8 | 300 | 14 | CB |
| 38F6 | 36 | 3000 | 0 | CB |
| 38F6 | 36 | 1500 | 15 | CB |
| 38F6 | 36 | 600 | 29 | CB |
| 38F6 | 36 | 300 | 41 | CB |
| 48F6 | 18 | 24000 | 0 | CB |
| 48F6 | 18 | 12000 | 14 | CB |
| 48F6 | 18 | 3000 | 27 | CB |
| 48F6 | 18 | 1500 | 41 | CB |
| 180F6 | -4 | 0 | 0 | Not possible |
| 180F6 | -4 | 6000 | 14 | CB |
| 210F6 | 92 | 30 | 0 | CB |
| 210F6 | 92 | 0 | 14 | 2.37 |
| 340F6 | 30 | 24000 | 0 | CB |
| 340F6 | 30 | 6000 | 16 | CB |
| 340F6 | 30 | 1500 | 26 | CB |
| 340F6 | 30 | 1500 | 58 | 0.16 |
| 420F6 | 59 | 1500 | 0 | CB |
| 420F6 | 59 | 30 | 13 | CB |
| 670F6 | -22 | 0 | 0 | Not possible |
| 670F6 | -22 | 60 | 28 | CB |
| 760F6 | 170 | 3000 | 0 | CB |
| 760F6 | 170 | 1200 | 14 | CB |
| 760F6 | 170 | 1500 | 27 | CB |
| 760F6 | 170 | 3000 | 42 | CB |
| 880F6 | -17 | 0 | 0 | Not possible |
| 880F6 | -17 | 3000 | 26 | CB |
| 1770F6 | 40 | 6000 | 0 | CB |
| 1770F6 | 40 | 150 | 13 | CB |
| 2680F6 | 51 | 1500 | 0 | CB |
| 2680F6 | 51 | 600 | 13 | CB |
| 3100F6 | 79 | 1.1 | 0 | 2.23 |
| 3100F6 | 79 | 60 | 15 | CB |
| 3100F6 | 79 | 0 | 26 | 2.13 |
| 3100F6 | 79 | 0 | 40 | 2.26 |
| 4530F6 | 34 | 12000 | 0 | CB |
| 4530F6 | 34 | 6000 | 16 | CB |
| 8100F6 | 55 | 1200 | 0 | CB |
| 8100F6 | 55 | 300 | 11 | CB |
| 11090F6 | 10 | 6000 | 0 | CB |
| 11090F6 | 10 | 300 | 52 | CB |
| 30100F6 | 11 | 12000 | 0 | CB |
| 30100F6 | 11 | 12000 | 11 | CB |
| 30100F6 | 11 | 12000 | 26 | CB |
| 110100F6 | 140 | 600 | 0 | CB |
| 110100F6 | 140 | 600 | 15 | CB |

**Table 4:** Comparison of Morogoro virus (MORV) antibody (Ab) titer and vRNA presence/absence between field and laboratory mice. For each field data point the difference with the confidence band (CB) of the laboratory was given. ‘CB’ indicates that the field data point falls within the CB. Time of infection (TOI) was based on method 2 (both Ab and vRNA titer in blood and urine).

| Individual | TOI | Ab titers | MORV RNA blood | MORV RNA excretions | Day capture | Difference with CB Ab | Difference with CB blood | Difference with CB excretions |
| --- | --- | --- | --- | --- | --- | --- | --- | --- |
| 354060F1 | 5 | 0 | yes | yes | 0 | CB | CB | CB |
| 354060F1 | 5 | 60 | No | NA | 40 | 1.77 | CB | NA |
| 364060F1 | 6 | 30 | yes | No | 0 | CB | CB | CB |
| 364060F1 | 6 | 300 | No | No | 41 | CB | CB | CB |
| 371090F1 | 9 | 3000 | yes | No | 0 | CB | CB | CB |
| 371090F1 | 9 | 12000 | No | No | 12 | CB | CB | CB |
| 3510100F1 | 7 | 600 | yes | No | 0 | CB | CB | CB |
| 3510100F1 | 7 | 3000 | No | No | 14 | 0.77 | CB | CB |
| 3510100F1 | 7 | 6000 | No | No | 29 | CB | CB | CB |
| 3710100F1 | -2 | 0 | No | No | 0 | Not possible | Not possible | Not possible |
| 3710100F1 | -2 | 12000 | NA | NA | 14 | CB | NA | NA |
| 15F3a | 10 | 6000 | No | No | 0 | CB | CB | CB |
| 15F3a | 10 | 3000 | yes | yes | 27 | CB | 0.01 | CB |
| 15F3a | 10 | 3000 | No | No | 41 | CB | CB | CB |
| 15F3a | 10 | 3000 | No | No | 53 | CB | CB | CB |
| 15F3b | 10 | 12000 | NA | No | 0 | CB | NA | CB |
| 15F3b | 10 | 6000 | yes | No | 32 | CB | 0.01 | CB |
| 38F3 | -50 | 0 | No | No | 0 | Not possible | Not possible | Not possible |
| 38F3 | -50 | 0 | No | No | 27 | Not possible | Not possible | Not possible |
| 38F3 | -50 | 60 | yes | No | 56 | CB | CB | CB |
| 210F3 | -46 | 0 | No | No | 0 | Not possible | Not possible | Not possible |
| 210F3 | -46 | 0 | No | No | 29 | Not possible | Not possible | Not possible |
| 210F3 | -46 | 0 | No | No | 33 | Not possible | Not possible | Not possible |
| 210F3 | -46 | 6000 | No | No | 57 | CB | CB | CB |
| 330F3 | 13 | 24000 | No | No | 0 | CB | CB | CB |
| 330F3 | 13 | 1500 | yes | yes | 56 | CB | 0.03 | 0.03 |
| 340F3 | -19 | 0 | NA | NA | 0 | Not possible | NA | NA |
| 340F3 | -19 | 6000 | NA | No | 29 | CB | NA | CB |
| 390F3 | 88 | 0 | No | yes | 0 | 2.43 | CB | 0.03 |
| 390F3 | 88 | 150 | No | yes | 39 | CB | CB | 0.03 |
| 550F3 | 14 | 12000 | NA | No | 0 | CB | NA | CB |
| 550F3 | 14 | 18000 | NA | No | 17 | CB | NA | CB |
| 1670F3 | 9 | 3000 | yes | No | 0 | CB | CB | CB |
| 1670F3 | 9 | 1500 | yes | yes | 13 | 1.63 | CB | CB |
| 20100F3 | 5 | 0 | yes | No | 0 | CB | CB | CB |
| 20100F3 | 5 | 30 | yes | yes | 28 | 4.04 | 0.02 | CB |
| 140F4 | -36 | 0 | No | NA | 0 | Not possible | Not possible | NA |
| 140F4 | -36 | 30 | yes | No | 42 | CB | CB | CB |
| 360F4 | 170 | 12000 | NA | No | 0 | CB | NA | CB |
| 360F4 | 170 | 3000 | NA | No | 16 | CB | NA | CB |
| 360F4 | 170 | 3000 | No | No | 29 | CB | CB | CB |
| 360F4 | 170 | 3000 | NA | NA | 59 | CB | NA | NA |
| 890F4 | 32 | 12000 | No | NA | 0 | CB | CB | NA |
| 890F4 | 32 | 6000 | No | No | 14 | CB | CB | CB |
| 890F4 | 32 | 3000 | No | No | 26 | CB | CB | CB |
| 1100F4 | 133 | 300 | No | No | 0 | CB | CB | CB |
| 1100F4 | 133 | 1200 | NA | No | 14 | CB | NA | CB |
| 4080F4 | 170 | 3000 | No | No | 0 | CB | CB | CB |
| 4080F4 | 170 | 3000 | No | No | 14 | CB | CB | CB |
| 260F5 | 60 | 30 | NA | No | 0 | Not possible | NA | CB |
| 260F5 | 60 | 0 | yes | No | 14 | Not possible | 0.02 | CB |
| 260F5 | 60 | 0 | No | NA | 27 | Not possible | CB | NA |
| 370F5 | 38 | 1.1 | No | No | 0 | 6.28 | CB | CB |
| 370F5 | 38 | 60 | yes | No | 16 | 0.52 | 0.01 | CB |
| 370F5 | 38 | 60 | No | No | 29 | CB | CB | CB |
| 370F5 | 38 | 60 | No | No | 59 | CB | CB | CB |
| 630F5 | -22 | 0 | No | No | 0 | Not possible | Not possible | Not possible |
| 630F5 | -22 | 60 | NA | No | 28 | CB | NA | CB |
| 3090F5 | 113 | 0 | No | yes | 0 | 2.42 | CB | 0.03 |
| 3090F5 | 113 | 150 | No | No | 14 | CB | CB | CB |
| 4090F5 | -5 | 0 | No | NA | 0 | Not possible | Not possible | NA |
| 4090F5 | -5 | 300 | No | yes | 14 | CB | CB | CB |
| 38F6 | 6 | 3000 | yes | yes | 0 | 0.93 | CB | CB |
| 38F6 | 6 | 1500 | yes | yes | 15 | 1.36 | CB | CB |
| 38F6 | 6 | 600 | yes | No | 29 | 0.52 | 0.01 | CB |
| 38F6 | 6 | 300 | yes | yes | 41 | CB | 0.01 | 0.01 |
| 48F6 | 10 | 24000 | No | No | 0 | CB | CB | CB |
| 48F6 | 10 | 12000 | No | NA | 14 | CB | CB | NA |
| 48F6 | 10 | 3000 | No | No | 27 | CB | CB | CB |
| 48F6 | 10 | 1500 | yes | yes | 41 | CB | 0.01 | 0.01 |
| 180F6 | -4 | 0 | No | No | 0 | Not possible | Not possible | Not possible |
| 180F6 | -4 | 6000 | yes | NA | 14 | CB | CB | NA |
| 210F6 | 94 | 30 | No | No | 0 | CB | CB | CB |
| 210F6 | 94 | 0 | No | No | 14 | 2.38 | CB | CB |
| 340F6 | 18 | 24000 | No | No | 0 | CB | CB | CB |
| 340F6 | 18 | 6000 | No | yes | 16 | CB | CB | CB |
| 340F6 | 18 | 1500 | No | No | 26 | CB | CB | CB |
| 340F6 | 18 | 1500 | No | No | 58 | CB | CB | CB |
| 420F6 | 59 | 1500 | No | No | 0 | CB | CB | CB |
| 420F6 | 59 | 30 | No | NA | 13 | CB | CB | NA |
| 670F6 | -19 | 0 | NA | NA | 0 | Not possible | NA | NA |
| 670F6 | -19 | 60 | No | yes | 28 | 0.47 | CB | CB |
| 760F6 | 24 | 3000 | No | yes | 0 | 0.75 | CB | CB |
| 760F6 | 24 | 1200 | No | No | 14 | CB | CB | CB |
| 760F6 | 24 | 1500 | No | No | 27 | CB | CB | CB |
| 760F6 | 24 | 3000 | No | No | 42 | CB | CB | CB |
| 880F6 | -16 | 0 | No | NA | 0 | Not possible | Not possible | NA |
| 880F6 | -16 | 3000 | No | NA | 26 | CB | CB | NA |
| 1770F6 | 47 | 6000 | No | No | 0 | CB | CB | CB |
| 1770F6 | 47 | 150 | No | No | 13 | CB | CB | CB |
| 2680F6 | 40 | 1500 | No | NA | 0 | CB | CB | NA |
| 2680F6 | 40 | 600 | yes | No | 13 | CB | 0.03 | CB |
| 3100F6 | 48 | 1.1 | No | No | 0 | 5.25 | CB | CB |
| 3100F6 | 48 | 60 | No | No | 15 | CB | CB | CB |
| 3100F6 | 48 | 0 | NA | NA | 26 | 2.53 | NA | NA |
| 3100F6 | 48 | 0 | yes | NA | 40 | 2.19 | 0.01 | NA |
| 4530F6 | 12 | 12000 | yes | No | 0 | CB | CB | CB |
| 4530F6 | 12 | 6000 | No | yes | 16 | CB | CB | CB |
| 8100F6 | 55 | 1200 | No | No | 0 | CB | CB | CB |
| 8100F6 | 55 | 300 | No | No | 11 | CB | CB | CB |
| 11090F6 | 11 | 6000 | yes | yes | 0 | CB | CB | CB |
| 11090F6 | 11 | 300 | yes | No | 52 | CB | 0.03 | CB |
| 30100F6 | 9 | 12000 | yes | No | 0 | 0.21 | CB | CB |
| 30100F6 | 9 | 12000 | NA | yes | 11 | CB | NA | CB |
| 30100F6 | 9 | 12000 | No | No | 26 | CB | CB | CB |
| 110100F6 | 140 | 600 | No | No | 0 | CB | CB | CB |
| 110100F6 | 140 | 600 | No | No | 15 | CB | CB | CB |

## References

Abbott, K.D., Ksiazek, T.G. & Mills, J.N., 1999. Long-term hantavirus persistence in rodent populations in central Arizona. Emerging infectious diseases, 5(1), pp.102–12. Available at: http://www.pubmedcentral.nih.gov/articlerender.fcgi?artid=2627700&tool=pmcentrez&rendertype=abstract.

Asano, B.M.S. & Ahmed, R., 1996. CD8 T Cell Memory in B Cell-deficient Mice. J. Exp. Med., 183(May).

Begon, M. et al., 2009. Effects of abundance on infection in natural populations: field voles and cowpox virus. Epidemics, 1(1), pp.35–46.

Billings, A.N. et al., 2010a. Pathology of Black Creek Canal Virus Infection. VECTOR-BORNE AND ZOONOTIC DISEASES, 10(6).

Billings, A.N. et al., 2010b. Pathology of Black Creek Canal virus infection in juvenile hispid cotton rats (Sigmodon hispidus). Vector borne and zoonotic diseases, 10(6), pp.621–628.

Borremans, B., 2014. Ammonium improves elution of fixed dried blood spots without affecting immunofluorescence assay quality. Tropical Medicine & International Health, 19(4), pp.413–416.

Borremans, B. et al., 2016. Estimating Time of Infection Using Prior Serological and Individual Information Can Greatly Improve Incidence Estimation of Human and Wildlife Infections. PLoS computational biology, 12(5), p.e1004882. Available at: http://www.ncbi.nlm.nih.gov/pubmed/27177244 [Accessed May 17, 2016].

Borremans, B., Sluydts, V., et al., 2015. Evaluation of short-, mid- and long-term effects of toe clipping on a wild rodent. CSIRO Wildlife Research, 42, pp.143–148.

Borremans, B. et al., 2011. Presence of Mopeia virus, an African arenavirus, related to biotope and individual rodent host characteristics: implications for virus transmission. Vector borne and zoonotic diseases (Larchmont, N.Y.), 11, pp.1125–31.

Borremans, B., Vossen, R., et al., 2015. Shedding dynamics of Morogoro virus, an African arenavirus closely related to Lassa virus, in its natural reservoir host Mastomys natalensis. Nature Publishing Group, pp.1–8.

Borucki, M.K. et al., 2000. Role of maternal antibody in natural infection of Peromyscus maniculatus with Sin Nombre virus. Journal of Virology, 74(5), pp.2426–2429.

Botten, J. et al., 2000. Experimental infection model for Sin Nombre hantavirus in the deer mouse (Peromyscus maniculatus). Proceedings of the National Academy of Sciences of the United States of America, 97(19), pp.10578–10583.

Burnet, F.M.; Fenner, F., 1949. The Production of Antibodies, 2nd ed, Melbourne, Australia,: Macmillan Magazines Ltd.

Chantrey, J., 1999. The epidemiology of cowpox in its reservoir hosts.

Childs, J.E. & Peters, C.J., 1993. Ecology and epidemiology of arenaviruses and their hosts. In M. S. Salvato, ed. The Arenaviridae. New York: Springer US, pp. 331–384.

Cooch, E.G. et al., 2012. Disease dynamics in wild populations: modeling and estimation: a review. Journal of Ornithology, 152(S2), pp.485–509. Available at: http://link.springer.com/10.1007/s10336-010-0636-3 [Accessed September 13, 2016].

Fisher-Hoch, S.P. & McCormick, J.B., 2001. Towards a human Lassa fever vaccine. Reviews In Medical Virology, 11, pp.331–341.

Fulhorst, C.F. et al., 2001. Experimental infection of Neotoma albigula (Muridae) with Whitewater Arroyo virus (Arenaviridae). The American journal of tropical medicine and hygiene, 65(2), pp.147–51.

Fulhorst, C.F. et al., 1999. Experimental infection of the cane mouse Zygodontomys brevicauda (family Muridae) with guanarito virus (Arenaviridae), the etiologic agent of Venezuelan hemorrhagic fever. Journal of Infectious Diseases, 180(4), p.966.

Fulhorst, C.F. et al., 2002. Experimental infection of the Sigmodon alstoni cotton rat with Caño Delgadito virus, a South American hantavirus. The American journal of tropical medicine and hygiene, 67(1), pp.107–11.

Gannon W.L., S.R.S., 2007. Guidelines of the American Society of Mammalogists for the use of wild mammals in research. JOurnal of Mammalogy, 88(3), pp.809–823.

Glass, G.E. et al., 1990. Determining Matrilines By Antibody-Response To Exotic Antigens. Journal of Mammalogy, 71(2), pp.129–138.

Goyens, J. et al., 2013. Density thresholds for Mopeia virus invasion and persistence in its host Mastomys natalensis. Journal of theoretical biology, 317, pp.55–61.

Günther, S., Hoofd, G., Charrel, R., Röser, C., Becker-Ziaja, B., Lloyd, G., Sabuni, C., Verhagen, R., van Der Groen, G., et al., 2009. Mopeia Virus-related Arenavirus in Natal Multimammate Mice, Morogoro, Tanzania. Emerging Infectious Diseases, 15(12), pp.6–10.

Günther, S., Hoofd, G., Charrel, R., Röser, C., Becker-Ziaja, B., Lloyd, G., Sabuni, C., Verhagen, R., van der Groen, G., et al., 2009. Mopeia virus-related arenavirus in natal multimammate mice, Morogoro, Tanzania. Emerging infectious diseases, 15(12), pp.2008–12. Available at: http://www.pubmedcentral.nih.gov/articlerender.fcgi?artid=3044542&tool=pmcentrez&rendertype=abstract [Accessed June 16, 2013].

Günther, S. & Lenz, O., 2004. Lassa virus. Critical Reviews In Clinical Laboratory Sciences, 41(4), pp.339390.

Hardestam, J. et al., 2008. Puumala Hantavirus Excretion Kinetics in Bank Voles (Myodes glareolus). Emerging Infectious Diseases, 14(8), pp.1209–1215.

Katakweba, A., Mulungu, L. & Eiseb, S., 2012. Prevalence of haemoparasites, leptospires and coccobacilli with potential for human infection in the blood of rodents and shrews from selected localities in Tanzania, Namibia and Swaziland. African Zoology, 47(April), pp.119–127.

Kearse, M., Moir, R., Wilson, A., Stones-Havas, S., Cheung, M., Sturrock, S., Buxton, S., Cooper, A., Markowitz, S., Duran, C., Thierer, T., Ashton, B., MeKearse, M., Moir, R., Wilson, A., Stones-Havas, S., Cheung, M., Sturrock, S., Buxton, S., Cooper, A., T. & r, T., Ashton, B., Mentjies, P., & Drummond, Antjies, P., & Drummond, A., 2012. Geneious Basic: an integrated and extendable desktop software platform for the organization and analysis of sequence data. Bioinformatics, 28(12), pp.1647–1649.

Leirs, H., 1994. Population ecology of Mastomys natalensis (Smith, 1834). Implications for rodent control in Africa A. Belgian Administration for Development Cooperation, ed., Brussels.

Lukashevich, I.S. et al., 1999. Lassa and Mopeia virus replication in human monocytes/macrophages and in endothelial cells: different effects on IL-8 and TNF-alpha gene expression. Journal of Medical Virology, 59, pp.552–560.

M. J. Buchmeier, R. M. Welsh, F. J. Dutko, A.M.B.A.O., 1978. The Virology a n d Immunobiology o f Lymphocytic C horiomen ing itis Virus Infection. ADVANCES IN IMMUNOLOGY,, 30, pp.275–331.

Marchandeau, S., Chaval, Y. & Goff, E. Le, 2000. … in the abundance of wild European rabbits Oryctolagus cuniculus and high immunity level over three years following the arrival of rabbit haemorrhagic disease Prolonged decline in the abundance of wild European rabbits Oryctolagus cuniculus and high immu. WILDLIFE BIOLOGY, 6(3), pp.141–147.

Martin, L.B., Weil, Z.M. & Nelson, R.J., 2008. Seasonal changes in vertebrate immune activity: mediation by physiological trade-offs. Philosophical transactions of the Royal Society of London. Series B, Biological sciences, 363(1490), pp.321–39. Available at: http://www.pubmedcentral.nih.gov/articlerender.fcgi?artid=2606753&tool=pmcentrez&rendertype=abstract [Accessed June 23, 2016].

Matloubian, M., Concepcion, R.J. & Ahmed, R., 1994. CD4 + T Cells Are Required To Sustain CD8 + Cytotoxic T-Cell Responses during Chronic Viral Infection. JOURNAL OF VIROLOGY,, 68(12), pp.8056–8063.

Milazzo, M.L. & Fulhorst, C.F., 2012. Duration of Catarina virus infection in the southern plains woodrat (Neotoma micropus). Vector borne and zoonotic diseases (Larchmont, N.Y.), 12(4), pp.321–4. Available at: http://www.pubmedcentral.nih.gov/articlerender.fcgi?artid=3319932&tool=pmcentrez&rendertype=abstract [Accessed September 13, 2016].

Mills, S.C. et al., 2010. Fitness trade-offs mediated by immunosuppression costs in a small mammal. Evolution; international journal of organic evolution, 64(1), pp.166–79. Available at: http://www.ncbi.nlm.nih.gov/pubmed/19686266 [Accessed May 10, 2016].

Murphy, F. A., Winn, W. C., Walker, D. H., Flemister, M. R. & Whitfield, S.G., 1976. Early lymphoreticular viral tropism and antigen persistence Tamiami virus infection in the cotton rat. Lab. Invest., 34, pp.125–140.

Mwanjabe, P.S., Sirima, F.B. & Lusingu, J., 2002. Crop losses due to outbreaks of Mastomys natalensis (Smith, 1834) Muridae, Rodentia, in the Lindi Region of Tanzania. International Biodeterioration & Biodegradation, 49(2-3), pp.133–137. Available at: http://linkinghub.elsevier.com/retrieve/pii/S0964830501001135.

Oldstone, M. B. A., 2002. Biology and pathogenesis of lymphocytic choriomeningitis virus infection A. Oldstone; Clarke, ed., Berlin Heidelberg: Springer-Verlag.

Oldstone, M.B.; Dixon, F.J., 1967. Lymphocytic choriomeningitis: Production of antibody by “tolerant” infected mice. Science, (158), pp.1193–1195.

Pollock, K.H. et al., 1990. Statistical Inference for Capture-Recapture Experiments. Wildlife Monographs, (107), pp.3–97.

R Core Team, 2014. R: A language and environment for statistical computing.

Rieger, T., Merkler, D. & Günther, S., 2013. Infection of type I interferon receptor-deficient mice with various old world arenaviruses: a model for studying virulence and host species barriers. PloS one, 8(8), p.e72290.

Sluydts, V. et al., 2007. Survival and maturation rates of the African rodent, Mastomys natalensis: density-dependence and rainfall. Integrative zoology, 2(4), pp.220–32. Available at: http://www.ncbi.nlm.nih.gov/pubmed/21396039 [Accessed June 21, 2013].

Telfer, S. et al., 2002. The effects of cowpox virus on survival in natural rodent populations: increases and decreases. Journal of Animal Ecology, 71, pp.558–568.

Tersago, K. et al., 2012. IMPACT OF PUUMALA VIRUS INFECTION ON MATURATION AND SURVIVAL IN BANK VOLES: A CAPTURE-MARK-RECAPTURE ANALYSIS IMPACT OF PUUMALA VIRUS INFECTION ON MATURATION AND SURVIVAL IN BANK VOLES: A CAPTURE-MARK-RECAPTURE. Journal of wildlife diseases, 48(1), pp.148–156.

Vitullo, A., Hodara, V. & Merani, M.S., 1987. Effect of persistent infection with Junin virus on growth and reproduction of its natural reservoir, Calomys musculinus. Am. J. Trop. Med. Hyg., 37, pp.663–669.

Voutilainen, L. et al., 2015. Life-Long Shedding of Puumala Hantavirus in Wild Bank Voles (Myodes glareolus). Journal of General Virology.

Walker, D.H. et al., 1975. Comparative pathology of Lassa virus infection in monkeys, guinea-pigs, and Mastomys natalensis. Bulletin of the World Health Organization, 52, pp.523–34.

Webb, P., Justines, G. & Johnson, K., 1975. Infection of wild and laboratory animals with Machupo and Latino viruses. Bulletin of the World Health Organization, 52, pp.493–9.

Weigand, H.; Hotchin, J., 1961. Studies of lymphocytic choriomeningitis in mice. J. Immunol, 86, p.401.

Wulff, H. et al., 1977. Isolation of an arenavirus closely related to Lassa virus from Mastomys natalensis in south-east Africa. Bulletin of the World Health Organization, 55, pp.441–444.

Yanagihara, R., Amyx, H.L. & Gajdusek, D.C., 1985. Experimental infection with Puumala virus, the etiologic agent of nephropathia epidemica, in bank voles (Clethrionomys glareolus). Journal of virology, 55(1), pp.34–8.

